# Poly(acrylamido) PEG-alternatives Enhance mRNA-LNP Efficacy in Immune Cells and Evade Anti-PEG Antibodies after Repeated Dosing

**DOI:** 10.64898/2026.02.26.708093

**Authors:** Benjamin M. Fiedler, Charlotte Galley, Margarita Strimaite, Nga Man Cheng, Najet Mahmoudi, Zhiping Feng, Tammy Kalber, Maria-Jose Martinez-Bravo, Chris Morris, Jenny K. W. Lam, Daniel J. Stuckey, Gareth R. Williams, Clare L. Bennett, Pratik Gurnani

## Abstract

Following the successes of the messenger RNA (mRNA) lipid nanoparticle (LNP) vaccines during the COVID-19 pandemic, mRNA-LNPs are being explored for many critical disease indications including infectious disease vaccination, cancer immunotherapies, and protein replacement therapies. LNPs require a polymer coating to provide stability in storage, and to minimise clearance from the body by reducing protein adsorption after injection. Poly(ethylene glycol)-lipids (PEG-lipids) have fulfilled this role to date, however increasing prevalence of antibodies against PEG in the general population jeopardises the efficacy of future PEGylated LNP doses and increases the likelihood of adverse pseudo-allergic responses. There is, therefore, an urgent unmet need to develop LNPs with new surfaces of PEG-alternative polymers which can evade anti-PEG antibodies, particularly where repeat dosing is required. Here, we present a family of polymer lipids, poly(acrylamido) (PAM) lipids, which effectively replace conventional PEG-lipids in mRNA-LNP formulations. We identify key design parameters to show that PAM-lipid monomer chemistry, molar mass and end-group all have critical effects on LNP size, polydispersity and *in vitro* transfection efficiencies, while having little impact on LNP morphology or internal structure. We determine that side-group (monomer) chemistry is a key mediator in alleviating anti-polymer antibody cross-reactivity. Compared to clinical benchmark PEGylated LNPs, several PAM-LNPs displayed improved transfection efficacy across multiple mRNA cargos in diverse cell types, organs, and routes of administration, both *in vitro* and *in vivo*. In particular, mRNA transfection improved in immune cells both *in vitro* (up-to 120-fold), and *in vivo* (up-to 5-fold), including superior mRNA expression in lymph nodes (2.5-fold). In part, this is likely because PAMs increase LNP uptake/association with primary immune cells (BMDCs), and increase biodistribution to the lymphoid tissues (LNs, spleen). Crucially, PAM-LNPs avoid circulating anti-PEG antibodies to recover lost mRNA efficacy after repeated dosing *in vivo*, 300% higher than PEG-LNPs. Overall, our findings establish the PAM-lipid family as a versatile platform of chemically varied PEG-alternatives, towards the next generation of therapeutic mRNA-LNP technologies.

**Graphical Abstract:** 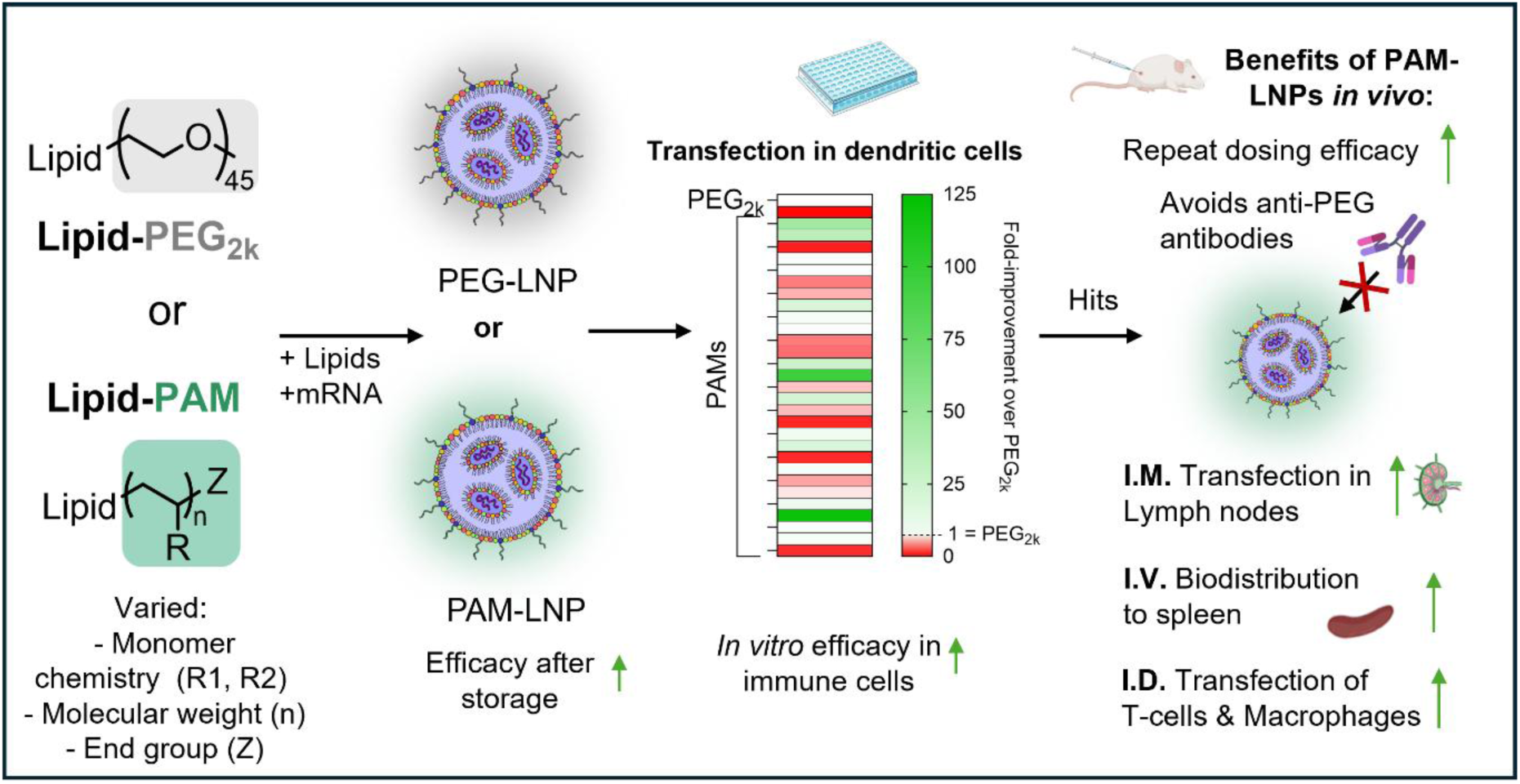

## Introduction

With the successful rollout of mRNA vaccines against COVID-19 (Comirnaty, Spikevax, Zapomeran and Gemcovac), there has been increased development of mRNA lipid nanoparticle (LNP) technologies for a vast array of critical indications including cancer, cardiovascular disease and genetic disorders^1–3^. Currently there are over 150 clinical trials for LNP-mediated gene delivery^2^. LNPs are a crucial part of this technology as they protect RNA from degradation by nucleases and enhance cellular uptake/trafficking. Conventional LNPs are composed of four different components: an ionizable (cationic) lipid, cholesterol, a helper phospholipid, and a polymer-lipid conjugate. This multicomponent formulation allows control over the physical properties of the LNP, which in turn impact biological activity such as mRNA transfection efficiency, tissue tropism and cell targeting. All clinically approved LNPs, and most under investigation, utilize poly(ethylene glycol) (PEG)-lipids within their formulation to control particle size, aid storage stability, increase circulatory half-life, and reduce protein adsorption to the LNP surface^1^. However the ubiquity of PEG in other drug and cosmetic formulations meant that ∼72% of human blood samples tested positive for anti-PEG antibodies before the COVID-19 pandemic (2016)^4^. This increased up-to 69-fold after the use of PEG in immunogenic COVID-19 vaccines^5^. These pre-existing anti-PEG antibodies reduce the efficacy of future doses of PEGylated therapies in humans^6^. This is partly through the accelerated blood clearance effect (ABC), where opsonisation by pre-existing anti-PEG antibodies increases clearance of LNPs by liver resident macrophages (Kupffer cells)^7–10^. In addition to impacting intracellular trafficking (e.g. cellular uptake), opsonisation can destabilise LNPs causing premature cargo leakage^8,11^. By inducing structural instability and promoting immunogenicity, anti-PEG antibodies directly counteract the intended stealth and stabilizing properties of PEGylation in LNP formulations. It is therefore likely that future mRNA therapies may exhibit poor treatment outcomes in patients with pre-existing anti-PEG antibodies, for example after COVID-19 mRNA-LNP vaccination, if PEGylated LNPs (PEG-LNPs) are used. It has been previously hypothesised, but not demonstrated, that it may be possible to recover this lost efficacy by dosing LNPs with a PEG-alternative surface, evading opsonisation from anti-PEG antibodies since these are unable to bind to PEG-alternatives with different chemical structures^12^. Furthermore, sensitisation to PEG has been associated with adverse pseudo-allergic responses (complement-activation-related pseudo allergy, CARPA), with these events rare but increasing in prevalence^13^. The overuse of PEG in mRNA-LNP vaccine formulations urgently necessitates the development of new PEG alternatives which convey the benefits of PEG while avoiding neutralising anti-PEG antibodies.^45,47,48^

Existing work into the discovery of PEG alternatives for mRNA-LNPs has often followed a “one-by-one” approach. While this has yielded promising options such as poly(sarcosine) and poly(oxazoline), these polymers still stimulate an adverse anti-polymer immune response, meaning individually they will not solve the general problem of anti-polymer immunogenicity^12,14–16^. Akin to antibiotics loosing efficacy when overused, a wide selection of PEG alternatives is likely needed to secure the future efficacy of mRNA-LNP therapies. Therefore, a broader approach to the discovery of PEG alternatives must be taken.

To address this, we have developed a family of effective replacements for PEG-lipids in LNP formulations: N-substituted poly(acrylamido) lipids (PAM-lipids). The synthetic control offered by synthesising PAMs via reversible-addition fragmentation chain transfer (RAFT) polymerisation allowed us to generate a library of 30 chemically varied PAM-lipids, all conjugated to the dimyristoyl glycerol (DMG) lipid, as DMG-PEG conjugates are used in FDA-approved LNP formulations. By using this library to replace PEG-lipids in Moderna’s clinical LNP formulation (SpikeVax™) we explored the key chemical design features of PAMs that result in mRNA-LNPs with favourable small particle sizes (∼100 nm) and high transfection efficiencies *in vitro* compared to PEGylated LNPs. As immune cells such as dendritic cells are key targets for >80% of RNA-LNP clinical trials^2^, we demonstrate that PAMs are superior alternatives to PEG in terms of mRNA-LNP transfection efficiency of immortalised and primary immune cells *in vitro* (up to 120-fold) and after 3 weeks of storage. We have characterised the nanoparticle structure of PAM-LNPs, and investigated their use across different common routes of administration (IM, ID, IV), with multiple PAM-LNPs showing effective transfection *in vivo*. In particular, DMA-based PAM-LNPs significantly enhanced mRNA expression within lymph nodes (2.5-fold), and increased *in vivo* transfection of immune cells (macrophages, and T-cells, ∼5-fold) compared to PEG-LNPs. Crucially, we demonstrate that PAM-lipids avoid anti-PEG antibodies to recover the lost transfection efficiency caused by repeated dosing of PEG-LNPs. Finally, we discover that PAM monomer chemistry, rather than end group, is the key structural feature for polymers to effectively evade anti-polymer antibody binding. Ultimately, these data establish PAM-lipids as a family of effective replacements for PEG-lipids in mRNA-LNP technologies.

## Results and Discussion

### PAM-lipids with -H or -OH end group enhance efficacy of mRNA-LNPs compared to PEG-lipids

To identify novel and effective PEG alternatives, we synthesised a library of poly(acrylamido) lipids (PAM-lipids) *via* reversible-addition fragmentation chain-transfer (RAFT) polymerisation (Figure 1A, S4). This allowed us to produce a chemically diverse library of PAM-lipids, with highly controlled polymer chemistry, molecular weights and end-groups, in agreement with results reported for other systems in previous literature reports^17^. These parameters were hypothesised to be critical design parameters for LNP function. RAFT polymerisation was mediated by a synthesised dimyristoyl glycerol (DMG) functionalised chain transfer agent (CTA) (Schematic S1-4) to impart a DMG lipid tail end-group to the polymer, which enables integration into the LNP bilayer. Five substituted acrylamide monomers with different chemical properties (e.g. functional groups, steric bulk) were chosen due their hydrophilic nature, which is a key property of PEG in stabilising LNPs. These monomers were used to prepare polymers of two chain lengths (degree of polymerisation, DP = 20 or 60). These specifications were chosen to roughly match the molar mass (DP20, ∼2000 kDa), fully extended length (DP60, ∼120 atoms long) and lipid chain end (DMG) of PEGylated lipid DMG-PEG_2k_ used in Moderna’s SpikeVax® formulation ^1,18–20^. The -S_3_C_5_H_9_ trithiocarbonate end-group of all PAM-lipids was then modified to either -OH or -H *via* oxidation or hydrogen radical donation respectively, resulting in a library of 30 unique PAM-lipids each with one of: five monomers, two chain lengths, three end groups. ^1^H NMR spectroscopy confirmed high monomer conversion (>95%), that the expected structure was synthesised, and that any residual monomer was removed during purification (table S1). Gel permeation chromatography (GPC) confirmed efficient RAFT-mediated control of molecular weight and dispersity from using the DMG-CTA, with PAM-lipids displaying low dispersities (mean dispersity = 1.16 ± 0.10). GPC-coupled UV/Vis detection at 309 nm verified the presence, and subsequent removal, of the trithiocarbonyl end group, corresponding to polymers with the –S₃C₅H₉ or -OH/-H end groups, respectively (Table S1).

**Figure 1.**
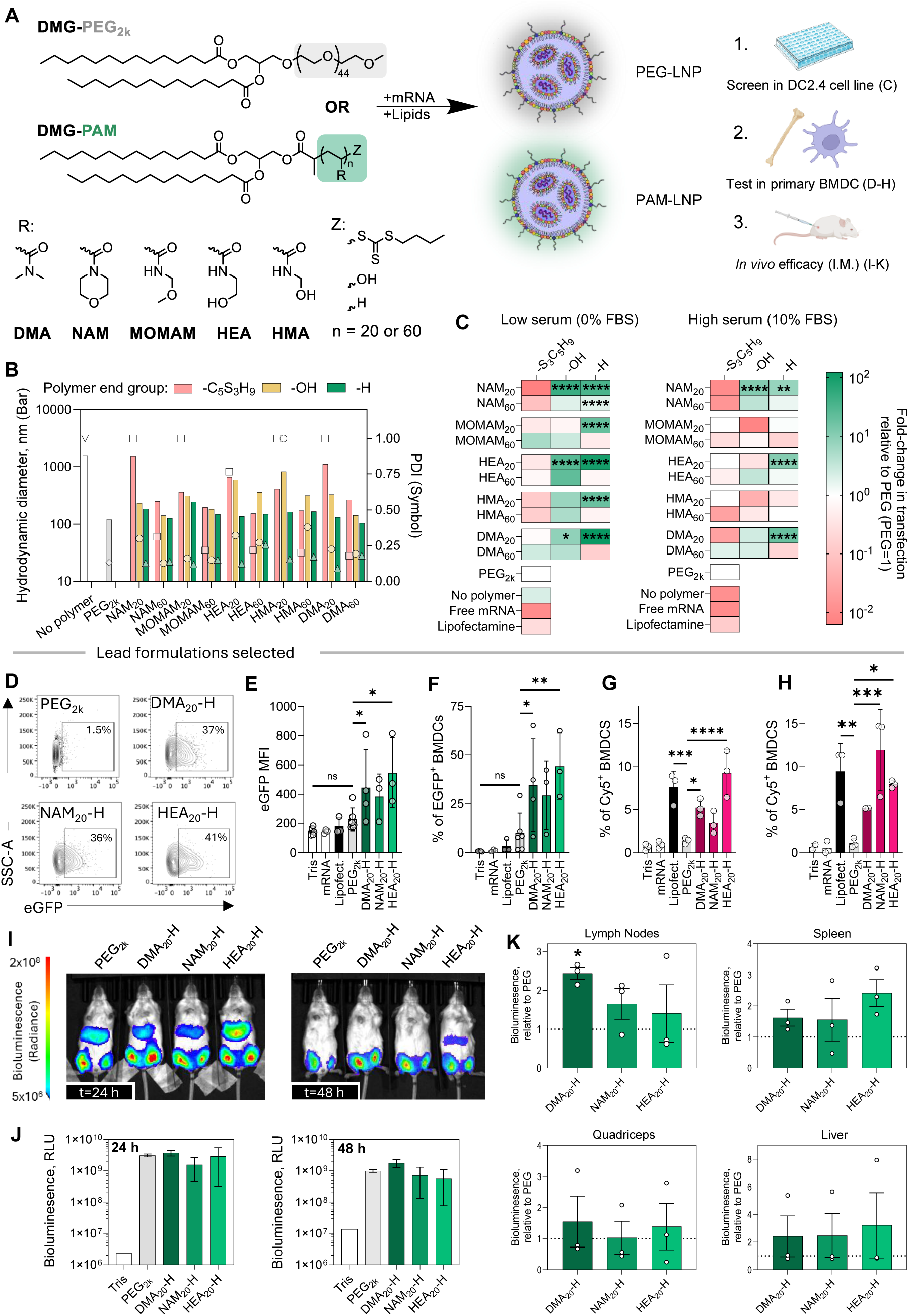
PAM-lipid chemistry modulates LNP size and mRNA delivery performance *in vitro* and *in vivo*. **A** Formulation of PAM-LNPs from synthesised DMG-based PAM-lipids with varied chemistries (5 monomers, 3 end-groups and 2 molar masses) and control PEG-LNPs, and the following testing cascades. R= monomer/polymer side group; Z=end group; n=target number of repeating units **B** Z-average hydrodynamic particle diameter (intensity) and polydispersity index of LNPs in Tris (50 mM, pH 7.4) at 25°C from dynamic light scattering. **C** Firefly luciferase activity (relative to control PEG-LNP) in DC2.4 cells 24 hours after treatment with PAM-LNPs loaded with mRNA encoding firefly luciferase in OptiMEM, with or without supplemental foetal bovine serum (10% w/v). Data are presented as mean of *n =* 3 technical replicates, n = 3 biological replicates, statistical significance was determined by one-way ANOVA against the control PEG-LNPs. \**P* < 0.05, \*\**P* < 0.01, \*\*\**P* < 0.001, \*\*\*\**P* < 0.0001. Quantification of eGFP expression in BMDCs treated with mRNA-LNPs encoding eGFP by flow cytometry via: **D** representative pseudoplots; **E** median fluorescence intensity; and **F** percentage of eGFP^+^ cells. Quantification of Cy5^+^ BMDCs treated with LNPs containtingCy5 tagged eGFP-mRNA via flow cytometry in **G** high serum and **H** low serum conditions. Data are presented as mean ± s.e.m. of biological replicates, statistical significance was determined by mixed effects model with Dunnett correction compared against the control PEG-LNPs. \**P* < 0.05, \*\**P* < 0.01. *In vivo* evaluation of fLuc mRNA (2.5 µµg of mRNA per quadricep, 2 injections per mouse) expression after delivery by PAM-LNPs following IM administration represented as absolute whole-body bioluminescence at 24 h and 48 h post administration (**I, J**) and as individual tissues excised after 48 hours (**K**) relative to PEG-LNPs. Quadriceps and lymph node data points were summed from each quadricep and corresponding draining lymph node for each animal respectively (*n*=2 lymph nodes per animal). Data are presented as mean ± s.e.m. of biological replicates (n = 4 for PEG_2k_; n = 3 for DMA_20_-H, NAM_20_-H, and HEA_20_-H; n=1 for Tris control), statistical significance was determined by Welch one-way ANOVA against the control PEG-LNPs, with Dunnett Correction. \**P* < 0.05.

PAM-lipids were then used to prepare LNPs (PAM-LNPs) by directly replacing the DMG-PEG_2k_ in Moderna’s commercial SpikeVax® formulation (Figure 1A). Otherwise, the formulation was kept the same regardless of polymer used, comprising; ionizable lipid SM102, DPSC, cholesterol, and either DMG-PEG or a DMG-PAM at a 50:10:38.5:1.5 molar ratio, respectively. An organic solution of lipids was rapidly mixed with mRNA dissolved in an acidic aqueous buffer to produce PEG-LNPs or PAM-LNPs at a nitrogen-to-phosphate ratio (N/P) of 6, following established protocols^21^. Particle size measurements revealed that over half of formulations containing PAM-lipids produced LNPs below 200 nm diameter and PDI < 0.25, similar to the PEG-2k LNP (∼120 nm) (Figure 1B, 3D, 4A). The majority of LNPs derived from DMG-PAMs that retained the DMG-CTA’s native hydrophobic trithiocarbonate end-group (-C_5_S_3_H_9_) exhibited larger particle diameters (> 200 nm) and higher PDI values (> 0.25) than when this end group was replaced with -OH or -H (Figure 1B). In contrast, PAM-LNPs with - OH and -H end groups predominantly yielded smaller and more uniform LNPs, indicating the benefit of non-hydrophobic end-groups to stabilise LNP formation. mRNA encapsulation efficiency was >90% for all PAM-LNPs and PEG-LNPs (Figure S1).

The efficacy of mRNA encapsulated within PAM-LNPs was first screened in a dendritic cell line (DC2.4) as dendritic cells (DCs) are a crucial target for key therapeutic applications of mRNA-LNPs (Figure 1C)^2^. PAM- and PEG-LNPs were incubated with cells in high and low serum conditions (OptiMEM, +/- 10% FBS w/v, respectively) for 24 hours. Figure 1C demonstrates that 9/30 PAM-LNPs significantly outperformed the control PEG-LNP transfection efficiency in either low serum (up to 120-fold) or high serum media (up to 20-fold). Inclusion of serum in *in vitro* studies of mRNA-LNPs is often overlooked, but improves the physiological relevance of the assay given that PEG’s role in controlling protein adsorption to the LNP surface when exposed to serum after injection^22^. Which serum proteins adsorb to nanoparticle surfaces can change between particles with different polymer surface chemistries^23–25^, which in turn can impact LNP uptake/endosomal escape^26^, inflammatory response^27^, and organ biodistribution^28^. Moreover, serum can abstract PEG-lipids, modulating LNP uptake, endosomal escape and mRNA efficacy^27–33^. Serum proteins such as apolipoprotein E can also restructure LNPs and cause cargo leakage^34^. Unsurprisingly then, LNPs prepared without a PEG-lipid or PAM-lipid transfected ∼100-fold less effectively than PEG-LNPs in high serum conditions, despite transfecting well in low serum media (Figure 1C). Broadly, the highest transfection efficiencies were observed for LNPs prepared with short ∼DP20 PAM-lipids bearing either hydrophilic -OH or -H end groups, while near-complete loss in activity was regularly seen when PAMs with the more hydrophobic -S_3_C_5_H_9_ end group were used. This is not due to any noticeable toxicity (Figure S2A), but instead large particle size may reduce the efficiency of endocytic uptake^35^. This end-group effect on mRNA efficacy was most pronounced for short DP20 polymers (figure S3), where end group represents a greater portion of the polymer as a whole. Generally, the -H end group was best for DP20 polymers, while -OH was best for DP60 polymers. These data highlight how surface polymer chemistry can greatly modulate LNP efficacy. In particular, the presence/absence of small hydrophobic end groups are a disproportionately impactful parameter for the design of polymers for mRNA-LNPs, considering their small size relative to that of the polymer chain (∼5% of polymer molecular weight), which makes up only ∼6% of the whole LNP formulation mass. As such, the -S_3_C_5_H_9_ end group is only ∼0.3% of the total mRNA-LNP formulation by weight, but can modulate efficacy across 4 orders of magnitude. To the best of our knowledge, this has not yet been reported as a key design feature of PEG-alternative polymers.

To determine whether PAM-LNPs are also superior at transfecting primary DCs, we used three high-performing PAMs (DMA_20_-H, HEA_20_-H and NAM_20_-H) from the screen in DC2.4s to prepare LNPs loaded with eGFP-encoding mRNA. Bone marrow dendritic cells (BMDCs) were derived with Flt3L from mice (C57BL/6), incubated with PAM-LNPs or PEG-LNPs and analysed *via* flow cytometry (gating strategy demonstrated in SI, Section 1.3). All three PAM-LNPs effectively transfected BMDCs, with DMA_20_-H and HEA_20_-H LNPs both significantly outperforming the benchmark PEG-LNPs, increasing the percentage of transfected cells to 34% and 44% respectively compared to 10% with PEG. Transfection using fluorescently tagged mRNA (Cy5-mRNA) revealed increased association of PAM-LNP (∼4 - 10x increase) by BMDCs compared to PEG-LNPs (Figure 1D-F, Figure S1A-G) in both high and low serum media. This is likely due to increased cellular uptake and or/retention of LNPs (e.g. *via* increased endosomal escape), which could mechanistically explain how modulating LNP surfaces with PAMs increases mRNA-LNP transfection in BMDCs compared to PEG. Replacing PEG with these lead PAMs formulations did not impact cellular viability (Figure S2A) or activation (CD86 expression, Figure S4D) of BMDCs.

We then investigated if the superiority of PAM-LNPs at mRNA transfection *in vitro* translated to enhanced mRNA expression *in vivo*, and whether PAM-LNPs with different surface chemistries had different tissue tropisms. We injected the same three high-performing PAM-LNP formulations prepared with mRNA encoding firefly luciferase intramuscularly (IM) into BALB/c mice, as IM is a common route of administration for vaccines/immunotherapy^36,37^. Bioluminescence location and intensity were compared to a PEG-LNP control. Bioluminescence imaging demonstrated PAM-LNPs effectively transfect FLuc-mRNA *in vivo* 24- and 48-hours post-injection, with most expression visualised at the site of injection (muscle) and in the liver for all formulations. Direct measurement of excised tissues after 48 hours revealed that DMA_20_-H LNPs showed a significant 2.5-fold increase in mRNA expression in lymph nodes (LNs) relative to PEG-LNPs. We suggest that by screening PAM-LNPs in physiologically relevant media and cell types (high serum, DC2.4s), and confirming their efficacy in BMDCs, we have selected PAM-LNP formulations with a high affinity for tissues rich in immune cells. mRNA expression was otherwise comparable between PEG-LNPs and PAM-LNPs in the quadriceps and liver. Together these data demonstrate that replacement of PEG-lipids with PAM-lipids in mRNA-LNP formulations results in enhanced transfection of dendritic cells *in vitro* and antigen expression in lymphoid organs *in vivo*, particularly after selecting optimal polymer chain length (DP20) and end group (-H).

In particular, LNPs prepared with DMA_20_-H displayed more potent performance than PEG in primary dendritic cells and immune rich tissues, and are therefore promising for vaccine and immunotherapy applications. Despite only representing 1.5 mol% of the formulation, polymer-lipids (and in particular their end groups) are furthest away from the LNP surface and thus likely to interact first with biological entities and strongly influence potency. Promisingly, PAMs prepared from 3 of the 5 monomers investigated (DMA, NAM, HEA) can be used to replace PEG without compromising particle size, *in vitro* or *in vivo* performance, highlighting PAMs as a family of useful polymers to replace PEG in LNP-mediated mRNA delivery.

Having established the efficacy of PAM-lipids for mRNA-LNPs *in vitro* and *in vivo*, we sought to investigate: (i) if PAM-LNPs improve transfection in immune cells *in vivo*; (ii) if the improved efficacy by using PAM-lipids over PEG-lipids can be attributed to changes in LNP structure or stabilisation; and (iii) if PAM-lipids improve transfection in populations with high anti-PEG antibodies during repeat dosing regimens. We address each of these in the following 3 sections.

### PAM LNPs enhance mRNA expression in lymph node immune cells over PEG2k LNPs

Having demonstrated that PAMs enhanced mRNA transfection efficacy of mRNA-LNPs in immune cells *in vitro* and in lymphoid organs *in vivo* compared to clinical gold-standard PEG-LNPs, we sought to investigate transfection efficiency at a single cell resolution *in vivo*. Mice (C57BL/6) were administered intradermally (ID)^38,39^ with LNPs prepared with PAM-lipid DMG-DMA_20_-H (DMA-LNPs) or DMG-PEG_2000_ (PEG-LNPs) containing mRNA encoding enhanced green fluorescent protein (eGFP) as a marker for positive transfection (Figure 2A). Skin (site of injection), draining lymph nodes (vicinal), spleen and liver were collected and analysed *via* flow cytometry. DMA-LNPs induced a significantly stronger transfection signal in transfected immune cells (CD45^+^) than PEG-LNPs across both timepoints combined (Figure 2B). This aligns with the increased lymph node transfection caused by DMA-LNPs compared to PEG-LNPs after IM injection (from Figure 1I-K), which is particularly promising as it suggests than the lymphatic targeting of DMA-LNPs is independent of mRNA cargo, route of administration, or animal strain. As with IM administration, PAM-LNPs also transfected the liver and spleen comparably to PEG-LNPs, while cell numbers isolated from skin were too low for robust analysis.

**Figure 2.**
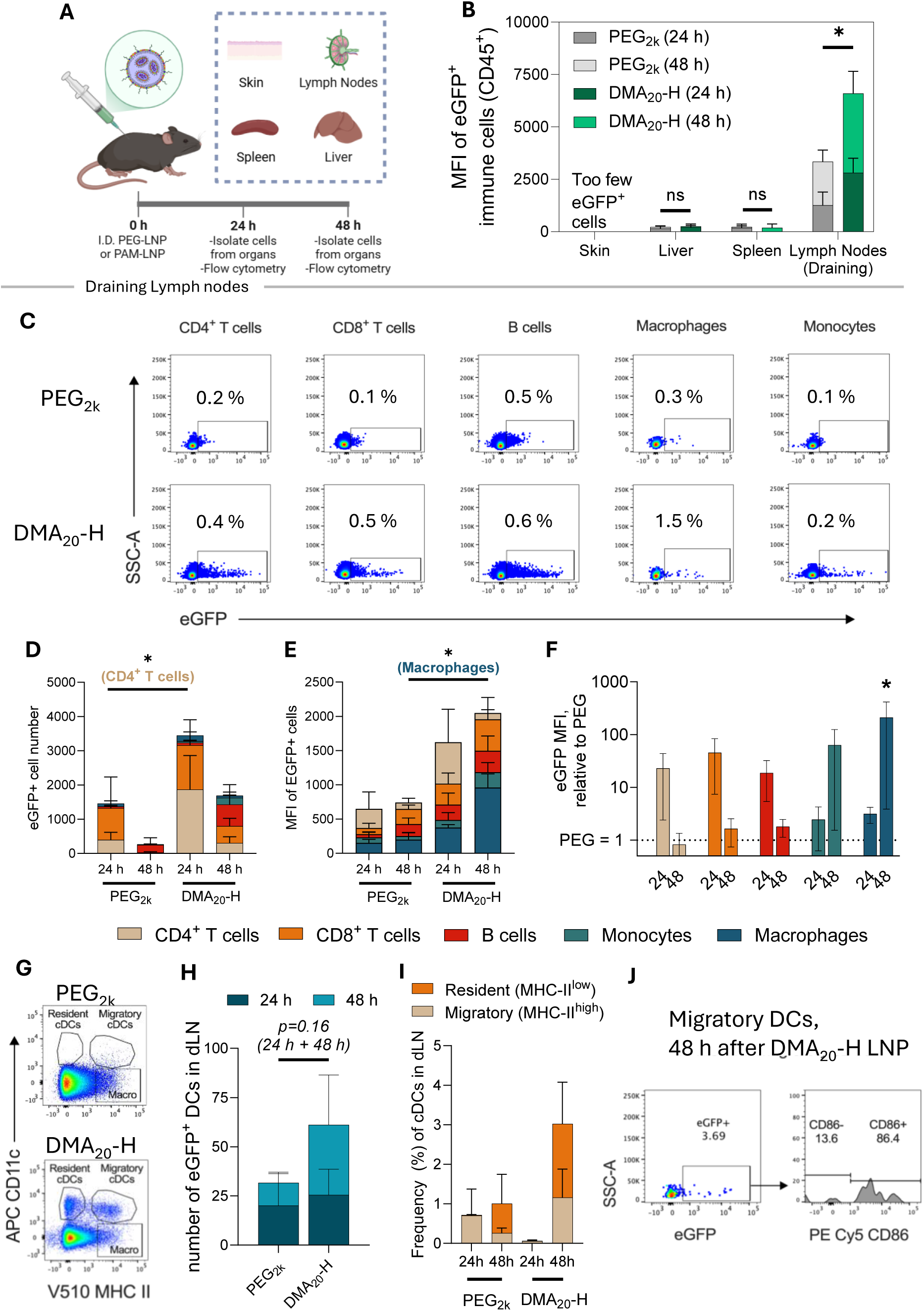
DMA-LNPs demonstrate broad-spectrum improvement in transfection of immune cells relative to PEG-LNPs in draining lymph nodes. **A.** Schematic showing experiment where C57B/L6 mice were injected ID with eGFP mRNA-LNPs, formulated with PEG-lipids or the PAM-lipid DMA_20_-H, or Tris buffer control. Organs (skin, liver, spleen and dLN) were harvested for analysis *via* flow cytometry **B.** Stacked bar chart of eGFP MFI of CD45^+^eGFP^+^ cells (above background) at 24 and 48 hours. Data are presented as mean ± s.e.m., n = 3 biological repeats. Statistical significance was determined by two-tailed paired t-test of total values for 24 and 48 h, *P < 0.05 **C.** Representative flow plots showing eGFP expression in gated immune cell populations in draining LNs (see supplemental Figure S12 for gating strategy). **D.** Stacked bar graph showing eGFP+ cell numbers in dLN at 24 and 48 hours **E.** Stacked bar graph showing eGFP MFI of transfected cells (eGFP+) in dLN in each cell type over 24 and 48 hours. **F**. Bar graph showing the eGFP MFI of cells in dLN for each cell type over 24 and 48 hours, relative to PEG. **G.** Representative flow plots of MCH II and CD11c expression in myeloid cells (Live, single, CD45+, Lin- (CD19, B220, CD3e, Ly6c, ly6g)) at 48 hours in the dLN. **H.**Stacked bar graph showing the frequency of cDCs in dLN over 24 and 48 hours. **I.** Stacked bar graph showing the number of eGFP+DCs in dLN. **J.** Left, representative flow plot of eGFP expression in migratory cDCs (Live, single, CD45+, Lin- (CD19, B220, CD3e, Ly6c, ly6g) MHC II^high^ CD11c+) in dLN at 48 hours. Right, representative histogram of CD86 expression in eGFP+ migratory cDCs (Live, single, CD45+, Lin-(CD19, B220, CD3e, Ly6c, ly6g) MHC II^high^ CD11c+) in dLN at 48 hours. For **D-J** data are presented as mean (above control) ± s.e.m. of n = 3 biological repeats. Statistical significance was determined by two-way ANOVA with Tukey correction (**D-F**) or two-tailed unpaired T-test (**H**), *P < 0.05

Immunophenotyping of cell populations in the draining lymph nodes (dLN) show that replacing PEG with the PAM-lipid DMA_20_-H gives broad spectrum increases in the percentage of transfected cells across different immune cell types (**Figure 2C-D**). After 24 hours, DMA-LNPs transfected significantly more CD4^+^ T cells (4-fold increase) than PEG-LNPs, and also showed a non-significant 40-fold increase in transfected CD8+ T cells at 48 hours. This could directly improve vaccination outcomes, as transfection of CD4^+^ T-cells has recently been shown to be a functional target for generating adaptive immunity to viral infections (COVID-19) by mRNA-LNPs^40^. Moreover, the potential application of DMA-LNPs to generate T cells *in situ* for CAR-T cell therapy offers a strategy to bypass the current requirement for T cell extraction and *ex vivo* modification^41–43^.

Furthermore, after 48 hours, DMA-LNPs significantly increased transfection in macrophages, evidenced by higher overall eGFP MFI relative to PEG-LNPs, and higher MFI within the eGFP⁺ population (Figure 2F and 2E respectively). Macrophages in the dLN have been shown to be a key target for mRNA-LNP cancer immunotherapies^44^. Increased transfection of macrophages could benefit various other therapeutic mRNA targets including those related to wound healing and nerve regeneration^45,46^.

Finally, a key aim for vaccine design is targeting and activation of DCs to prime the naive T cell response. We observed that PAM-LNPs effectively transfect DCs in the dLN, with a non-significant 2-fold increase in number of transfected DCs over PEG-LNPs (p=0.16) (Figure 2H). This could be due to either DMA-LNPs draining to the dLNs themselves over time, or DCs transfected elsewhere (e.g. the site of injection) draining to the dLN with time. An increase in percentage of both migrating and resident cDCs (but not CD11c+ MHCII^-^ macrophages) at the 48 hour timepoint suggests both of these mechanisms may be occurring simultaneously. Transfection efficiencies by DMA-LNPs of ∼3 % are high compared to PEGylated LNPs in this study and previously reported in the literature (0 - 1.5 %)^47^. Most of the transfected migrating cDCs are effectively activated (86% positive for CD86), suggesting they are effectively poised to prime T cells (Figure 2J).

Taken together, these data demonstrate PAM-LNP formulations prepared with DMA_20_-H outperform PEG-LNPs at transfecting immune cells in the dLN after ID administration. Notably, DMA-LNPs significantly improve transfection in CD4^+^ T-cells and macrophages, and effectively recruit of cDCs to the dLN, highlighting their potential for use across a variety of immune-cell targeted therapeutic applications.

### PAM-lipids form similar LNP structure to PEG-lipids with improved efficacy before/after storage

Having characterised the biological efficacy of PAM-lipids and identified DMA_20_-H, NAM_20_-H and HEA_20_-H as effective replacements for PEG_2k_ in mRNA-LNP formulations, we then sought to establish how polymer-lipid chemistry affects LNP size, morphology, internal structure and stability. Any changes in these physicochemical properties could explain PAMs ability to modulate the efficacy of encapsulated mRNA *in vitro* and *in vivo*. PEG-lipids have a crucial role in stabilising the LNP structure during the formulation process by inserting into the surface of the nanoparticles after self-assembly and preventing excessive particle agglomeration. LNPs prepared with PEG_2k_, DMA_20_-H, NAM_20_-H, or HEA_20_-H were analysed via cryo-electron microscopy (cryoEM), small-angle light scattering (SANS), small-angle X-ray scattering (SAXS), and dynamic light scattering (DLS) as orthogonal techniques to characterise particle size, morphology, internal structure and heterogeneity (Figure 3).

**Figure 3.**
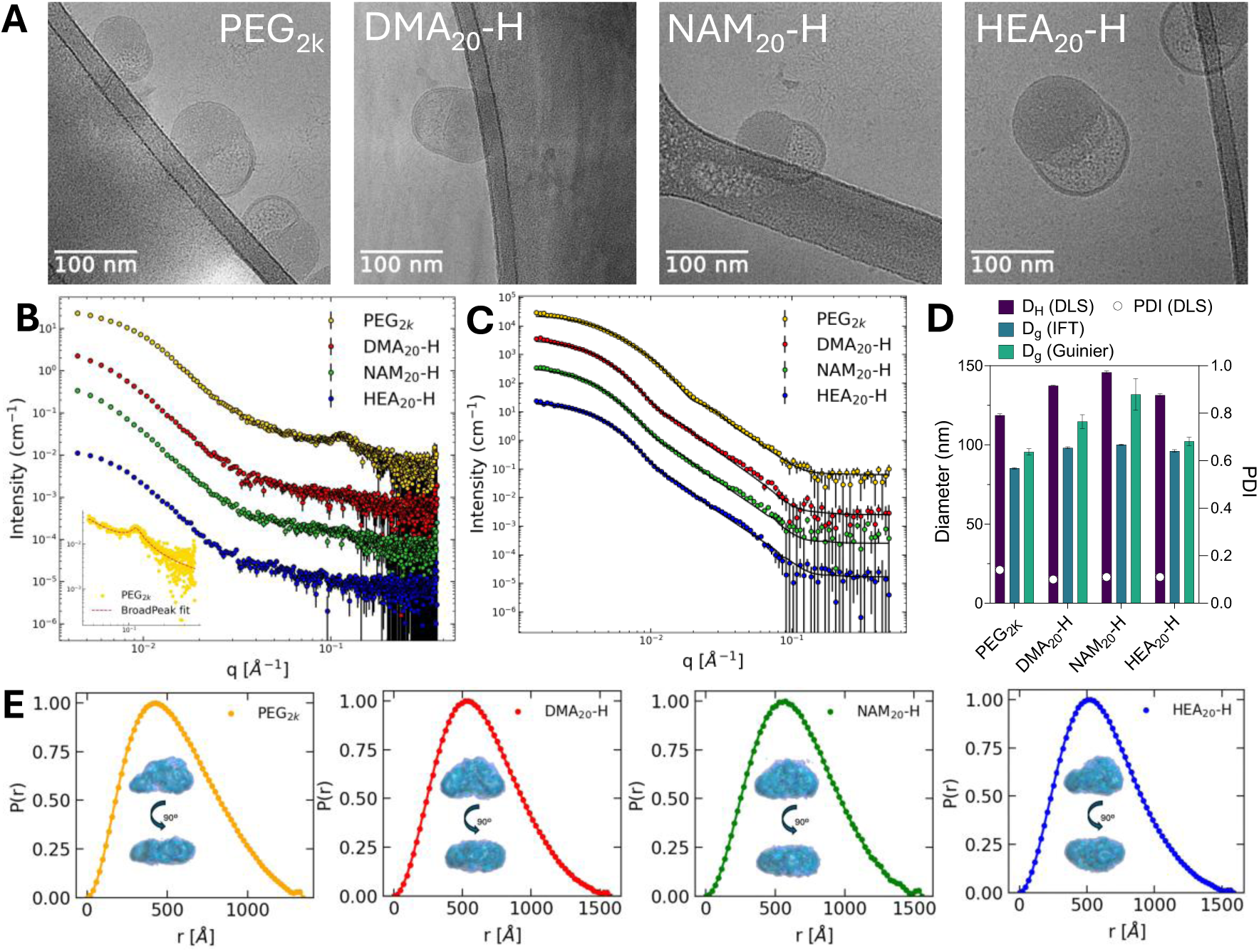
PAM-lipids and PEG-lipids produce LNPs with similar internal structure and morphology. Cryo-electron microscopy images (**A**) of LNPs prepared with PEG-lipids or PAM-lipids, loaded with eGFP-mRNA. (**B**) SAXS (**B**) and SANS (**C**) profiles of LNPs prepared with PEG-lipids or PAM-lipids loaded with polyA, with the inset of **B** showing a broad peak fit of the high-q PEG_2k_ experimental data. The solid lines are fits using a core-two-shell cylinder model. (**D)** Hydrodynamic diameter (Bar, left Y axis) and polydispersity (PDI, symbol right axis) of LNPs prepared with PEG-lipids or PAM-lipids, loaded with eGFP-mRNA, analysed with dynamic light scattering (DLS). Data presented as mean ± s.d. n = 3 and compared to the diameter of gyration from the Guinier analysis (table S2) and the *P*(*r*) evaluation. (E) *P*(*r*) analyses and DENSS ab initio reconstructions (insets) of SANS data.

DLS revealed that the nanoparticle size (hydrodynamic diameter, D_H_) and polydispersity were 100-150 nm and ∼0.1, respectively, regardless of the polymer-lipid used (Figure 1B, 3D, 4A), with particle sizes (Diameter of Gyration, D_g_) determined via SANS analysis similar in magnitude (Figure 3D). CryoEM images revealed slightly elongated structures with phase separation between the internal aqueous and lipid-rich compartments (“blebbing”) observed in all formulations (Figure 3A), consistent with the bleb structures observed for PEGylated mRNA-LNPs in other studies. Blebs have been suggested as crucial for effective mRNA transfection^48^. SAXS analysis of the internal structure revealed a small Bragg peak at q=0.11 Å^-1^ in LNPs prepared with PEG_2k_, corresponding to an internal mesophase of ∼5.5. nm, consistent with other PEGylated mRNA-LNPs (Figure 3B)^49,50^. Interestingly this peak was not well resolved in the PAM-LNPs, suggesting a less ordered internal structure. Decreases in this peak, but not complete disappearance, has been seen in other reports of LNPs with other PEG-alternatives that also improved mRNA efficacy ^50^.This elongated, anisotropic structure observed in cryoEM was confirmed with SANS *P*(*r*) pair distance distribution function analysis and *ab initio* three-dimensional structures (DENSS; insets in figure 3E, Table S1))^49,51,52^ and is consistent with the core-shell bleb morphology seen in cryoEM (Figure 3A). Model-based analysis (cylindrical core-two-shell model, Figure 3C; and core-shell ellipsoid model, Figure S5) implies that the representative LNP shape is elongated with long and short dimensions respectively varying from 112 nm and 26 nm for PEG to 138 and 50 nm for NAM (table S4). While some report spherical mRNA-LNPs, they are typically prepared with different ionizable lipids to SM-102, such as MC3 (D-Ln-Mc3-DMA) ^34,53^. Ultimately, PAM-lipids form LNPs of similar size and morphology to PEG-lipids, potentially with small differences in internal organisation.

Alongside stablishing LNP structure during the formulation process, one of the key properties that PEGylation bestows on nanoparticles is increased storage stability^54^. The polymer shell acts as a steric barrier to minimise aggregation which, over time, can change nanoparticle structure and negatively impact biological efficacy, since the efficacy of clathrin-mediated endocytosis begins diminishing for particles above ∼100 nm^35^. Effective PEG-alternatives must therefore fulfil this stabilising role. The medium- to long-term instability at fridge temperature of both Moderna and Pfizer/BioNTech mRNA COVID-19 vaccines also posed a significant issue during the early phase of the recent pandemic, requiring these products to be frozen below -20°C or -80°C for extended periods^55^. Hence, innovations to reduce the cold-chain requirement for mRNA LNPs are critical to minimising the cost and expanding the scope of mRNA medicines. We therefore compared the stability of the three high-performing PAM-lipids, DMA_20_-H, NAM_20_-H and HEA_20_-H against the benchmark PEG_2k_ in this context. Figure 4A shows that PAM-LNPs and PEG-LNPs had similar initial particle sizes (∼100 nm) and displayed minimal change in particle size (hydrodynamic diameter) over a 5-week period stored at 4°C.

**Figure 4.**
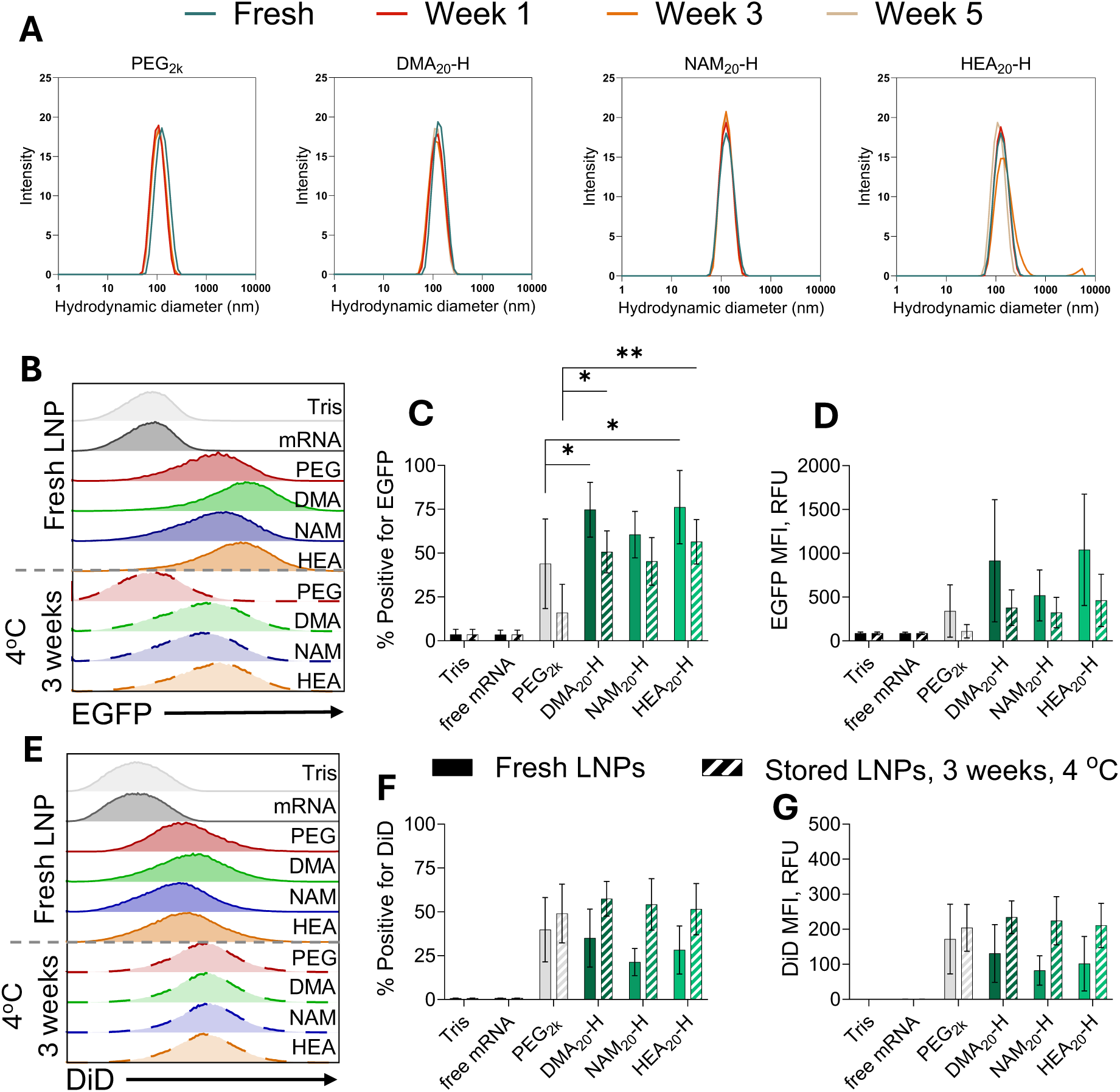
PAM-LNPs retain colloidal stability and biological performance equivalent to PEG-LNPs in fridge storage. PAM- and PEG-LNPs were formulated with eGFP encoding mRNA then stored at 4°C for up to 5 weeks. (**A)** Intensity weighted particle size distributions as measured by dynamic light scattering. Fresh and stored LNPs containing eGFP mRNA and fluorescently labelled with DiD fluorescent dye were incubated with THP-1 cells for 24 h, compared with free mRNA and an untreated Tris buffer vehicle control analysed by flow cytometry (gating strategy figure S11). Data are represented as **B, E** flow cytometry histograms, **C, F** % of positive fluorescent cells and **D, G** median fluorescence intensity for eGFP and DiD respectively. Data are presented as mean ± s.d of *n =* 3 biological repeats. Statistical significance was determined by one way ANOVA against the control PEG-LNPs. \**P* < 0.05, \*\**P* < 0.01.

To investigate the impact of storage on transfection efficiency, 3 week-old and fresh PAM-LNPs (DMA_20_-H, NAM_20_-H, HEA_20_-H) and PEG-LNPs loaded with eGFP mRNA and DiD fluorescent dye were incubated with human monocyte (THP-1) cells. As THP-1s grow in suspension, they are theoretically less susceptible to aggregates that may form during storage. This is because aggregates may sink to the bottom (or float) when left for long periods in suspension, such as the 24 hour incubation when applied to cells. This can alter the local concentration of nanoparticles above a 2D monolayer^56,57^. While all formulations displayed loss in efficacy over 3 weeks of storage, fresh and stored PAM-LNPs containing DMA_20_-H and HEA_20_-H both outperformed PEG-LNPs, showing higher percentage of eGFP transfected cells, when fresh (75% and 50%, respectively) and stored for 3 weeks (50% and 15%, respectively). We suggest efficacy loss seen during storage is likely due to inherent degradation of the mRNA cargo via hydrolysis or reaction with impurities in the formulation^55,58,59^. DLS data shows it is not due to changes in LNP size, and DiD fluorescence data suggest that changes in transfection due to storage cannot be attributed to changes in cellular association/uptake of PAM-LNPs compared to PEG-LNPs. Moreover, as PAM-lipids are likely on the surface of the LNPs (as PEG-lipids are), it is unlikely that they would greatly impact degradation of encapsulated mRNA, which could be why efficacy loss occurs in all formulations regardless of polymer surface. Overall, our findings show that PAM-lipids exhibit equivalent colloidal stability and enhanced efficacy after storage compared to PEG-LNPs, highlighting their suitability as potential PEG alternatives.

### PAMs avoid anti-PEG antibodies and recover lost mRNA expression after repeated dosing of PEG-LNP

The major driver for the development of PEG alternative LNPs is the avoidance of anti-PEG antibodies which reduce the transfection efficiency of encapsulated mRNA. anti-PEG antibodies can compromise LNP integrity leading cargo leakage^11^, and cause ABC of LNPs from the body^7,8,10,60^. This is particularly pertinent after the global rollout of COVID-19 vaccination, where anti-PEG antibody IgM increased 69-fold after patient vaccination with Moderna’s SpikeVax mRNA-LNP formulation, as the composition of which is replicated in this study^5^. It has been previously hypothesised, but not demonstrated, that it may be possible to recover this lost efficacy by dosing LNPs with a PEG-alternative surface, evading opsonisation from anti-PEG antibodies which are unable to bind to PEG-alternatives with different chemical structures^12^.

Therefore we first established that our leading PEG-alternative PAM-lipid (DMA_20_-H) does not bind to anti-PEG antibodies (IgM) across concentrations relevant to clinically determined anti-PEG antibody titres (Figure 5A, S7B)^4,5^. Other PAM-lipids with varied monomer and end group chemistry also show no binding to anti-PEG antibodies (Figure S7A, B). We hypothesised that PAM-LNPs would therefore evade circulating anti-PEG IgM in mice previously injected with PEG-LNPs, and result in enhanced transfection efficiencies compared to the PEG-LNP. To test this, we conducted a repeat dosing experiment comparing three different dosing regimens: 1) a single dose of either PEG-LNPs (1PEG) or DMA_20_-H PAM-LNPs (1DMA); 2) four doses of either PEG-LNPs (4PEG) or DMA_20_-H PAM-LNPs (4DMA) administered weekly; 3) three doses of the PEG-LNP, followed by a fourth dose of the DMA_20_-H PAM-LNP administered weekly (3PEG-1DMA); (Figure 5B). This allowed us to investigate how PAM-LNPs perform in animals with/without circulating anti-polymer antibodies^7,61^. All LNPs were prepared with fLuc mRNA and fluorescent lipid dye DiR to allow independent tracking of mRNA transfection (bioluminescence) from LNP organ biodistribution (fluorescence). Organs and plasma were collected 24 hours after the final dose in each regimen and imaged *ex vivo* or analysed for anti-PEG_2k_ and anti-DMA_20_-H antibody titres, respectively.

**Figure 5.**
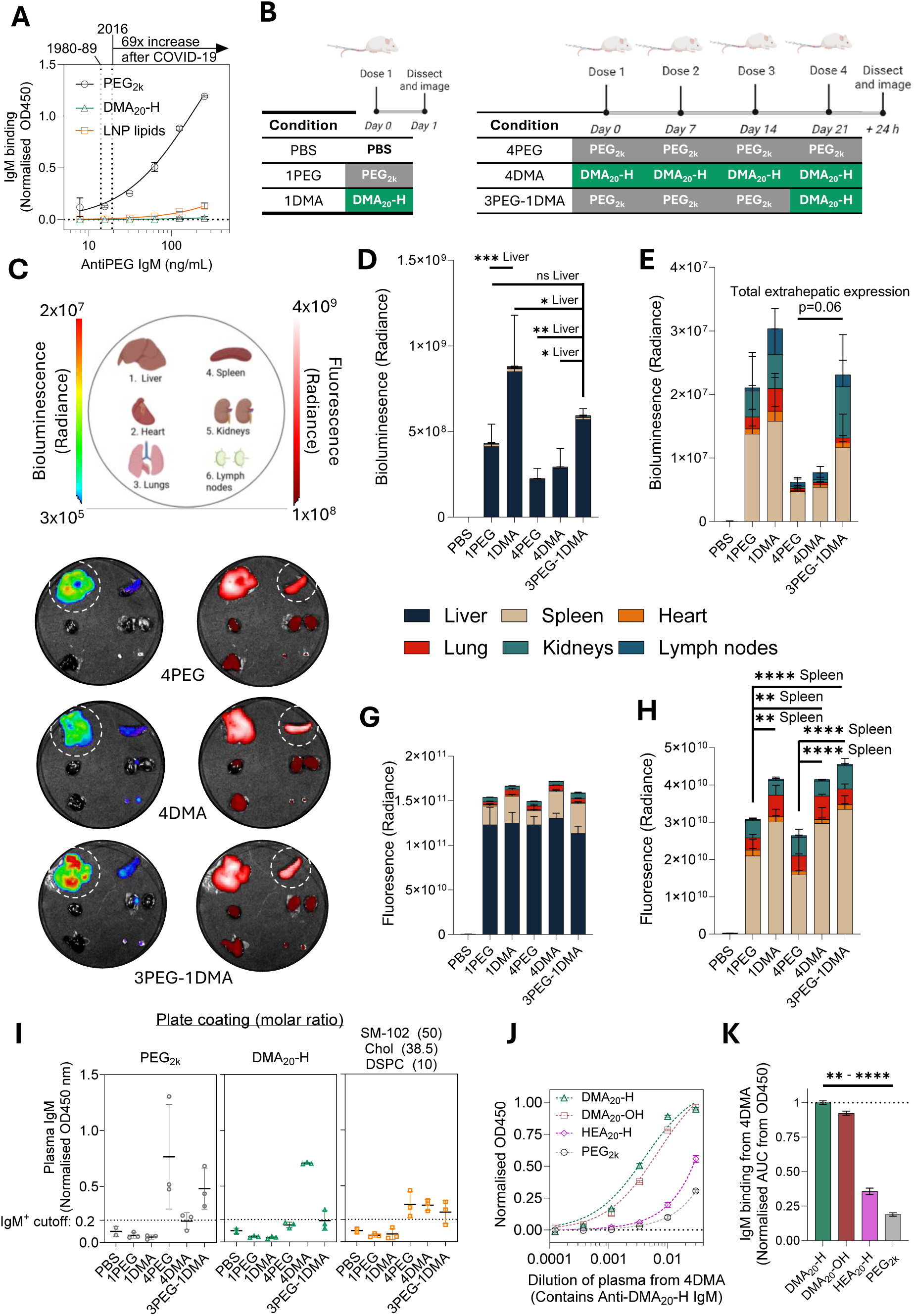
PAM-LNPs recover hepatic and extrahepatic efficacy loss compared to PEG-LNPs when used in heterologous dosing regimen. **A**) Binding of murine anti-PEG IgM to PEG_2k_, DMA_20_-H, or non-polymer LNP components SM102, cholesterol, and DSPC mixed in 50:38.5:10 molar ratios. Dotted lines represent clinically determined anti-PEG IgM levels from specified years before COVID-19, and fold-increase after COVID-19 vaccination^4,5^ **B**) Schematics showing of single dosing and repeat dosing of FLuc mRNA LNPs (5 µg of mRNA per dose) prepared with PAM-lipid DMA_20_-H (green shaded cells) or PEG-lipid PEG_2k_ (grey shaded cells). Organs and blood collected 4 h after final dose (either 1 dose or 4 doses). Organs imaged using *in vivo* imaging system (IVIS) to image bioluminescence (**C**) from mRNA and fluorescence from FLuc mRNA expression and fluorescently tagged LNP (DiR) respectively. Dotted circles highlight organs with statistically significantly differences. Quantified bioluminescence (**D, E**) and fluorescence (**G, H**) determined by regions of interest (equal area between images) are presented as mean of biological repeats ± s.e.m. (n=3 for all LNP conditions; n=2 for PBS vehicle control). Statistical significance was determined by two-way ANOVA with Tukey correction, *P < 0.05. **I** Plasma IgM detected *via* enzyme-linked immunosorption assay (ELISA) of plates coated with specified polymer-lipids/lipids (LNP components). Data were normalised to OD450 nm of PBS control, set to OD450=0.1. IgM^+^ Cutoff set as 2x OD450nm of PBS control. Data presented as mean of biological repeats ± s.d. (n=3) **J**) Binding of IgM in anti-DMA_20_-H Ab^+^ plasma dilutions to different polymer-lipids **K**) Total IgM binding of IgM in anti-DMA_20_-H Ab^+^ plasma determined by area under curve (AUC) integration analysis of figure, normalised to DMA_20_-H **J.** Data for A, J and k presented as mean of technical replicates ± s.e.m.. (n=3), **P < 0.01, ****P < 0.0001

As expected after IV administration, the majority of the fLuc expression was seen in the liver^18^, where after a single dose DMA_20_-H LNPs delivered a 2-fold greater total luciferase expression than the benchmark PEG-LNPs (p<0.05) (Figure 5C-E, Figure S6A-C). This contrasts with IM and ID administration routes, where DMA-LNPs performed comparably to PEG in the liver, suggesting tissue tropism is dependent upon both polymer-lipid type and route of administration. Interestingly, DMA_20_-H LNPs also showed significantly higher propensity to transfect lymph nodes (Figure S6.D) when normalised for LNP biodistribution, consistent with improvements in lymphoid transfection when administered IM and ID. As expected, when the PEG-LNPs were dosed repeatedly, a ∼2-fold drop in luciferase expression was seen, coherent with ELISA assays showing the presence of anti-PEG IgM antibodies (Figure 5D, 5I respectively) and consistent with previous reports^7^. Interestingly a similar reduction in expression was also observed after repeatedly dosing DMA_20_-H LNPs, which also generated anti-DMA IgM. The production of antibodies against polymers other than PEG was not unsurprising, as prior work has reported antibody formation against other established PEG alternatives poly(oxazoline) and poly(sarcosine)^12,14^. Crucially, however, we for the first time demonstrate that we can fully recover all lost hepatic and extrahepatic transfection efficacy back to the level of a single PEG LNP dose (1PEG) by simply swapping only the polymer-lipid used in the final dose formulation, from a PEG_2k_ to DMA_20_-H (3PEG-1DMA condition). This is likely due to DMA_20_-H LNPs ability to evade opsonisation from circulating anti-PEG IgM, as there is minimal cross-reactivity with DMA_20_-H (Figure 5A, S7B). PAM-LNPs are therefore well suited to replace PEG-LNPs in both single-dose and repeated dosing applications, particularly in patient populations with pre-existing anti-PEG antibodies (up-to 90% of people)^5^.

We noted that that while the level of luciferase expression from the final dose of DMA_20_-H LNPs in 3PEG-1DMA fully recovered to the level of 4PEG, which was our key endpoint, it did not entirely recover to the levels of a single dose of DMA_20_-H LNPs in 1DMA. We hypothesised that this effect may arise from opsonisation by non-polymer related antibodies which are generated from LNP doses. To investigate antibodies against non-polymer components within the LNPs (e.g. ionizable lipid, cholesterol, and helper lipids), ELISA assays were conducted using plates coated with all of the lipids found in the PEG LNPs, DMA_20_-H LNPs (Figure 5I, S7C) and a three-lipid mixture corresponding to the LNP formulation but without a polymer-lipid (i.e. only SM-102, DSPC and cholesterol; Figure 5I). Incubating plasma from repeated dosing experiments revealed low levels of “anti-lipid” antibodies in all repeat dose regimens (4PEG, 4DMA and 3PEG 1DMA). These anti-lipid antibodies could be responsible for incomplete recovery of transfection by 3PEG-1DMA to the initial level of 1DMA. The presence of these anti-LNP antibodies is rarely checked but should be monitored more frequently by those assessing the immunogenicity of mRNA-LNPs, particularly as anti-LNP antibodies have also been detected in the population^62^. Nonetheless, our data suggests that anti-polymer antibodies are the main driving force for LNP sensitisation, causing efficacy loss after repeat dosing, and highlighting the importance of this feature in future mRNA-LNP design.

Assessing the biodistribution of LNPs *via* DiR fluorescence highlighted that, aside from the expected liver-dominated biodistribution^63^, PAM-LNPs had increased biodistribution to the spleen over PEG-LNPs (Figure 5C, G, H), consistent with improved cellular association with primary DCs (Figure 1), although the corresponding 1.1- and 2.5-fold increases in splenic transfection were statistically insignificant. Looking broadly across all organs shows that, regardless of dosing regimen, there was not the expected reduction in LNP biodistribution as a result of ABC in mouse groups with high anti-polymer IgM. This indicates that the transfection efficacy loss observed from repeat dosing is not dependent on organ accumulation. This is in contradiction to the conventional wisdom that reduction in efficacy from antibody opsonisation is caused only by increased clearance of nanoparticles^7^. For example, Pilkington and co-workers have shown that the ABC effect of LNPs in blood is minimised when switching LNP surface chemistries for a two dose regimen, however do not report impact on mRNA efficacy^60^. Our data suggest a separate mechanism from ABC as the reason behind reduced efficacy during repeat dosing. Instead, opsonisation *via* anti-polymer antibodies (and potentially some anti-LNP antibodies) may modulate LNP transfection capability, possibly by impacting the cellular response to LNPs (e.g. reducing uptake or endosomal escape) or by destabilising LNPs and causing premature mRNA cargo release^8,11,64^. This is corroborated by Figure S5D, which shows that, when bioluminescence is normalised to LNP biodistribution (DiR fluorescence), the same LNP formulation will transfect the same organs to different degrees depending on the presence/absence of anti-polymer antibodies. The underlying mechanism behind why avoiding anti-PEG antibodies with polymers such as DMA_20_-H results in improved efficacy is the subject of ongoing work. Overall, this is the first demonstration of efficacy recovery in mRNA-LNPs using a heterologous dosing regimen between LNPs with different polymer surfaces, highlighting the potential for PAM-lipids as an alternative to PEG in cases of pre-existing anti-PEG antibodies.

As the reduction in mRNA-LNP efficacy caused by repeated dosing is predominantly mediated by polymer surface and associated antibodies, we then sought to investigate which features of polymer-lipid chemistry are implicated in anti-polymer antibody binding/evasion. Murine plasma positive for anti-DMA_20_-H IgM from the 4DMA condition was tested for binding to different polymers: 1) the original antigen (DMA_20_-H); 2) the same polymer but with a different end group (DMA_20_-OH); 3) a PAM-lipid made of a different monomer (HEA_20_-H); and the non-PAM polymer PEG_2k_ (structures in Figure 1A). Some cross-reactivity is seen for all polymers, likely due to polyclonal cross-reactivity with conserved elements present in the chemistry of all of the polymers, such as glycerol linker or DMG lipid tail. While changing end group has been suggested as a potential route to evade existing anti-PEG antibodies^65,66^, we only see a modest 10% reduction in anti-DMA_20_-H IgM binding with DMA_20_-OH compared to DMA_20_-H. Promisingly, however, we see that changing the monomer chemistry of the PAM from DMA to HEA significantly alleviates cross-reactivity almost as effectively as switching to an entirely different polymer (PEG_2k_) (Figure 5J, 7K). This is particularly encouraging as LNPs prepared with HEA_20_-H were effective transfection reagents in primary cells and *in vivo* (Figure 1D-J). As side-group chemistry changes between polymers made of different PAM monomers, but polymer backbone chemistry is conserved, this suggests that side-group chemistry is the dominant binding epitope and thus is the most judicious aspect of polymer chemistry to redesign for anti-polymer antibody evasion. This is a key benefit of the development of the family of PAM-lipids, as PAM-lipids prepared with every tested monomer chemistry showed effective mRNA-LNP transfection *in vitro* and/or *in vivo*, including HEA_20_-H (Figure 1). We therefore propose that, rather than the one-by-one development of new PEG alternatives, PAM-lipids represent a family of viable PEG-alternative polymers for the future development of mRNA-LNPs.

Interestingly we note that alongside PAMs, aforementioned existing PEG-alternatives poly(oxazoline), and poly(sarcosine), and poly(vinyl pyrrolidine) all possess substituted amide groups in the repeat-units of their polymer chemistry, albeit arranged differently from a structural perspective (Figure S8)^14,15,67^. This suggests that amide-containing polymers may be particularly suited to producing effective PEG-alternative polymers. Within this context, PAM-lipids provide a distinct benefit over these three alternatives (and also PEG) as PAMs have far greater synthetic accessibility of different monomer chemistries as a result of their synthesis *via* RAFT polymerisation^17,68^. PAM-lipids, therefore, are uniquely situated to form the basis of a chemically varied family of viable PEG-alternatives.

### Limitations and future directions

While PAM-lipids effectively replace PEG_2k_ and provide improved mRNA transfection across multiple organs/cell types and during repeated dosing, they do not fully abrogate polymer-directed immune responses. As observed for other polymer-lipids, PAM-lipids remain susceptible to reduced efficacy upon repeated dosing and can still induce anti-polymer antibodies, indicating that their use may require monitoring for potential adverse immune-mediated events, including hypersensitivity, in clinical settings. Importantly, although the higher initial transfection efficiency of PAM-lipids may partially mitigate the impact of antibody formation during repeat administration, any single specific PAM-lipid may not represent a universal solution for all mRNA–LNP applications, highlighting the need for a library of viable PEG-alternatives for formulators to choose from. Consistent with the low cross-reactivity observed between antibodies directed against PEG_2k_ and PAM-lipids, strategies incorporating multiple polymer-lipids while minimising repeated patient exposure to any individual chemistry may be required to maximise therapeutic durability of mRNA-LNPs. In this context, further expansion of the PAM-lipid family through exploration of alternative monomer chemistries represents a promising direction for future investigation. Moreover, due to the ubiquity of PEG in other nanoparticle, biologic and cosmetic formulations, deployment of PAMs could benefit a wide variety of applications outside of gene therapies. Logical starting points would be other formulations susceptible to anti-PEG antibody mediated ABC, such as PEGylated peptides (Pegfilgrastim)^69^ or liposomal chemotherapeutics (Doxil)^70^ .

### Conclusions

We present PAMs as a chemically varied family of polymers suitable for replacing immunogenic PEG in mRNA-LNP formulations. Crucially, we highlight that the polymer chemistry presented on the surface of mRNA-LNPs is a critical design parameter, having profound effects on their absolute efficacy, cell specificity, and antibody evasion despite representing less than 2 mol% of the overall formulation. We observed that small non-hydrophobic end-groups are required to promote stable mRNA-LNP formations with suitable transfection capability, while polymer side-group is a key parameter for attenuating anti-polymer antibody binding. PAM-lipids were found to produce stable LNPs with similar structures to benchmark PEG-lipids. The improved mRNA transfection efficacy bestowed by replacing PEG with PAM lipids is maintained before and after storage. High performing PAM-lipids significantly enhance transfection efficiency in various organs (lymph nodes, liver) *in vivo* and in various immune cell subpopulations *in/ex vivo*, across varied routes of administration. Finally, we demonstrated that PAM-LNPs can efficiently evade opsonisation from circulating anti-PEG antibodies, and hence recover the efficacy lost from repeatedly dosing conventional PEGylated mRNA-LNPs. Together, these data establish the potential role of the PAM-lipid family in next-generation mRNA therapeutics and prophylactics.

## Acknowledgements

We would like to sincerely thank and acknowledge: Dr. Nikita Harvey and the University College London School of Pharmacy Nuclear Magnetic Resonance Core Facility (RRID:SCR_027123); the Cancer Institute flow cytometry technology translation platform, and staff in the UCL Biological services unit; Prof. C. Alexander (University of Nottingham) for kindly gifting the DC2.4 cell line; Dr. Julia Rho (UCL Department of Chemistry) for facilitating access to gel permeation chromatography (GPC) facilities; Dr. Thomas Foran for cryo-TEM services (Birkbeck University of London). The ISIS Neutron and Muon Source (Rutherford Appleton Laboratory, STFC) is acknowledged for SANS beamtime and for access to SAXS (Materials Chracterisation Laboratory).

P.G. was funded by the Royal Society (RG\R1\251152) and MRC (UKRI516). B.F. would like to thank UKRI EPSRC (EP/W524335/1) for funding much of this work. M.S. was also funded by EPSRC EP/W524335/1. N.C. was funded by EPSRC, code: EP/S023054/1. C.G. and C.B. were funded by a PhD studentship from the Percy Stevens Foundation. D.J.S. would like thank the following funding sources: British Heart Foundation (BHF) FS/15/33/31608, FS/SBSRF/21/31020, RM/17/1/33377, Wellcome Trust 212937/Z/18/Z, Medical Research Council (MRC) MR/R026416/1.

Cartoon Illustrations in figures created in part by BioRender Software (Premium Account).

## Data access statement

All relevant data can be obtained upon request from the corresponding authors at. SANS data can be found at 10.5286/ISIS.E.RB2510742-1]

## Conflict of Interest

B. Fiedler and P. Gurnani disclose that they are both named as inventors on a pending UK priority patent application relating to this work recently filed by UCL Business Ltd.

## Materials and Methods

### Materials

All reagents were used as received unless otherwise stated. Materials listed below.

#### Sigma-Aldrich (Merck)

All solvents, cholesterol, 2-(dimethylamino)ethyl acrylate (DMA) monomer, 2-hydroxyethyl acrylamide (HEA), N-acryloylmorpholine (NAM), sodium acetate, Tris buffer (1 M), PBS tablets, poly(A) (poly Adenylic acid), RiboGreen RNA assay kit and dialysis tubing (1 kDa MWCO), RPMI-1640 medium (R0883) Amicon Ultra centrifugal ultrafiltration devices (30–50 kDa MWCO), 3,3′,5,5′-Tetramethylbenzidine (TMB) liquid substrate system for ELISA, Liberase (Cat. No 5401119001). **Tokyo Chemical Industry (TCI):** Hydroxy ethyl acrylamide (HMA), Methoxy methyl acrylamide (MOMOAM) **FUJIFILM Wako Chemicals:** 4,4′-Azobis(4-cyanovaleric acid) (ACVA), Va-044 initiator **Avanti Polar Lipids (Croda, UK):** SM-102, DSPC and DMG-PEG_2k_ **Stratech.** ARCA Cy5 eGFP mRNA (5-moUTP modified) **Tebubio (on behalf of TriLink Biotechnologies):** CleanCap EGFP mRNA (5moU), CleanCap FLuc luciferase mRNA (5moU) **Thermo Fisher Scientific:** Opti-MEM™ I Reduced Serum Medium, DAPI, trypsin, penicillin–streptomycin, Flt3-L cytokine (Cat. No. 550704), ACK lysis buffer (Cat. No. A1049201), DNase I (Cat No. AM2222), DNase buffer (Cat. No AM8170G), and bovine serum albumin (BSA) **Gibco (Thermo Fisher Scientific).** Fetal bovine serum (FBS) was purchased from Gibco (Thermo Fisher Scientific, UK), L-glutamine, non-essential amino acids and β-mercaptoethanol **Promega:** One-Glo™ Luciferase Assay Reagent. **Cayman Chemical:** DiD and DiR lipophilic dyes **GenScript:** Anti-PEG IgM monoclonal antibody (THE™ PEG Antibody, mAb, Mouse; Cat. No. A01795-100) **Invitrogen (Thermo Fisher Scientific)** HRP-conjugated goat anti-mouse IgM secondary antibody (Cat. No. 11839140) was purchased from Merck. **Scientific Laboratory supplies (on behalf of Corning)** High-binding half-area clear 96-well plates

### PAM lipid synthesis – RAFT polymerisation

PAM-lipids were synthesised using commercially available monomers and a synthesised DMG-based macro-CTA (see SI Section 1.2) *via* reversible addition-fragmentation chain transfer RAFT polymerisation. Exact conditions for the CTA and polymer synthesis and purification can be found in the supporting information (SI Sections 1.2.4-1.2.7). A generalised method is presented here. The monomer (∼200 mg) and the DMG-CTA were dissolved in dioxane/water and dioxane respectively and mixed. The final monomer concentration was set to 1.5 - 2 M. ACVA was added from a 10 mg/mL stock solution in dioxane. The mixture was added to a reaction vial equipped with a magnetic stirrer and sealed with a rubber septum. A sample was taken for NMR at t = 0 and the remaining solution was degassed for at least 15 minutes with nitrogen. The vial was then heated to 70 °C for 18 h. The reaction was then stopped by cooling and exposing the solution to air. The progress of the reaction was checked *via* ^1^H NMR. If conversion was deemed to low (below ∼80%) fresh ACVA initiator was added to the vial, and the vial was degassed and re-heated to 70 °C for at 6 - 24 h. Once satisfactory conversion was achieved, the polymer solution was diluted in the minimum volume needed to reduce viscosity to free-flowing liquid and purified by precipitation into organic solvent (e.g. THF, or THF:diethyl ether 1:1 v/v) and dried in a vacuum desiccator. Dialysis was conducted on redissolved samples against at least 100-fold the volume of deionised water, for at least 36 hours, with at least 2 water changes (1 kDa MWCO). Where no suitable precipitation solvent was found, polymers were first dialysed against at least 100--fold the volume of 2:1 v/v IPA:H_2_O for 12 hours, before dialysis against deionised water as above for 36 hours. The resulting solution was freeze dried and polymers were characterised *via* ^1^H NMR and GPC. Aliquots of these polymers were taken for replacement of the –S_3_C_5_H_9_ end group with either - H or -OH (methods and characterisation detailed in SI Sections S1.2.4-1.2.7). Stock solutions of these polymers were prepared to 1 mM in DMSO and stored at -20 °C.

### Preparation of LNPs

A lipid mix of SM-102:DSPC:Cholesterol:PEG-lipid or PAM-lipid at 50:10:38.5:1.5 molar ratio was prepared in ethanol by mixing stock solutions of lipids (10 mg/mL in ethanol) and PEG/PAM-lipids (1 mM in DMSO). LNPs were prepared by rapid injection of the lipid mix into aqueous mRNA *via* pipette-mixing^21^. The ethanol:aqueous v/v ratio was 3:1, with N/P ratio = 6, with a final mRNA concentration of either 25 µg/mL (*in vitro* experiments) or 50 µg/mL (*in vivo* experiments). mRNA encoding either eGFP or luciferase, or poly(a), was diluted in sodium acetate buffer (50 mM, pH 5.2). The resulting LNP suspension was incubated at room temperature for ∼10 minutes before being diluted with 1.5-fold the existing concentration of Tris buffer (50 mM, pH 7.4) and stored at 4 °C for a maximum of 24 hours before use (unless specified elsewhere). Particle sizes were characterised *via* (DLS) after further dilution in Tris buffer (50 mM, pH 7.4) to an mRNA concentration of 2.5 µg/mL. Encapsulation efficiency was determined by the RiboGreen assay, which was performed as per supplier instructions. *Buffer exchange:* LNPs for *in vivo* applications were buffer exchanged and concentrated with either Tris buffer 50 mM (IM, ID injections) or PBS (IV injections) *via* dialysis and/or centrifugal ultrafiltration (MWCO 30-50 kDa). *Fluorescent dyes:* For *in vitro* experiments in THP-1s DiD dye was incorporated into the ethanol phase at 200:1 molar ratio of lipids:DiD. For *in vivo* biodistribution experiments, DiR dye was incorporated into the ethanol phase at 50:1 molar ratio of lipids:DiR. For *ex vivo* experiments in BMDCs, Cy5-tagged eGFP-mRNA was used (Stratech, Cat. No. R1011-APE).

### Subculture of DC2.4 cell line

DC2.4 cells (immortalised murine dendritic cells) were thawed and sub-cultured according to supplier recommendations. Briefly, they were grown in sterile conditions as 2D monolayers in cell-culture treated T-75 flasks and kept at 37 °C, 5% CO_2_ (humidified). Cells were grown in ∼10 mL of media comprised of RPMI-1640 (Sigma Cat.No. R0883), 10% FBS (Gibco, Fisher UK), 1X L-Glutamine (Cat. No. TMS-002-C), 1X non-essential amino acids (Cat. No. TMS-001-C), and 0.0054X β-mercaptoethanol (Cat. No. ES-007-E). Cells were passaged at 50-90% confluency (every 1-4 days) at 1:3-1:12 ratios for a maximum of 10 passages. To achieve this, cells were detached from the flask by incubating in 2-3 mL Trypsin (5%) for ∼5-10 minutes at 37 °C and agitating mechanically. Trypsinisation was stopped by addition of complete media. Cells were centrifuged at 1500 rpm for 5 minutes to obtain a cell pellet. Supernatant was discarded before resuspension in complete media, and the relevant dilution of cells added to a new subculture flask containing ∼10 mL of fresh media. Cells were frozen in 1-2 mL complete media + 10 % DMSO v/v and stored at -80 °C or in liquid nitrogen vapour before use (2-5 ൷10^6^ cells per vial).

### *In vitro* screening in DC2.4 cells

DC2.4 cells were seeded at 20,000 cells per well in a white cell-culture treated 96 well plate in their regular growth medium. After 23 hours, media was removed and cells were rinsed once with 200 µL of PBS. Next, mRNA-LNPs prepared with either PEG-lipids or PAM-lipids and containing FLuc-mRNA (10 µg/mL, in Tris, 50 mM) were diluted 1:10 to an mRNA concentration of 1 µg/mL in either OptiMEM (low serum conditions) or OptiMEM + 10% FBS (high serum conditions) and incubated for 15 minutes. 100 µL was then added to each well (3-4 wells per condition). Negative controls of no treatment, only Tris buffer (50 mM), and free mRNA were also tested. A positive control of Lipofectamine messenger-MAX was used as per the supplier instructions. After 24 hours, the treatments were removed and the cells gently rinsed twice with 200 µL of PBS. 50 µL of One-Glo™ reagent was then added to each well and analysed using the One-Glo™ protocol on the Promega GloMax plate reader. Values presented as fold-change are relative to the PEG condition calculated *via* the following equation.

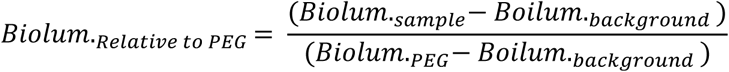

(Where Biolum. = measured bioluminescence of the One-Glo reagent after addition to cells (relative luminescent units, RLU)

### Subculture of THP-1 cells

THP-1 cells (human monocyte cells) were thawed and sub-cultured according to supplier recommendations. Briefly, they were grown in sterile conditions in suspension in T-75 flasks, and kept at 37 °C, 5% CO_2_ (humidified). Cells were grown in ∼15 mL of media comprised of RPMI-1640 (Sigma Cat.No. R0883), 10% FBS (Gibco, Fisher UK), 1X L-Glutamine (Cat. No. TMS-002-C), 1X non-essential amino acids (Cat. No. TMS-001-C), and 0.0054X β-mercaptoethanol (Cat. No. ES-007-E). Cells were passaged at 50-90% confluency (every 1-4 days) at 1:3-1:12 ratios for a maximum of 10 passages. To achieve this, cells were detached from the flask by incubating in 2-3 mL Trypsin (5%) for ∼5-10 minutes at 37 °C and agitating mechanically. Trypsinisation was stopped by addition of complete media. Cells were centrifuged at 1500 rpm for 5 minutes to obtain a cell pellet. Supernatant was discarded before resuspension in complete media, and the relevant dilution of cells added to a new subculture flask containing ∼15 mL of fresh media.

### *In vitro* uptake and stability study transfection THP-1

THP-1s were harvested from their growth flasks and centrifuged to a cell pellet to remove growth media before being resuspended to a concentrated stock in OptiMEM. 250k live cells were then diluted in OptiMEM and mixed with eGFP-LNPs containing fluorescent dye DiD to a final volume of 500 µL, and a final concentration of mRNA of 1 µg/mL. This suspension was added to a 24 well plate and incubated for 24 hours at 37 °C, 5% CO_2_ for 24 hours. After 24 hours, cells were rinsed with cold PBS, incubated with DAPI (1:1000 in PBS) for 15 minutes, followed by 3 rinses with 2% FBS in PBS and a final resuspension in 2% FBS in PBS and kept on ice while flow cytometric analysis was conducted.

### Isolation and culture of BMDCs

Bone marrow (BM) was crushed from C57BL/6 mice femurs and tibias to generate a single cell suspension in complete (c) RPMI, RMPI 1640 supplemented with 10% FCS, 1% glutamine, 1% P-S, 1 mM sodium pyruvate, 1X MEM NEAA, 10 mM HEPES, and 50 µM β-mercaptoethanol. Cells were plated at 2×10^6^ cells/well in non-tissue cultured treated 24 well plate with 150 ng/ml Flt3-L at 37 °C, 5% CO_2_. Cells were left to culture for 9 days.

### *In vitro* transfection of BMDCs

On the penultimate day of the BMDC culture (day 8), media was carefully removed and replaced with 500ul per/well Opti-MEM™ I Reduced Serum Medium with 1 ng/ml LNP for 4-6 hours. After incubation 1ml per well of cRPMI was added and cells were incubated for a further 24 hours at 37 °C, 5% CO_2_.

### *In vivo* procedures

All animal studies were approved by the University College London Biological Services Ethical Review Committee and performed with UK Home Office approval (PPL 70/8709, PP1692884, PP4506002). Animal work conformed to the UK Animals (Scientific Procedures) Act, 1986 and Directive 2010/63/EU of the European Parliament. Animals were assigned to experimental groups at random. Researchers were not blinded experimental conditions after groups were assigned.

### *In vivo* intramuscular (IM) injection of mRNA-LNPs

LNPs prepared with either PEG-lipids (DMG-PEG_2k_) or PAM-lipids containing FLuc mRNA (2.5 µg, 50 µL, 50 µg/mL) were injected into each hind quadriceps of Balb/c mice. Bioluminescence imaging was performed using an IVIS Spectrum (Revvity UK) at 24 and 48 hours. Mice were given anaesthesia using 1–2% isoflurane in 100% oxygen and maintained at 37 °C using an integrated heated bed. D-luciferin was administered intraperitoneally (IP) (15 mg/mL, 100 uL, Promega) and images were acquired consistently over a ∼30 minute period until bioluminescence peaked, (open filter, auto exposure, F/Stop = 2, medium binning) as described elsewhere^71^. Data in figures/text correspond to peak bioluminescence recorded. After 48 hours, mice were sacrificed and organs harvested and stored at -80 °C. Organs were thawed and submerged in warm D-luciferin solution (15 mg/mL) for 5 minutes before bioluminescence was measured at 37 °C using the IVIS Spectrum.

### *In vivo* intradermal (ID) injection of mRNA-LNPs

LNPs prepared with either PEG-lipids (DMG-PEG_2k_) or the PAM-lipid DMA_20_-H containing eGFP mRNA (3 µg, 60 µg/mL, 50 µL) were injected into each shaved quadriceps of C57BL/6 mice. After 24 or 48 hours, spleen, liver, skin both at the site of injection and far from the site of injection, dLN and cLN were collected and processed for flow cytometry.

### Processing organs for flow cytometry

#### Spleens

Spleens were dissociated through a 70uM filter in 1ml PBS using the end of a 1ml syringe. The filter was washed, and cells were centrifuged in 10ml of PBS at 400 x g for 5 minutes at room temperature. Splenocytes were then resuspended in 1 mL of ACK lysis buffer for 1 minute at room temp. 10mL of cRPMI was added and the cells were counted.

#### Liver

Livers were manually dissected with scissors in 1ml digestion buffer mix (1X DNase buffer, liberase 25 µg/ml, 0.625 µg/ml DNase I in PBS) and incubated in 37 °C, 5% CO_2_ for 1 hr. Cells were filtered through a 70 μm filter and centrifuged at 400 x g for 5 minutes at 5 °C. Cells were then resuspended in 1 mL of ACK lysis for 1 minute at room temp. 10mL of cRPMI was added, and the cells were centrifuged at 400 x g for 5 minutes at 5 °C and the supernatant removed from the cell pellet. 10mL of cRPMI was added and cells were counted.

#### Skin

Skin from both the site of injection and the other side of the mouse was shaved and placed in 1 mL of digestion buffer mix (1X DNase buffer, liberase 25 µg/ml, 0.625 µg/ml DNase I in PBS). Skin was minced using scissors and incubated in 37 °C, 5% CO_2_ for 1hr. Cells were then filtered using a 70um filter and washed with 5 mL of PBS at 400 x g for 5 minutes at 5 °C. 1mL of cold cRPMI was added and the cells were counted.

#### Lymph nodes (LNs)

LNs were dissociated on ice, on a 70um filter in 1 mL cold cRMPI using the end of a 1 mL syringe. The filter was washed, and cells were centrifuged with 5 mL of cold cRMPI at 400 x g for 5 minutes at 5 °C. After the supernatant was removed from the cell pellet, 1mL of cold cRPMI was added and cells were counted.

### *In vivo* Single and repeated intravenous (IV) dosing of mRNA-LNPs

LNPs prepared with either PEG-lipids or the PAM-lipid DMA_20_-H containing FLuc mRNA (5 µg, 50 µg/mL, 100 µL) and DiR dye (50:1 lipid:dye molar ratio) were injected into the tail vail of Balb/c mice. For live whole-body imaging, after 18 hours D-luciferin was injected IP (15 mg/mL, 100 uL) and mice were live imaged using the IVIS Spectrum as described above. For organ biodistribution imaging at 24 hours, D-luciferin was injected IP (15 mg/mL, 100 µL) and . After 4 minutes blood was collected *via* cervical puncture into EDTA-coated tubes, and organs harvested and underwent immediate bioluminescence and fluorescence (excitation/emission 745/800) imaging at 37 °C using the IVIS Spectrum.

### Enzyme-linked immunosorbent assay (ELISA) for anti-Polymer and anti-LNP antibodies

#### Diluted plasma isolation

Diluted plasma was isolated from whole blood by diluting 1:2 with PBS and centrifuging for up-to 15 minutes and collecting the supernatant. The supernatant was stored at -80 °C until used.

#### ELISA protocol

ELISA protocol was adapted from exising protocols^72,73^. Low-volume, high-binding clear 96 well plates were coated with PEG, DMA, PEG–LNP, DMA–LNP, or no-polymer lipid mixes (50 uL, 0.2 mM total lipids, 10 nmol per well, 80:20 Ethanol:DMSO) and left to incubate for 1 hour at room temperature to allow insertion/adsorption of lipids into/onto the well surface, followed by three washes with 60 µL of 50 mM Tris buffer (pH 7.5). Wells were blocked for 90 min with 150 µL of 2% (w/v) bovine serum albumin (BSA) in Tris-buffered saline (TBS; 50 mM Tris, 0.145 M NaCl, pH 7.5) then rinsed once with 0.25% BSA in TBS. Diluted plasma samples were further diluted 1:33.3 in 0.25% BSA in TBS, which results in a 1:100 final plasma dilution due to prior 1:2 dilution of plasma in PBS. 50 uL of these plasma samples, or primary anti-PEG IgM antibodies (1:2000-1:64,0000 serial dilution, 0.25% BSA in TBS, with further dilutions as required) were added to each well and incubated for 1 h at room temperature then removed. Plates were rinsed 3 times with 0.25% BSA in TBS. Wells were then re-blocked with 150 µL of 2% BSA in TBS for 30 min, followed by one additional rinse with 0.25% BSA in TBS. 50 uL of horseradish-peroxidase (HRP) linked secondary antibody (1:4000 dilution; 20 µL in 20 mL 0.25% BSA in TBS) was added and incubated for 1 h at room temperature. Plates were then washed six times with 0.25% BSA in TBS prior to detection. The chromogenic substrate 3,3’-,5,5’-tetramethylbenzidine (TMB) was added, and absorbance was monitored at 650 nm using a plate reader. The reaction was stopped by the addition of 2 N H₂SO₄ after a maximum of 5 min, or earlier when a maximum optical density of 0.4 was reached at 650 nm. The optical density at 450 nm (OD450) was then measured using a plate reader.

### Instrumentation and Analysis

#### NMR Spectroscopy

All ^1^H-NMR spectra were recorded in ppm (δ) at 400 MHz in d6-DMSO using a Bruker Advance III MHz spectrometer that was maintained at 25°C. To analyse the spectra MestReNova 6.0.2 copyright 2009 (Mestrelab Research S.L.) was applied.

#### Theoretical Molar Mass Calculation

The theoretical number average molar mass (*M*_n,th_) was calculated as demonstrated in the following way:

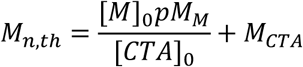

**Equation 2** Equation for the theoretical number averaged molecular weight (Mn,th). [M]_0_ and [CTA]_0_ are the initial concentrations (in M) of monomer and chain transfer agent respectively. *p* is the fractional monomer conversion as determined by ^1^H NMR spectroscopy. M_M_ and M_CTA_ are the molecular weights (g mol^-1^) of the monomer and chain transfer agent respectively.

#### Gel permeation Chromatography

The gel permeation chromatography (GPC) analysis was performed using a Shimadzu I-series GPC system equipped with a UV/Vis detector (wavelength set to 309 nm) and differential refractive index (DRI) detection. The system was equipped with a PLgel 5 μm guard column and a 2 × PLgel Mixed D columns (300 × 7.5 mm). Dimethylformamide (DMF) with 0.1% LiBr was utilised as an eluent with a flow rate of 1 mL/min at 50°C. The instrument was calibrated through Poly(methyl methacrylate) standards (Agilent EasyVials) between 550-955500 g mol^−1^. Using conventional calibration, Cirrus GPC software, dispersity (*Đ*) values, molecular weight (*M_w_*) and experimental molar mass (*M*_n,GPC_) were detected.

#### Dynamic light scattering

Dynamic light scattering (DLS) for 30 PAM-LNP formulations, PEG-LNP formulation, and PEG-free LNP formulation (Figure 1A) was conducted with a DynaPro NanoStar Plate reader (Wyatt Technology) at 25 °C to characterise the particles in terms of the size and PDI.

For all other dynamic light scattering (DLS) results (e.g. figure 3), a Zetasizer Ultra (Malvern, Inc.) instrument was used to characterise the particles in terms of the size, PDI, at 25°C, scattering angle 173°.

#### Small-angle X-ray scattering

SAXS measurements of LNPs ([LNP] ≈ 30 mg/mL in 50 mM Tris-HCl pH=7) were performed on the Nano-inXider (Xenocs, Grenoble, France) using a microfocus sealed tube Cu 30W/30mm X-ray source with λ = 1.54 Å and two Dectris Pilatus 3 hybrid photon counting detectors. Samples were loaded and sealed in borosilicate glass capillary tubes of 1.0 mm thickness, which were mounted on a sample holder. Data were collected over a scattering wavevector 𝑞 = ^4𝜋^⁄_𝜆_ sin(^𝜃^⁄_2_) range of 0.0045 – 0.37 Å^-1^, where *θ* is the scattering angle. Samples and their corresponding buffer were measured in borosilicate capillaries at room temperature. The 2D scattering pattern was radially averaged and the background buffer was subtracted using Foxtrot software (<u>SWING | French national synchrotron facility</u>).

#### Small angle neutron scattering

SANS measurements of LNPs were performed on the Sans2d beamline at the ISIS Neutron and Muon Source (Rutherford Appleton Laboratory, Oxfordshire, UK). The instrument configuration used a 12 m collimation, neutrons with a wavelength range of 1.75 – 12.5 Å in time-of-flight mode and two detectors placed 5 and 12 m from the sample, respectively, giving a q-range of 0.0015 – 0.5 Å^-1^ in 2-mm thick quartz cuvettes mounted in a temperature-controlled sample changer. Samples and their corresponding buffer were measured at 25 °C and [LNP] ≈ 3 mg/mL in 50 mM Tris-DCl with 100% D_2_O (pD 7.4).

Raw 2D data were corrected (incident neutron wavelength distribution, detector efficiency, measured transmission and sample volume),were radially averaged, scaled to absolute intensity, merged and background subtracted using Mantid Workbench^74,75^. The 1D reduced data corresponds to the coherent elastic differential scattering cross-section [*∂Σ*/*∂Ω*(q)], commonly referred to as intensity, *I* [cm^-1^].

#### Small-angle neutron/X-ray scattering data analysis

All the SANS/SAXS data were analysed using SasView (version 6, https://www.sasview.org/) for model fitting and BioXTAS RAW 2.3.1 (https://bioxtas-raw.readthedocs.io/en/latest/) for shape-independent fitting (inverse Fourier transform and Guinier approximation).

SAXS data were fitted, where appropriate, using the broad peak model in SasView to infer the internal structure of LNPs. The pair distance distribution function *P*(*r*) analysis was performed on all SAS data using the RAW^76^ implementation of the indirect Fourier transform method GNOM^77^, to determine the maximum dimension (D_max_) and the radius of gyration (R_g_), and estimate the shape of the particles. DENSS algorithm (density from solution scattering) (doi:10.1038/nmeth.4581) was then used to calculate the ab initio density map directly from the GNOM output and reconstruct the three-dimensional structure as described in. The D_max_ and R_g_ values from SAXS were consistently lower than those from SANS, due to the limited q-range accessed with SAXS. Guinier analysis was also performed on the SANS data and the R_g_ values determined were compared to the ones derived from *P*(*r*). The SANS data were then fitted using SasView with a core-shell ellipsoid model (CSE) and a cylinder model with a core and two shells. The core is assumed to contain the RNA and most of the SM-102 and cholesterol, the first shell is made up of DSPC and DMG, and the second shell represents the PEG layer. The core radius, shell thickness, SLD of the shells were allowed to vary with all other parameters estimated and fixed. The fittings were optimised using the DREAM fit algorithm.

#### Cryo-electron microscopy

mRNA-LNP samples (1.5 mg/mL of total lipids; eGFP mRNA) were prepared for cryo-electron microscopy (cryo-EM) using a GP2 plunge freezer (Leica Microsystems). 3 µL of LNP was applied to glow-discharged 300-mesh copper grids with a lacey carbon support film (Agar Scientific). Grid preparation was performed at 4 °C and 85% relative humidity. After application of the sample, the instrument was programmed with a 40 s wait time, followed by a 7 s front blot and 0 s drain time prior to vitrification. Grids were rapidly plunged into liquid ethane at −183 °C, then transferred to liquid nitrogen for storage.

Grids were clipped and loaded into a Titan Krios G3i transmission electron microscope (Thermo Fisher Scientific) equipped with a K3 direct electron detector and BioQuantum energy filter (Gatan). Data were collected in counting mode with a total electron dose of 85.4 e⁻/Å², nominal magnification of 105,000×, defocus of −3 µm, and a 20 eV energy slit width. Grids were first screened for ice thickness and particle distribution. Integrated images were saved as 2×-binned JPEG files, corresponding to a pixel size of 1.6712 Å.

### Statistics

Statistics were calculated using GraphPad prism computer software (version 10.6.0 and 10.6.1)

