## Supplementary Information for "Poly(acrylamido) PEG-alternatives Enhance mRNA-LNP Efficacy in Immune Cells and Evade Anti-PEG Antibodies after Repeated Dosing"

### 1 Supporting Information

### 1.1 Supporting results and discussion

#### 1.1.1 Characterisation and screening of library of 30 PAM-LNPs

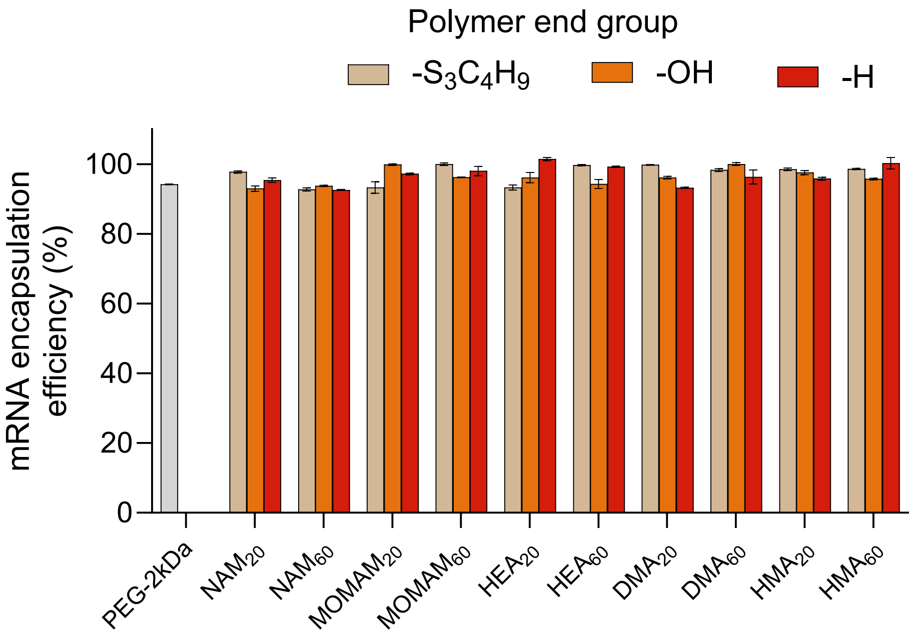

**Figure S1 Encapsulation efficiency of PEG-LNPs and PAM-LNPs** Determined by ribogreen assay (following manufacturers protocols). Data presented as mean of 3 technical replicates. Data presented as mean. Error bars represent ± s.d.

#### 1.1.2 Cytotoxicity of PAM-LNPS and PEG-LNPs

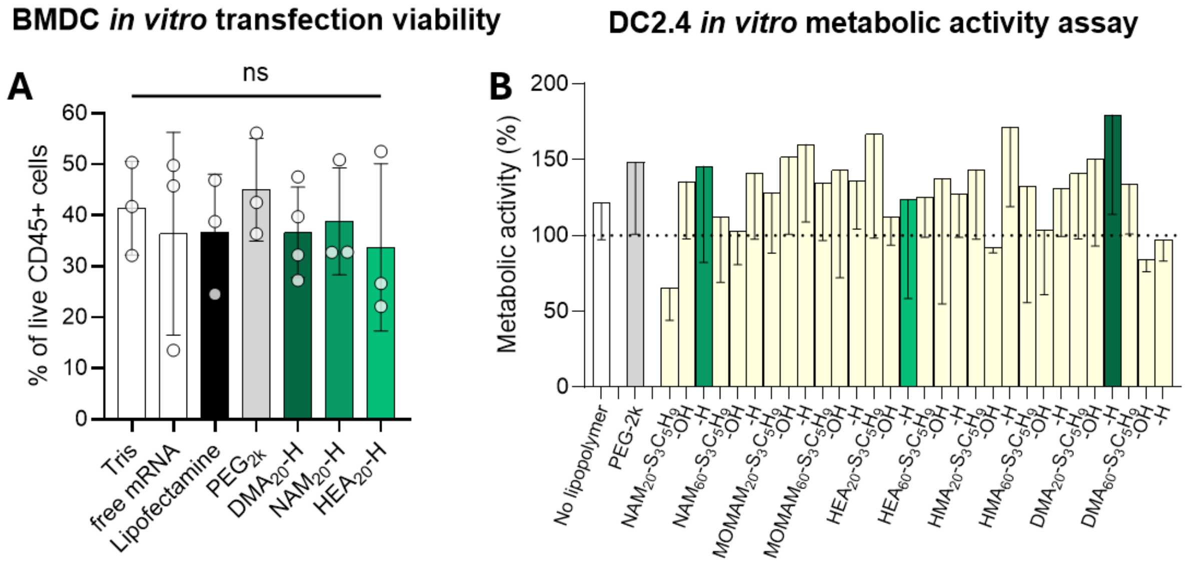

**Figure S2. PAM-LNPs are not cytotoxic and are well tolerated by immune cells. A** percentage of live CD45+ cells in BMDC cultures treated with vehicle control (tris), controls, or LNP formulations prepared with PEG-lipids or PAM-lipids for 24 hours ahead of analysis via flow cytometry. Data presented as mean biological repeats (n=3-4) ± s.d.. Statistical significance was determined by one way ANOVA with Dunnett correction. **B** Formulation of PAM-LNPs from DMG-based PAM-lipids with varied chemistries (5 monomers, 2 end-groups and 2 molar masses) and control PEG-LNPs, were incubated with DC2.4 cell line for 24 hours. Metabolic activity was determined via

PrestoBlue assay. Data presented as % of metabolic activity relative to cells treated with vehicle control (Tris buffer) and kill condition (1% Triton X-100) as the mean of $n$ = 3 biological repeats, error bars represent s.d..

##### 1.1.3 Effect of polymer end-group on mRNA-LNP transfection in DC2.4s

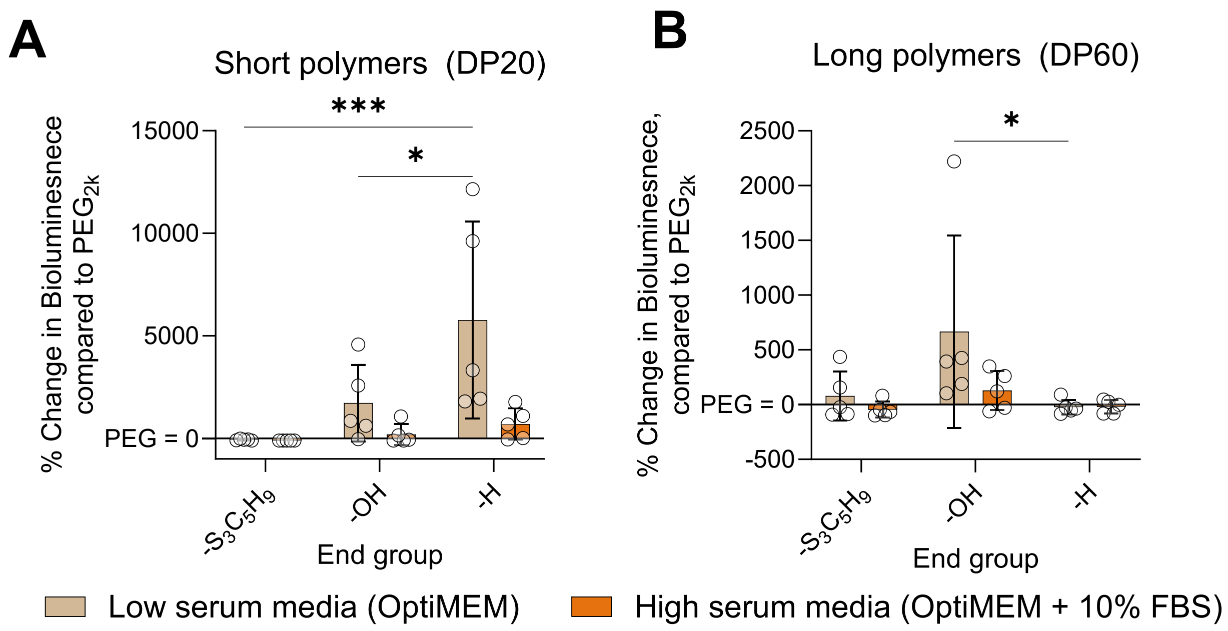

**Figure S3. End-group removal of PAM-lipids improves transfection of mRNA-LNPs more for shorter DP20 polymers than for longer DP60 PAMS (relative to PEG-LNPs).** Change in bioluminescence of DC2.4 cells after 24 hour incubation of with PAM-LNPs prepared with either short (**A**) or long (**B**) PAM-lipids. Each datapoint represents the mean of 3 biological repeats for LNPs prepared with a specific polymer with a different monomer/backbone chemistry to others within the group. Data presented as mean. Error bars represent ± s.d. Significance determined by two-way ANOVA with Tukey correction. *$P$ < 0.05, **$P$ < 0.01., ***$P$<0.001.

#### 1.1.4 PAM-LNPs and PEG-LNPS in primary BMDCS

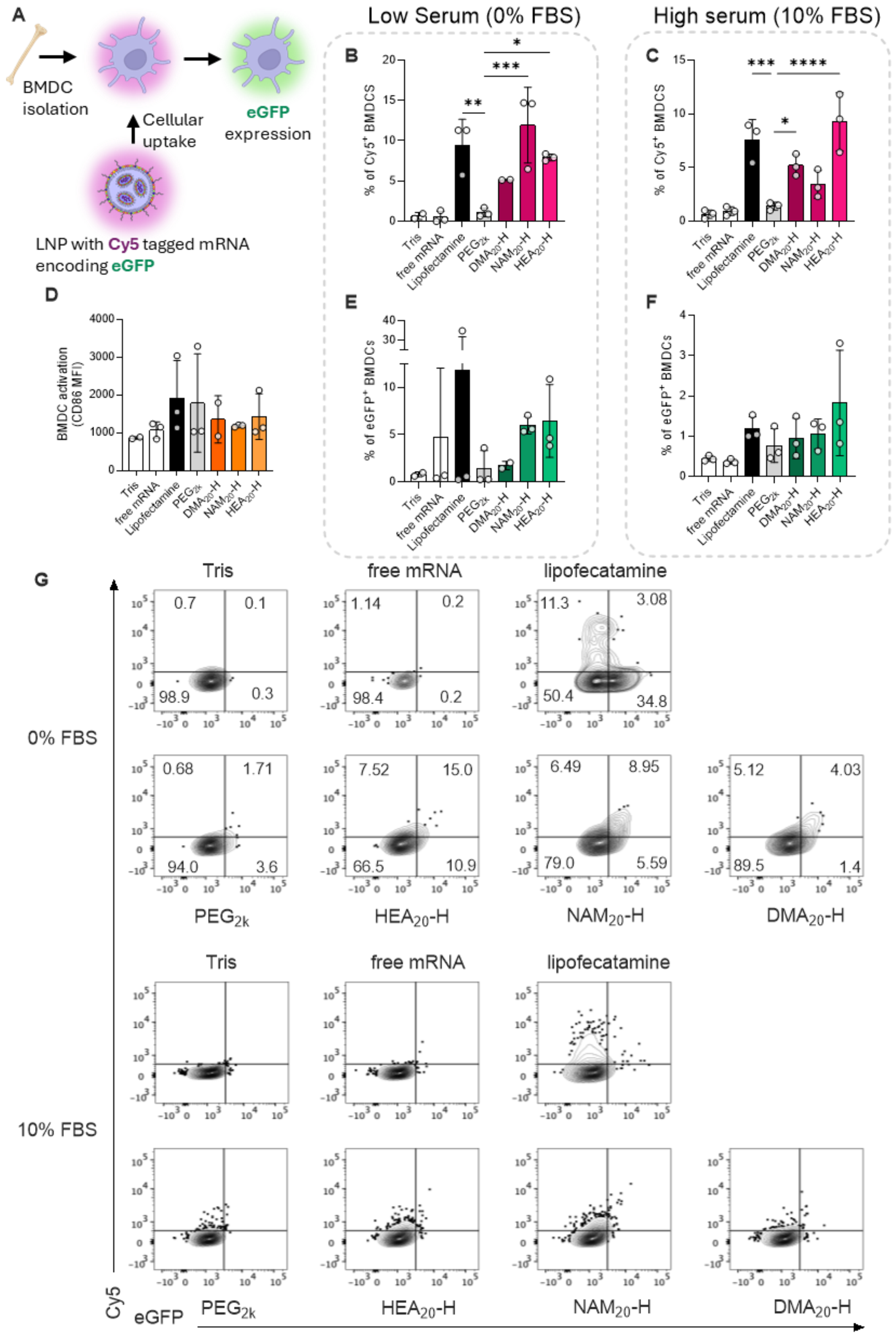

**Figure S4. PAM-LNPs show superior association/internalisation in BMDCs than PEG-LNPs** Internalisation/association (**A**) and transfection (**B**) of BMDCs by PAM-LNPS or PEG-LNPs containing Cy5-tagged mRNA encoding eGFP, and corresponding BMDC activation as measured by CD86 (**C**). Representative flow cytometry plots (**D**) of eGFP (X axis) Cy5 (Y axis). Data presented as mean of biological repeats (n=3 for all samples, except Tris and $DMA_{20}$-H where n=2) ± s.d.. Significance determined by one way ANOVA with Dunnett correction. *$P < 0.05$, **$P < 0.01$., ***$P<0.001$.

PAM-LNPs, and to a lesser extent PEG-LNPs, show a general population shift from $Cy5^-$/$eGFP^-$ to $Cy5^+$/$eGFP^+$, suggesting a correlation between mRNA uptake/association and eGFP expression. Interestingly these behave differently to lipoplex Lipofectamine, where the cell population bifurcates into $Cy5^+$/$eGFP^-$ and $Cy5^-$/$eGFP^+$. Improvement in eGFP expression by PAM-LNPs compared to PEG-LNPs is not statistically significant, which appears in contrast to the data in figure 1 showing significant improvements when using eGFP-mRNA, rather than Cy5-tagged mRNA. This is likely due to our observation that Cy5-tagged mRNA does not transfect as efficiently as unlabelled mRNA (confirmed by manufacturer).

#### 1.1.5 SANS and SAXS analysis of PAM-LNPs and PEG-LNPs

**Table S1 –** Maximum dimension ($D_{max}$), radius of gyration ($R_g$), and shape factor of the LNP samples calculated using GNOM IFT on the SANS data. The shape factor is calculated with $D_{max}/R_g$ from P(r) analysis where the shape factor of a sphere is 2.58 and a prolate ellipsoid is 3.[1]

| **Sample** | **$D_{max}$** | **$R_g$** | $\chi^2$ | **Shape factor** |
|---|---|---|---|---|
| $DMA_{20}$-H | 1575 | 490.7 ± 3.02 | 0.9253 | 3.21 |
| $HEA_{20}$-H | 1584 | 480.4 ± 4.13 | 0.8516 | 3.30 |
| $NAM_{20}$-H | 1556 | 500.4 ± 2.09 | 0.8885 | 3.11 |
| $PEG_{2k}$ | 1343 | 426.1 ± 2.20 | 0.9638 | 3.15 |

**Table S2** – Radius of gyration (Rg) estimated using Guinier analysis on the SANS data with BioXTAS RAW 2.1.1.

| **Sample** | **Rg (Å)** | **$r^2$ fit** | **q min*Rg** | **q max*Rg** |
|---|---|---|---|---|
| $DMA_{20}$-H | 573.83 ± 21.59 | 0.9480 | 1.0558 | 1.4977 |
| $HEA_{20}$-H | 510.88 ± 13.79 | 0.9078 | 0.7868 | 1.4969 |
| $NAM_{20}$-H | 659.30 ± 50.07 | 0.9376 | 1.0153 | 1.4439 |
| $PEG_{2k}$ | 477.47 ± 10.09 | 0.9934 | 0.8785 | 1.4849 |

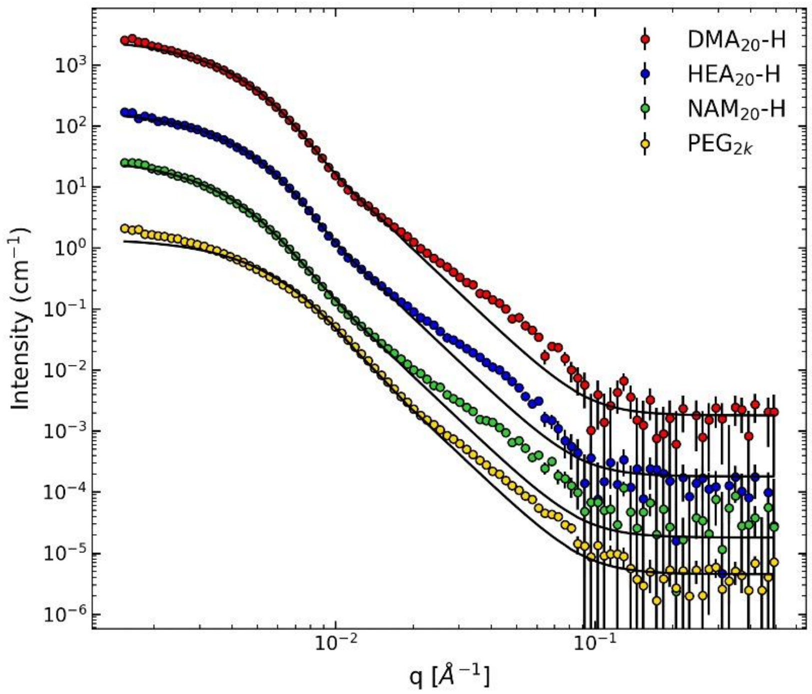

**Figure S5.** SANS data of LNPs prepared with PEG2k, DMA20-H, NAM20-H, or HEA20-H fitted using core-shell ellipsoid model with SasView and DREAM algorithm.

**Table S3** – Fitted parameters and $\chi^2$ from SANS using core-shell ellipsoid model with SasView and DREAM algorithm.

| **Sample** | **Core radius** | **x core** | **Shell thickness** | **SLD core** | **SLD shell** | **Core PDI** | **Shell PDI** | $\chi^2$ |
|---|---|---|---|---|---|---|---|---|
| $DMA_{20}$-H | 338.86 ± 8.10 | 2.00 ± 0.07 | 21.39 ± 4.05 | 0.72 ± 0.08 | 6.19 ± 0.23 | 0.29 ± 0.02 | 0.02 ± 0.12 | 29.18 |
| $HEA_{20}$-H | 334.55 ± 10.0 | 1.94 ± 0.19 | 20.14 ± 10.22 | 1.54 ± 0.18 | 6.18 ± 0.24 | 0.28 ± 0.03 | 0.02 ± 0.12 | 20.23 |
| $NAM_{20}$-H | 327.53 ± 11.2 | 1.98 ± 0.10 | 20.36 ± 5.86 | 1.12 ± 0.11 | 6.19 ± 0.31 | 0.36 ± 0.02 | 0.17 ± 0.13 | 26.20 |
| $PEG_{2k}$ | 201.39 ± 3.73 | 2.00 ± 0.03 | 20.02 ± 1.81 | 0.60 ± 0.06 | 6.16 ± 0.23 | 0.40 ± 0.01 | 0.21 ± 0.13 | 60.72 |

**Table S4** – Fitted parameters and $\chi^2$ from SANS fitting using core-two-shell cylinder model with SasView and DREAM algorithm.

| **Sample** | **Core radius** | **Core length** | **Shell thickness 1** | **Shell thickness 2** | **SLD core** | **SLD shell 1** | **SLD shell 2** | **Core PDI** | **Shell 1 PDI** | **Shell 2 PDI** | $\chi^2$ |
|---|---|---|---|---|---|---|---|---|---|---|---|
| DMA | 546.23 ± 4.93 | 465.13 ± 4.55 | 52.44 ± 1.65 | 34.37 ± 10.35 | 1.32 ± 0.01 | 0.12 ± 0.02 | 4.49 ± 1.08 | 0.40 ± 0.01 | 0.33 ± 0.14 | 0.40 ± 0.07 | 8.16 |
| HEA | 515.25 ± 5.74 | 482.77 ± 5.14 | 40.14 ± 1.03 | 32.29 ± 10.33 | 1.93 ± 0.02 | 0.20 ± 0.08 | 4.68 ± 1.10 | 0.40 ± 0.01 | 0.34 ± 0.13 | 0.39 ± 0.14 | 6.45 |
| NAM | 600.59 ± 7.16 | 498.14 ± 5.77 | 47.33 ± 1.88 | 42.61 ± 10.16 | 1.74 ± 0.02 | 0.12 ± 0.03 | 4.21 ± 1.07 | 0.40 ± 0.004 | 0.37 ± 0.11 | 0.39 ± 0.05 | 6.63 |
| PEG | 482.08 ± 3.01 | 257.32 ± 2.67 | 40.83 ± 1.22 | 37.80 ± 10.42 | 1.34 ± 0.01 | 0.12 ± 0.001 | 4.14 ± 1.14 | 0.40 ± 0.001 | 0.40 ± 0.01 | 0.40 ± 0.005 | 23.61 |

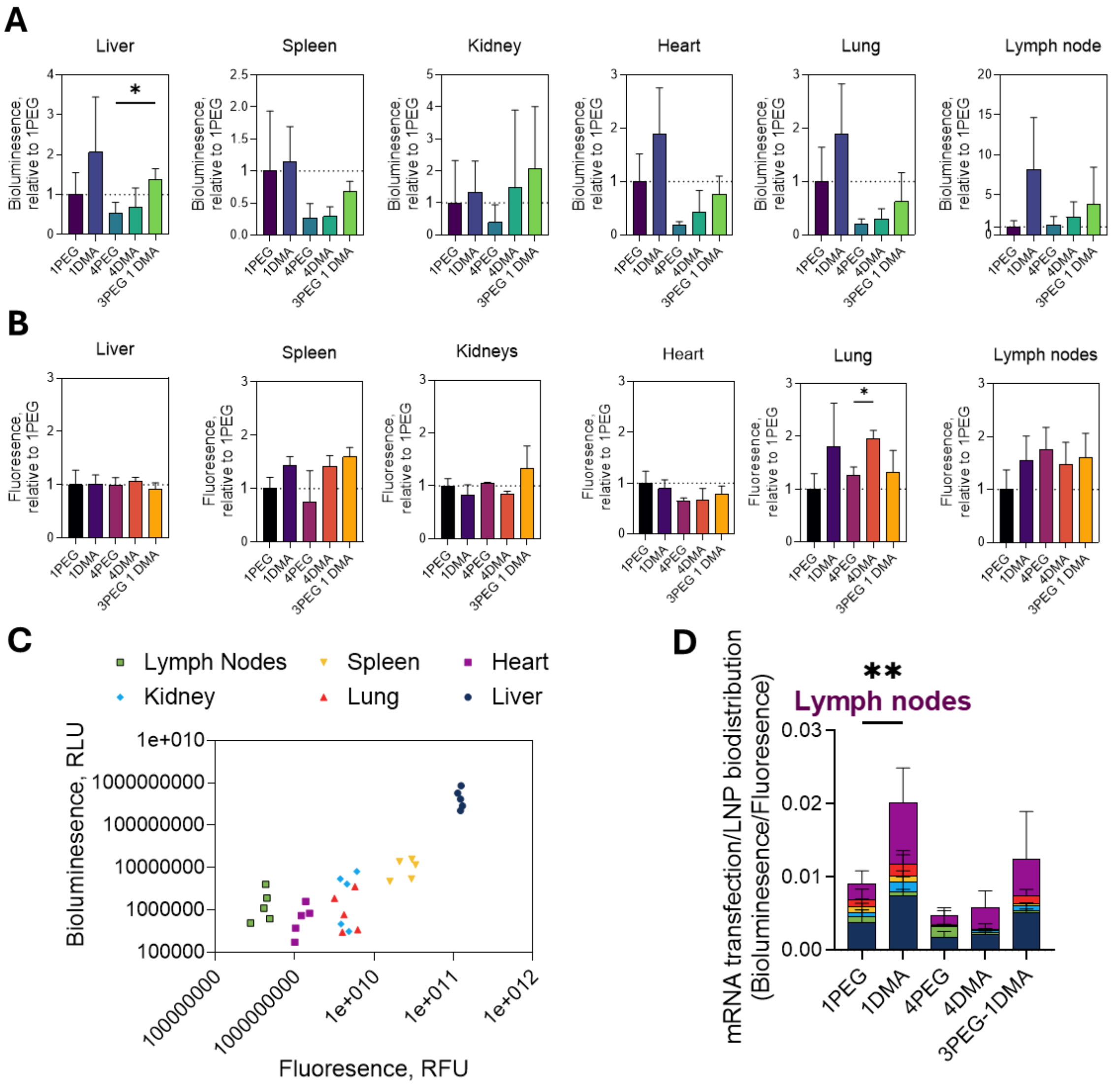

**Figure S6**. **Accompanies figure 5. PAM-lipids effectively replace PEG-lipids during repeat dosing of mRNA-LNPs** Bioluminescence (**A**) and fluorescence (**B**) of FLuc mRNA LNPs (5 ug of mRNA per dose) prepared with PAM-lipid $DMA_{20}$-H or PEG-lipid $PEG_{2k}$. Data presented for each organ as value relative to a single dose of PEG-LNP (1PEG condition). Organs and blood collected 24 h after final dose (either 1 dose or 4 doses) and imaged using in vivo imaging system (IVIS) to quantify bioluminescence (**A**) and fluorescence (**B**) from FLuc mRNA expression and fluorescently tagged LNP (DiR) respectively. Data presented as mean of biological repeats ± s.e.m. (n=3). Statistical significance was determined by one way ANOVA (Welch) with Dunnett correction compared against either 1PEG (1DMA) or 4PEG (4DMA, 3PEG-1DMA) condition, *P < 0.05. (**C**) shows corresponding bioluminescence and fluorescence values for each formulation, as the mean of 3 biological repeats. (**D**) shows mRNA transfection (bioluminescence) normalised against organ accumulation (fluorescence) of mRNA-LNPs. Data presented as mean of biological repeats (n=3). Statistical significance determined by two way ANOVA with Dunnett correction, compared against either 1PEG (1DMA) or 4PEG (4DMA, 3PEG-1DMA) condition, *P < 0.05, **P<0.01.

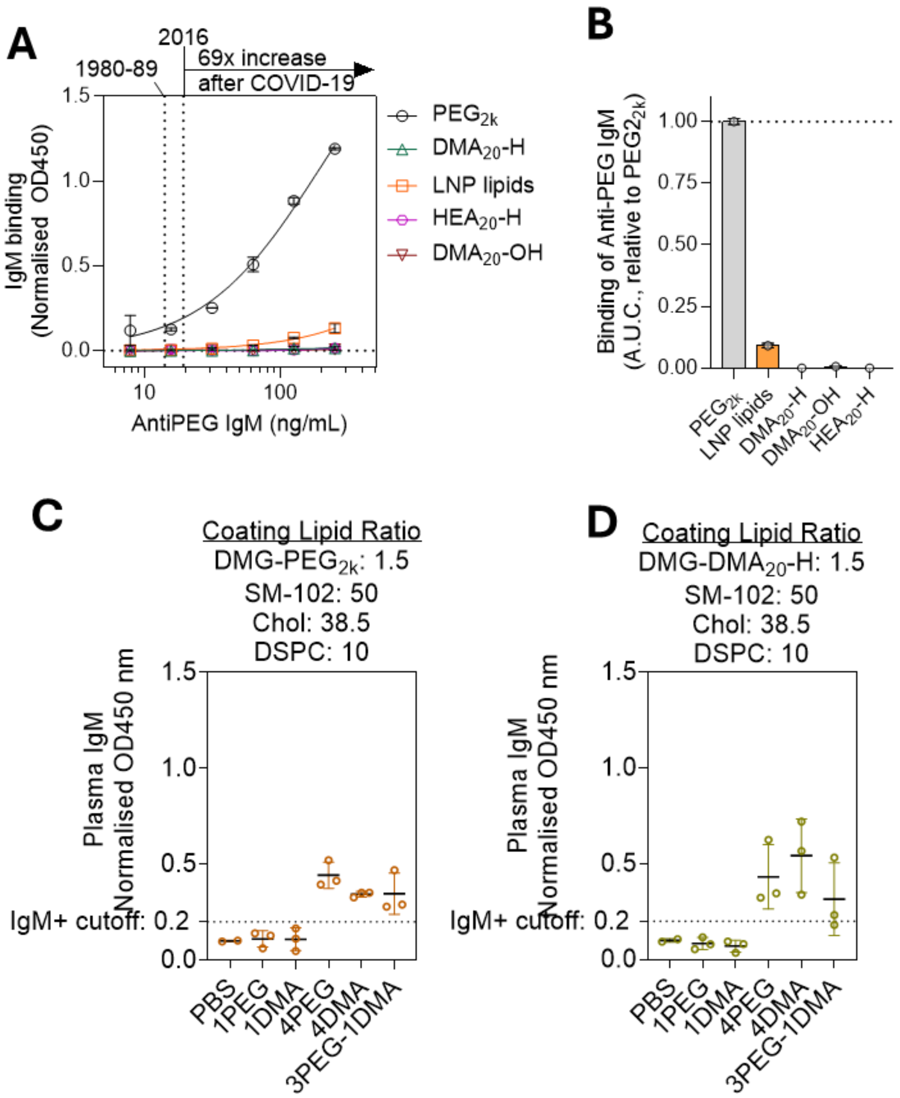

**Figure S7 – Accompanies figure 5 A**) Binding of commercially available murine anti-PEG IgM to $PEG_{2k}$, $DMA_{20}$-H, $HEA_{20}$-H and $DMA_{20}$-OH or non-polymer LNP components SM102, Cholesterol, and DSPC mixed in 50:38.5:10 molar ratios. Dotted lines represent clinically determined anti-PEG IgM levels from specified years before COVID-19, and fold-increase after COVID-19 vaccination[2,3]. **B)** Total IgM binding of commercially available anti-PEG IgM determined by area under curve (AUC) integration analysis of figure, normalised to $PEG_{2k}$. **C)** and **D**) Plasma IgM detected via enzyme-linked immunosorption assay (ELISA) of plates coated with specified polymer-lipids/lipids (LNP components). Data was normalised to OD450 nm of PBS control, set to OD450=0.1. $IgM^+$ Cutoff set as 2x OD450nm of PBS control. Data presented as mean of biological repeats ± S.D. (n=3)

##### 1.1.6 Structures of PEG, PAMs, and other common PEG alternatives

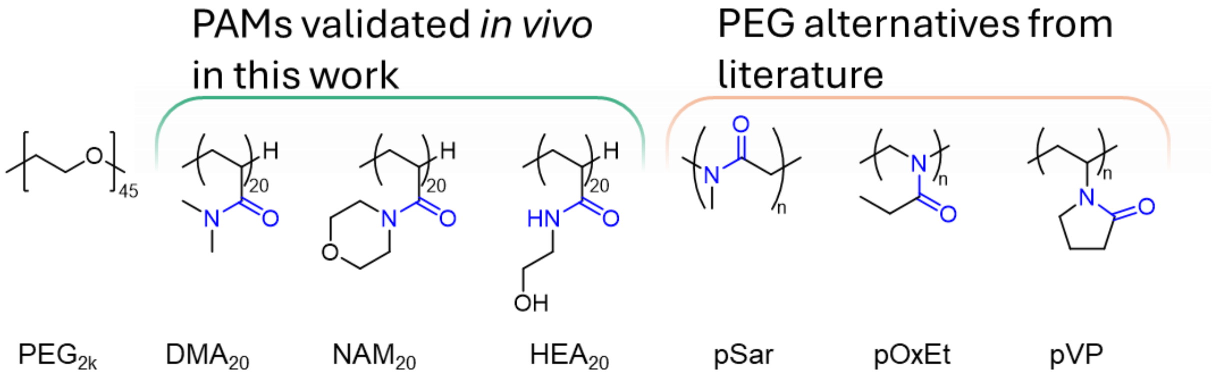

**Figure S8 –** Structures of PEG, poly(acrylamido) (PAM) polymers, and PEG alternative polymers from literature. Substituted amide groups highlighted in blue.

#### 1.2 Supporting materials and methods

##### 1.2.1 Materials

**Sigma-Aldrich (Merck)**: All solvents, solketal, 4-dimethylaminopyridine (DMAP), EDC·HCl (1-ethyl-3-(3-dimethylaminopropyl)carbodiimide hydrochloride), myristic acid (tetradecanoic acid), ammonium bromide ($NH_4Br$), sodium sulfate ($Na_2SO_4$), hydrogen peroxide (30% w/w aqueous solution), 1-ethylpiperidine hypophosphite (EPHP), deuterated NMR solvents Silica for column chromatography and silica gel 60 TLC aluminium sheets (4 cm × 8 cm) with fluorescent indicator (254 nm) for TLC. **FUJIFILM Wako Chemicals:** VA-044 initiator (2,2′-azobis[2-(2-imidazolin-2-yl)propane] dihydrochloride). **Thermo Fisher Scientific:** Amberlyst™ 15 hydrogen form ($H^+$)

##### 1.2.2 Chemical synthesis

A DMG-based macro-CTA DMG-PABTC DMG-PABTC was synthesised via a new route adapted from other methods[4,5]

##### 1.2.3 Synthesis of Solketal-PABTC

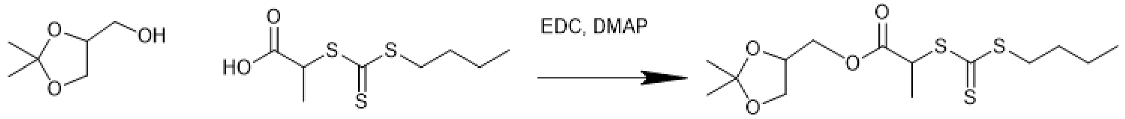

**Schematic S1: Synthesis of solketal-PABTC**

2-((butylthio)-carbonothioyl) thio propanoic acid (PABTC) was synthesized via previously reported methods[6]. PABTC (1 g, 4.02 mmol ) and Solketal (542 mg, 576 mL, 4.1 mmol) were dissolved in 10 mL dichloromethane (DCM) and cooled to 0 °C in a round bottom flask equipped with a magnetic stirrer bar. A solution of DMAP (219 mg, 1.8 mmol) dissolved in 5 mL of DCM was added slowly with stirring, and left for 5 minutes. A solution of EDC-HCl (886 mg, 4.62 mmol) in DCM (5 mL) was added dropwise with stirring and left to react overnight, allowing the ice bath melt and reach room temperature. A deep red colour was first observed which returned to bright yellow/orange overnight. Reaction progress was monitored to satisfactory completion with thin layer chromatography (TLC) using ethyl acetate (0-40%) in hexane . The resulting reaction mixture was diluted with ~25 mL of chloroform then washed twice with roughly equal volumes of $NH_4Br$ (0.5 M), then washed twice with roughly equal volumes of brine, then dried over $Na_2SO_4$. The product was purified via column chromatography (0-15% Ethyl acetate in hexane, gradient elution). The product fractions were concentrated via rotary evaporation to a viscous yellow oil and characterised by $^1$H NMR.

#### 1.2.4 Synthesis of Glycerol-PABTC

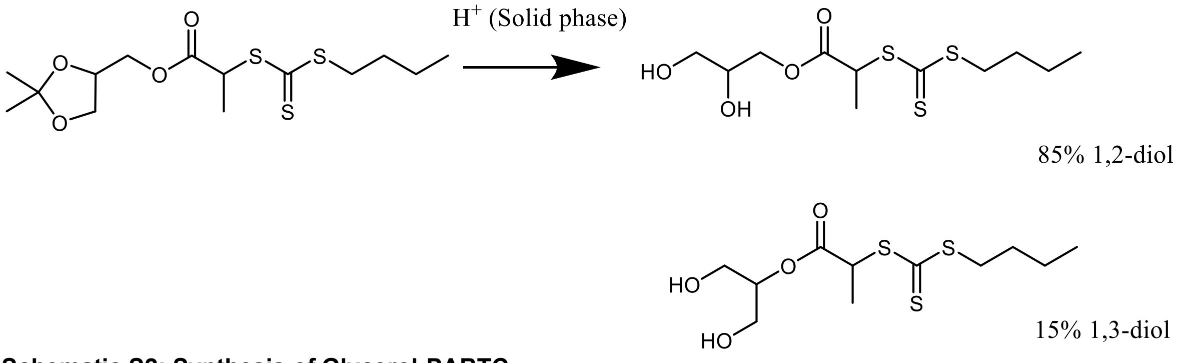

**Schematic S2: Synthesis of Glycerol-PABTC**

Method adapted from literature[7]. H+ amberlyst resin (300 mg) was added to a round bottom flask equipped with magnetic stirrer bar. Solketal-PABTC (500 mg) was dissolved in 15 mL methanol and added the flask. The solution was set heated to 60 °C and was monitored to satisfactory completion (~20 hours) via TLC (0-40% ethyl acetate in hexane). The product was separated from the amberlyst with filtration paper and the crude reaction mixture was then purified via column chromatography (0-40% Ethyl acetate in hexane, gradient elution). The product fractions were concentrated via rotary evaporation to a viscous yellow oil and characterised by $^1$H NMR.

At this stage it was determined *via* $^1$H NMR that ~15% of the product was the 1,3-diol isomer, rather than the 1,2-diol isomer (Schematic S2), which was consistent between batches. This has been seen in other reports of acid-mediated deprotection of solketal to glycerol-like molecules[4,8]. The is due to isomerisation during this reaction step. Regardless, the mixed product of 1,2- and 1,3- glycerol-PABTC was deemed acceptable and used without isomeric purification due to: 1) the potential difficulty of separating this isomers from each-other; and 2) we assumed this would have little impact on final properties of polymer-lipid; and 3) commercially available DMG-$PEG_{2k}$ is ~3% 1,3- isomer (Avanti Research, DMG-PEG 2000, product no. 80151). Moreover, no evidence of the 1,3 isomer was seen in the final product (DMG-PABTC) – potentially due to further purification.

#### 1.2.5 Synthesis of lipid RAFT agent (CTA) DMG-PABTC

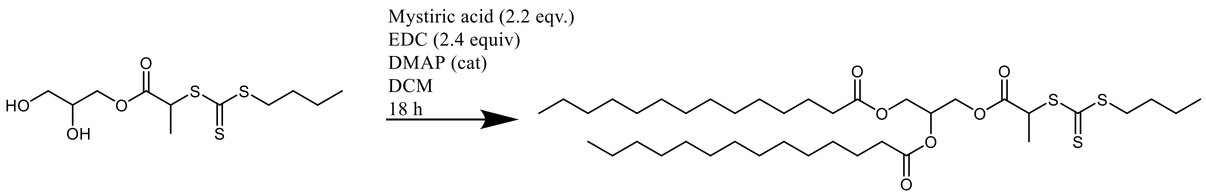

**Schematic S3: Synthesis of DMG-PABTC**

Glycerol-PABTC (1.6 g, 5.31 mmol) mystiric acid (tetradecanoic acid; 2.67 g, 11.7 mmol) were dissolved in 100 mL DCM, added to a round bottom flask with a magnetic stirrer and cooled to 0 °C with an ice bath. DMAP (525 mg, 4.3 mmol, 122.17 g/mol) was dissolved in 5 mL of DCM and added slowly to the reaction mixture with stirring and left for 5 minutes. A solution of EDC-HCl (2.00 g, 12.9 mmol) in DCM (20 mL) was add dropwise with stirring, and left to react overnight, allowing the ice bath left to melt and reach room temperature. A deep red colour was first observed which returned to bright yellow/orange overnight. Reaction progress was monitored to satisfactory completion with thin layer chromatography (TLC) using ethyl acetate (0-40%) in hexane. The resulting reaction mixture was diluted with ~50 mL of chloroform then washed twice with roughly equal volumes of $NH_4Br$ (0.5 M), then washed twice with roughly equal volumes of brine, then dried over $Na_2SO_4$. The product was purified via column chromatography (0-15% Ethyl acetate in hexane, gradient elution). The product fractions were concentrated via rotary evaporation to a viscous yellow oil which solidified upon storage at -20 °C, and was characterised by $^1H$ NMR and $^{13}C$ NMR.

#### 1.2.6 Polymer synthesis including end group removal

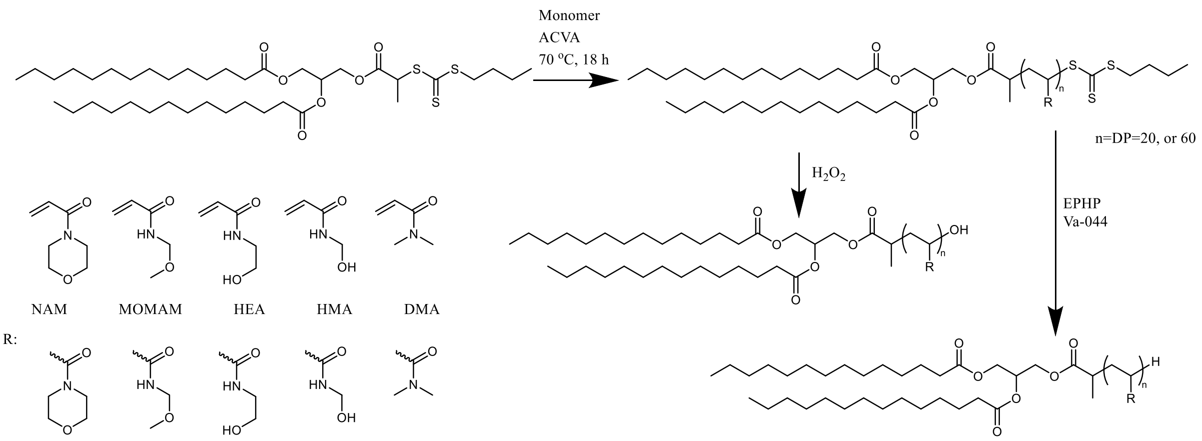

**Schematic S4: Synthesis of poly(acrylamido) lipids via RAFT polymerisation using DMG-CTA, followed by end group removal**

Synthesis of polymers (with $-S_3C_5H_9$ end group) is outlined in materials and methods, with specific conditions and characterisation shown in table 1. All polymers were prepared using the macro-CTA DMG-PABTC (synthesis described above). The monomers used for each polymer share the abbreviation used for the polymer I.D., e.g. $NAM_{20}$-$S_3C_5H_9$ was synthesised using n-acryloyl morpholine (NAM). Polymers with their end groups replaced were synthesised by taking a portion of the polymer with the $-S_3C_5H_9$ end group and subjecting them to the reaction conditions described below, to yield either -OH or -H end groups. All polymers were characterised by $^1H$ NMR and gel permeation chromatography (GPC) analysis.

#### 1.2.7 End-group replacement with -OH

End-group replacement with -OH group was achieved via methods modified from those previously described[9,10]. Briefly, 18 umol of PAM-lipid polymer with the $-S_3C_5H_9$ end group was dissolved to 100 mg/mL in a solvent mix with volumetric ratios of either: $H_2O$:IPA 2:1, $H_2O$:IPA 1:1, or $H_2O$:Dioxane:IPA 1:1:1 depending on polymer solubility. 180 umol of $H_2O_2$ was added from a 30% stock solution in water. The solution was added to a vial equipped with a magnetic stirrer bar stirred at 70 °C for 24 hours without degassing. End group replacement was deemed finished after loss of bright yellow colouration. Where the reaction had not reached satisfactory completion, the same amount of $H_2O_2$ was added to the vial again and left for a further 24 hours up to two more times.

#### 1.2.8 End-group replacement with -H

End-group replacement with -H was achieved via methods modified from those previously described[11]. Briefly, polymers with $-S_3C_5H_9$ (~17 umol) and 1-ethylpiperidine hypophosphite (170 umol, 20 mg) were dissolved in 600 uL of dioxane:water 2:1, IPA:water 2:1, or dioxane:IPA:water 1:1:1. Azoinitiator 2,2'-Azobis[2-(2-imidazolin-2-yl)propane]dihydrochloride (VA-044, 1.7 umol, [CTA]/[I]=10, 0.53 mg) was added from a stock solution prepared in $H_2O$. The solution was mixed and added to a reaction vial equipped with a stirrer and rubber septa. The solution was degassed via bubbling nitrogen through for at least 15 minutes then heated to 70 °C for 2-3 hours. End group removal was deemed finished after loss of bright yellow colouration. Where the reaction had not reached satisfactory completion, the same amount of initiator and EPHP was added to the vial, followed by degassing and heating to 70 °C for 2-3 hours as before.

#### 1.2.9 Polymer characterisation

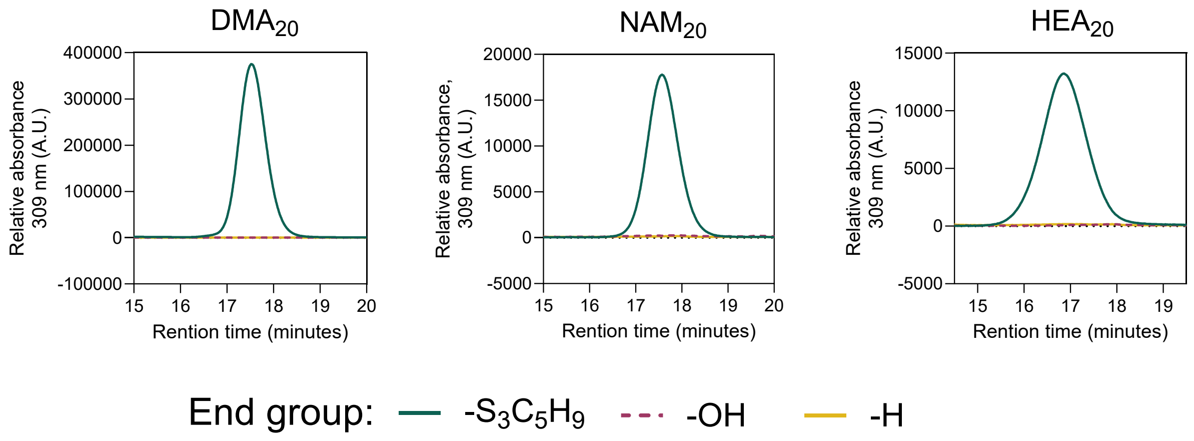

**Figure S9. confirmation of removal of $-S_3C_5H_9$ end group** Gel permeation chromatography (GPC) coupled to UV/Vis detection of PAM-lipids at characteristic absorbance for trithiocarbonyl group (309 nm) present in $-S_3C_5H_9$ end group.

*Table S5 - Synthesis conditions and characterisation of PAM-lipid polymers prepared with DMG-based macro-CTA before end group removal (-$S_3C_5H_9$) and after end group replacement with -OH, or -H. M = monomer, CTA = chain transfer agent, I = initiator, int= integration under relevant peak in $^1$H NMR. PAMs in green were validated in vivo.*

| Polymer I.D. (all prepared with DMG-CTA) | Monomer (mass, mg) | Solvent ratio (Dioxane:Water) | [M] | Target D.P., [M]/[CTA] | [M] at time=0 ($^1$H NMR) | [CTA]/[I] | Conversion, % (from $^1$H NMR) | Final D.P. at t=finished ($^1$H NMR) | $M_{n,\ th,}$ from conversion, g/mol | $M_w$ from GPC, (g/mol) | Dispersity (Mw/Mn) |
|---|---|---|---|---|---|---|---|---|---|---|---|
| $DMA_{20}$-$S_3C_5H_9$ | 250 | 10:0 | 2 | 20 | 19.2 | 20 | 97.0 % | 18.7 | 2586 | 3357 | 1.08 |
| $DMA_{20}$-OH | | | | | | | | | 2438 | 3255 | 1.12 |
| $DMA_{20}$-H | | | | | | | | | 2423 | 2839 | 1.09 |
| $DMA_{60}$-$S_3C_5H_9$ | 350 | 10:0 | 2 | 60 | 66.0 | 20 | 95.5 % | 63.0 | 6978 | 8652 | 1.05 |
| $DMA_{60}$-OH | | | | | | | | | 6830 | 9520 | 1.10 |
| $DMA_{60}$-H | | | | | | | | | 6814 | 8712 | 1.07 |
| $NAM_{20}$-$S_3C_5H_9$ | 250 | 10:0 | 2 | 20 | 25.34 | 20 | 98.7 % | 25 | 4262 | 3363 | 1.07 |
| $NAM_{20}$-OH | | | | | | | | | 4114 | 3732 | 1.09 |
| $NAM_{20}$-H | | | | | | | | | 4098 | 3517 | 1.06 |
| $NAM_{60}$-$S_3C_5H_9$ | 350 | 10:0 | 2 | 60 | 55.7 | 20 | 99.0% | 55.1 | 8497 | 7249 | 1.20 |
| $NAM_{60}$-OH | | | | | | | | | 8349 | 7765 | 1.24 |
| $NAM_{60}$-H | | | | | | | | | 8333 | 8045 | 1.11 |
| $MOMAM_{20}$-$S_3C_5H_9$ | 250 | 10:0 | 1.5 | 20 | 19.3 | 20 (2 additions) | 98.5 % | 19 | 2920 | 4825 | 1.07 |
| $MOMAM_{20}$-OH | | | | | | | | | 2772 | 5691 | 1.13 |
| $MOMAM_{20}$-H | | | | | | | | | 2756 | 5309 | 1.17 |
| $MOMAM_{60}$-$S_3C_5H_9$ | 350 | 10:0 | 1.5 | 60 | - | 20 (3 additions) | - | - | - | 22253 | 1.34 |
| $MOMAM_{60}$-OH | | | | | | | | | - | 29119 | 1.31 |
| $MOMAM_{60}$-H | | | | | | | | | - | 24612 | 1.43 |
| $HEA_{20}$-$S_3C_5H_9$ | 250 | 8:2 | 2 | 20 | 17.5 | 20 | 95.4 % | 16.7 | 2656 | 6551 | 1.18 |
| I | | | | | | | | | 2508 | - | - |
| $HEA_{20}$-H | | | | | | | | | 2492 | - | - |
| $HEA_{60}$-$S_3C_5H_9$ | 350 | 8:2 | 2 | 60 | 65 | 20 | 98.5% | 64 | 8101 | 10363 | 1.32 |
| $HEA_{60}$-OH | | | | | | | | | 7953 | 6817 | 1.12 |

| | | | | | | | | | | | |
|---|---|---|---|---|---|---|---|---|---|---|---|
| $HEA_{60}$-H | | | | | | | | | 7937 | 23492 | 1.30 |
| $HMA_{20}$-$S_3C_5H_9$ | 250 | 4:6 | 1 | 20 | 20.5 | 20 | 98.0 % | 20.0 | 2755 | 4964 | 1.16 |
| $HMA_{20}$-OH | | | | | | | | | 2607 | 6529 | 1.20 |
| $HMA_{20}$-H | | | | | | | | | 2591 | | |
| $HMA_{60}$-$S_3C_5H_9$ | 350 | 4:6 | 1 | 60 | 52.1 | 20 | 96.2 % | 50.0 | 5788 | 13856 | 1.07 |
| $HMA_{60}$-OH | | | | | | | | | 5640 | - | - |
| $HMA_{60}$-H | | | | | | | | | 5624 | - | - |

### 1.3 Flow cytometry gating strategies

#### 1.3.1 BMDCs

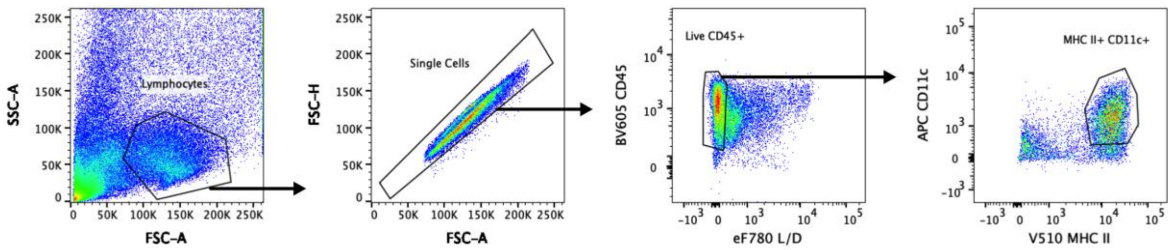

**Figure S10 – Flow cytometry gating strategy for BMDCs.**
Representative gating strategy for live BMDCs (cells, single cells, live CD45+, Lin – (CD19 NK1.1 B220 Ly6c Ly6g), MHC II+ CD11c+).

#### 1.3.2 THP-1s

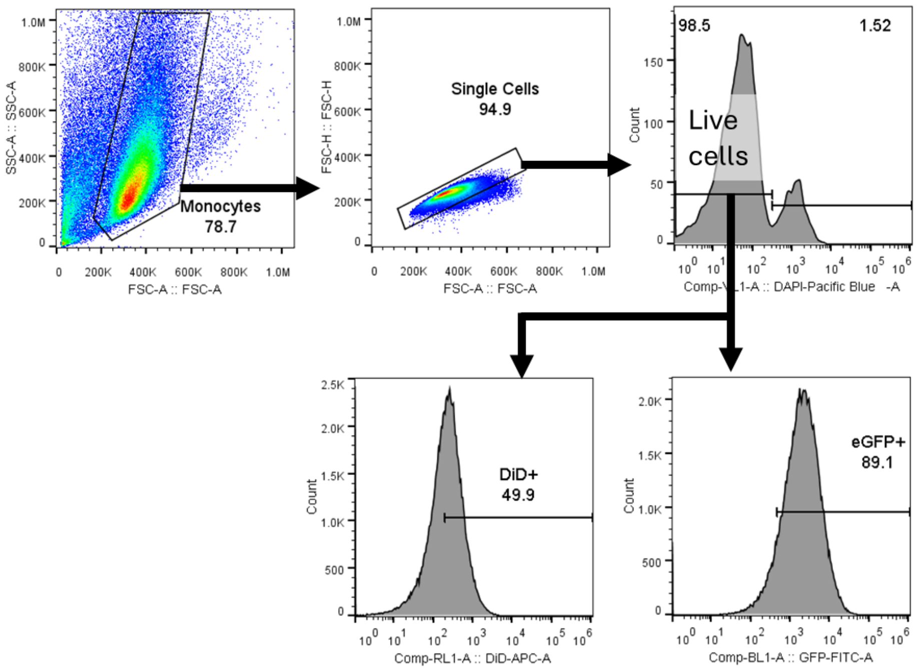

**Figure S11 – Flow cytometry gating strategy for THP-1s** Monocytes, single cells, then live cells (DAPI-negative) before analysis of DiD and eGFP fluorescence.

#### 1.3.3 In vivo

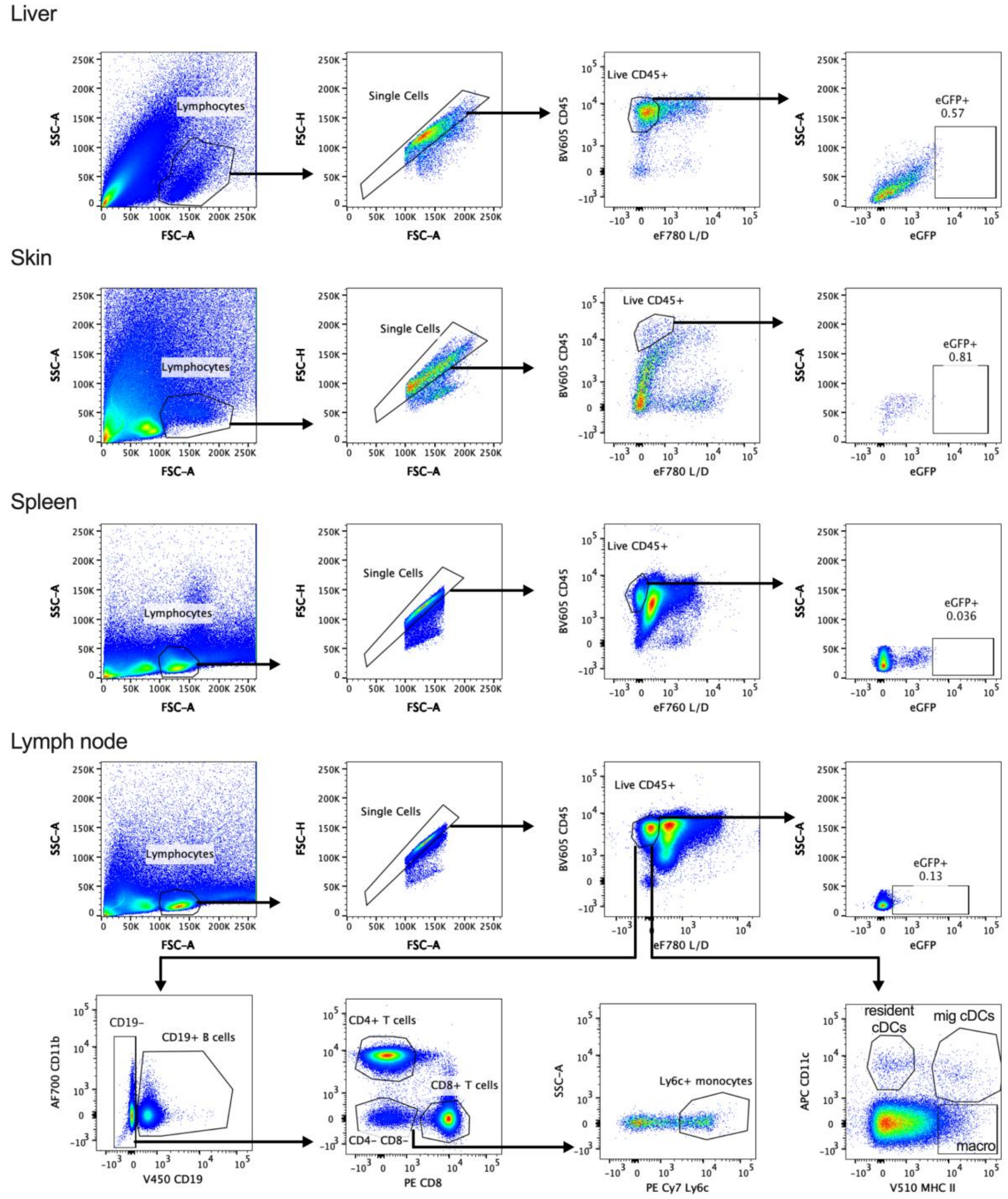

**Figure S12 – Flow cytometry gating strategy for liver, skin, spleen and lymph nodes**
Representative flow cytometry plots for gating of the total immune cell populations (live CD45+) in the liver, skin, spleen and lymoh node. Additional representative flow plots for immune cell population in dLN to identify B cells (Single, live, CD45+ CD19+), CD8+ T cells (Single, live, CD45+, CD19- CD8+), CD4+ T cells (Single, live, CD45+, CD19- CD4+), monocytes (Single, live, CD45+ CD19- CD4-, CD8-, ly6c+), macrophages (Single, live, CD45+, lin- (NK1.1 CD19, B220, CD3e, Ly6c, ly6g) MHC II- CD11c-) , resident cDCs (Single, live, CD45+, lin- (NK1.1 CD19, B220, CD3e, Ly6c, ly6g) MHC II+ $CD11c^{low}$, migratory cDCs (Single, live, CD45+, lin- (NK1.1 CD19, B220, CD3e, Ly6c, ly6g) MHC II+ $CD11c^{high}$.

### 1.4 NMR spectra

#### 1.4.1 DMG-PABTC lipid RAFT agent CTA

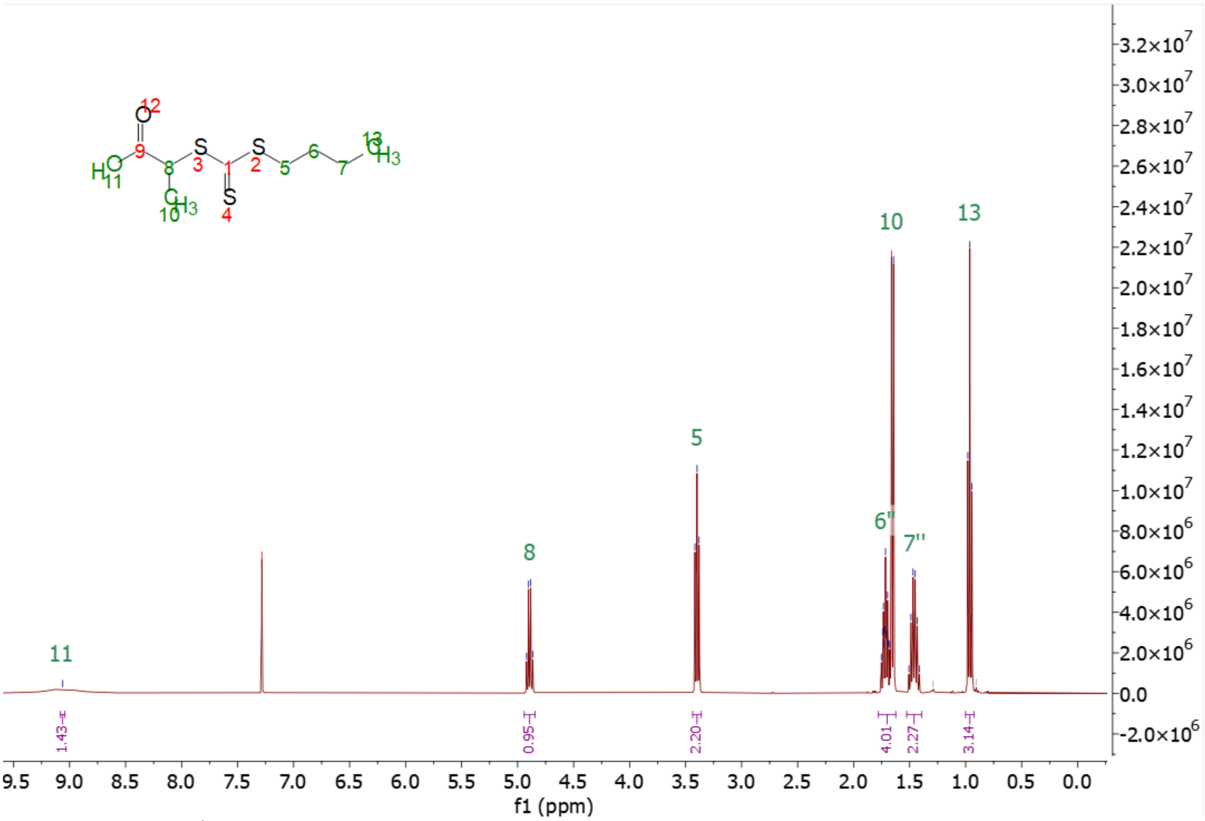

Figure S13 – [1]H NMR ($CDCl_3$) of PABTC

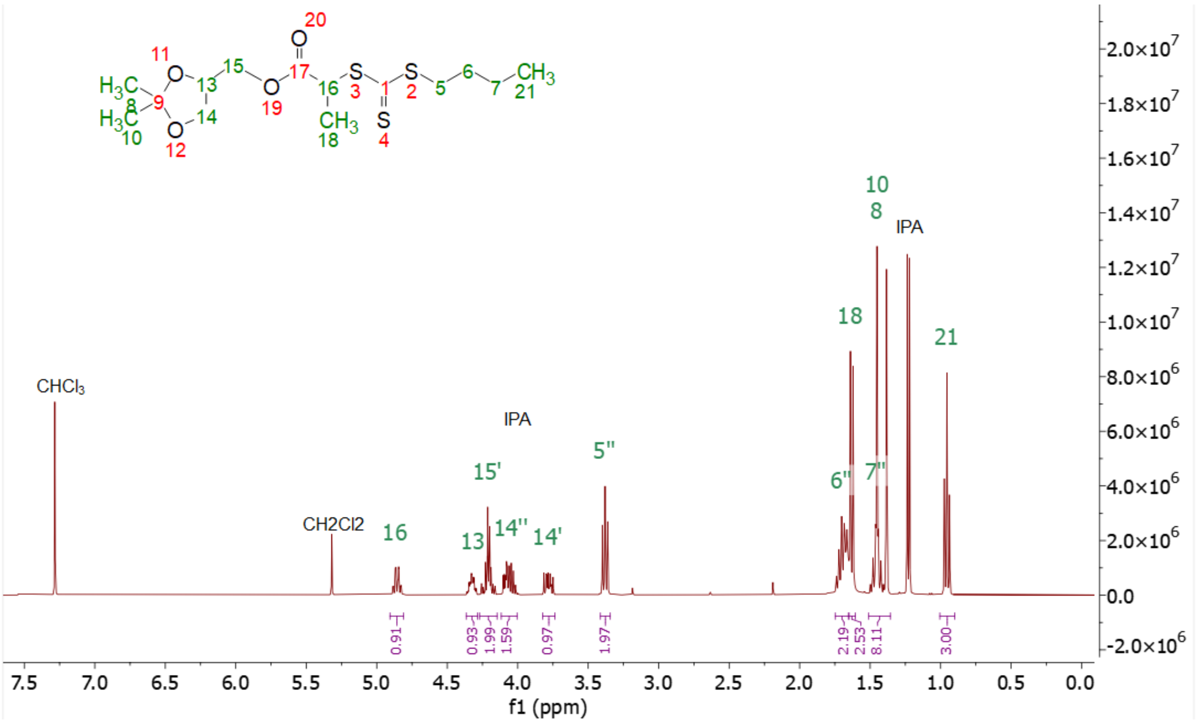

Figure S14 – [1]H NMR ($CDCl_3$) of Solketal-PABTC

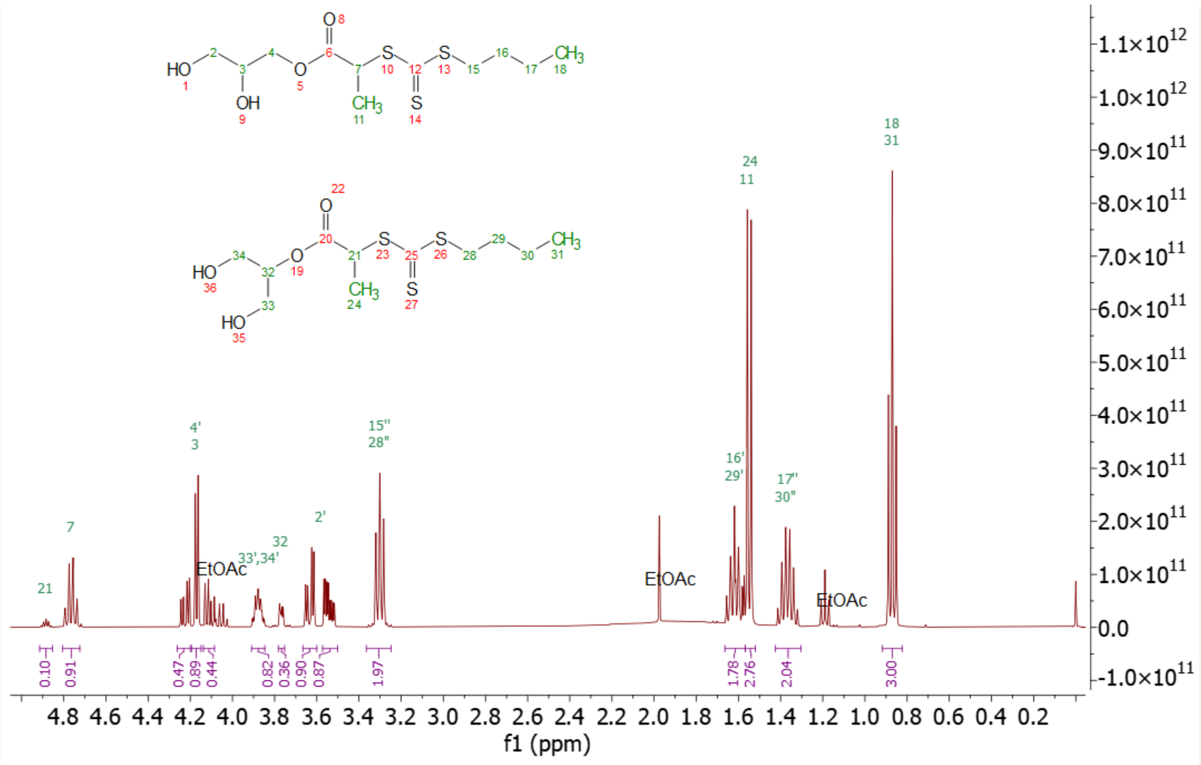

Figure S15 – $^1H$ NMR ($CDCl_3$) of Glycerol-PABTC

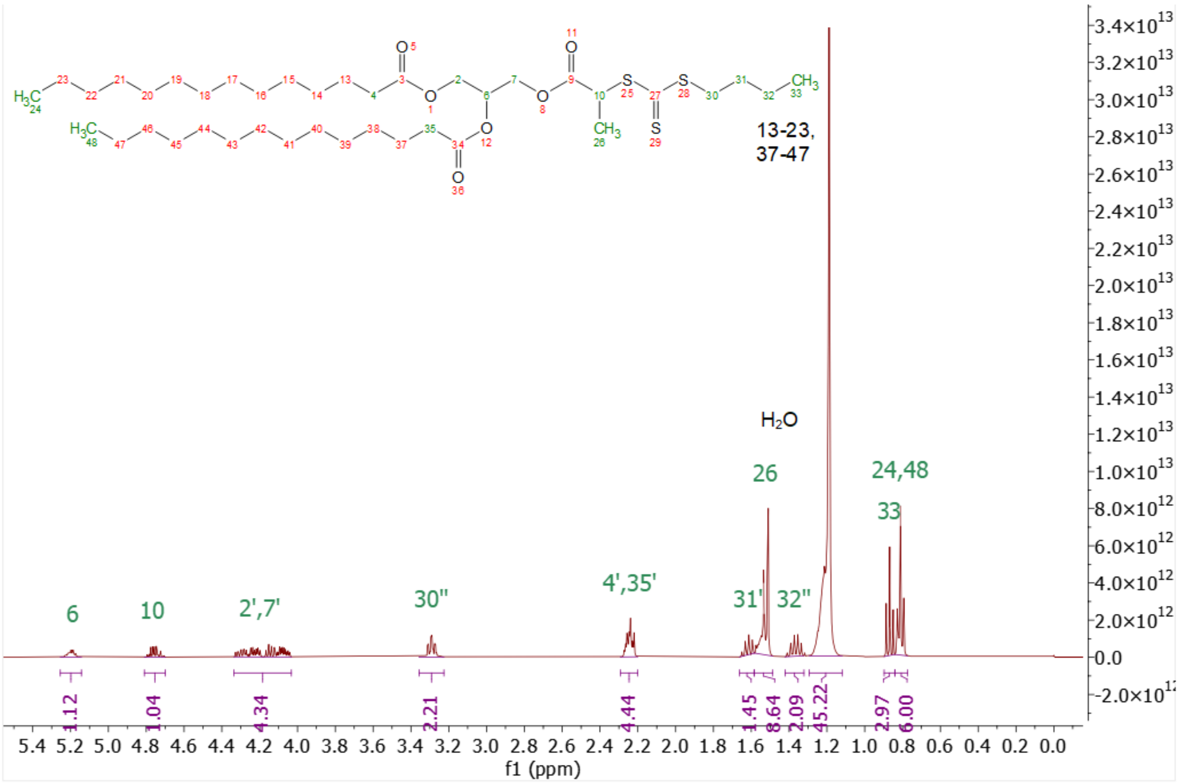

Figure S16 – $^1H$ NMR ($CDCl_3$) of DMG-PABTC

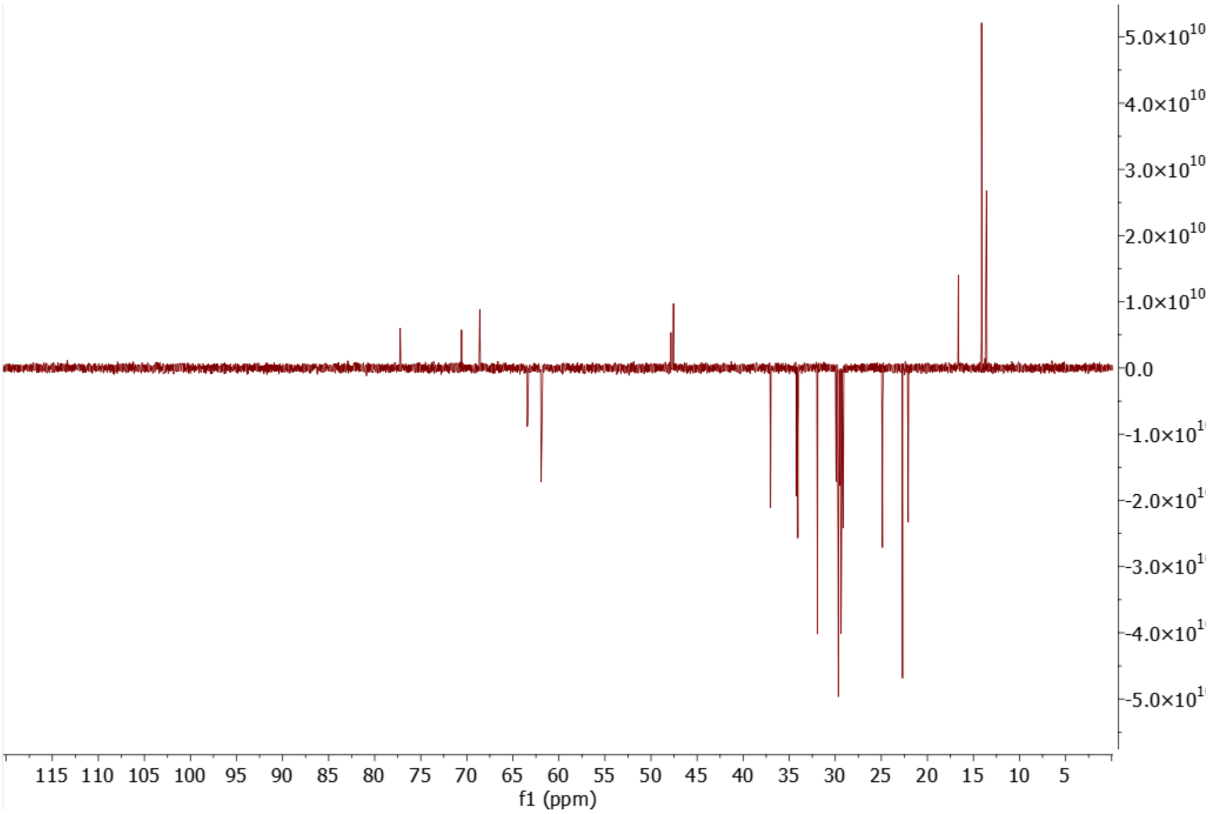

Figure S17 – $^{13}C$ NMR ($CDCl_3$) of DMG-PABTC

#### 1.4.2 PAM polymer Synthesis

##### *1.4.2.1 Lead PAM-lipids DMA$_{20}$-H, NAM$_{20}$-H, and HEA$_{20}$-H*

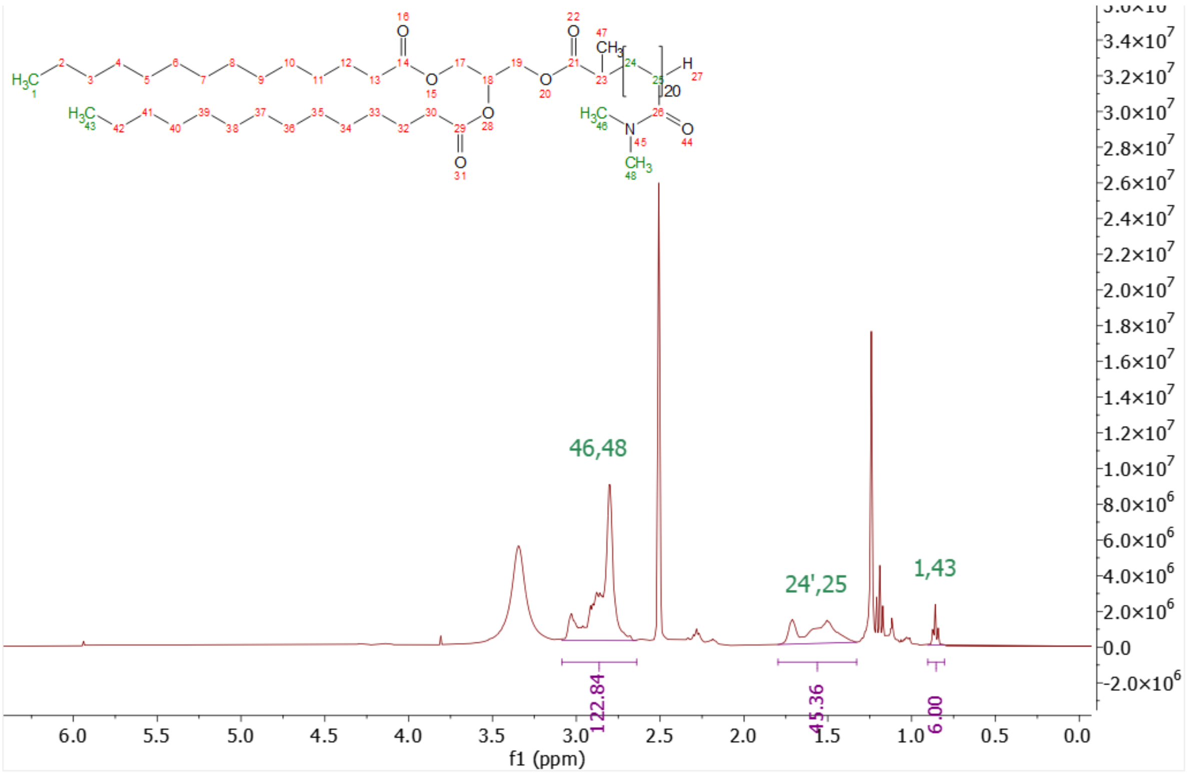

Figure S18 – $^{1}H$ NMR (d6-DMSO, 400 MHz) of DMA$_{20}$-H

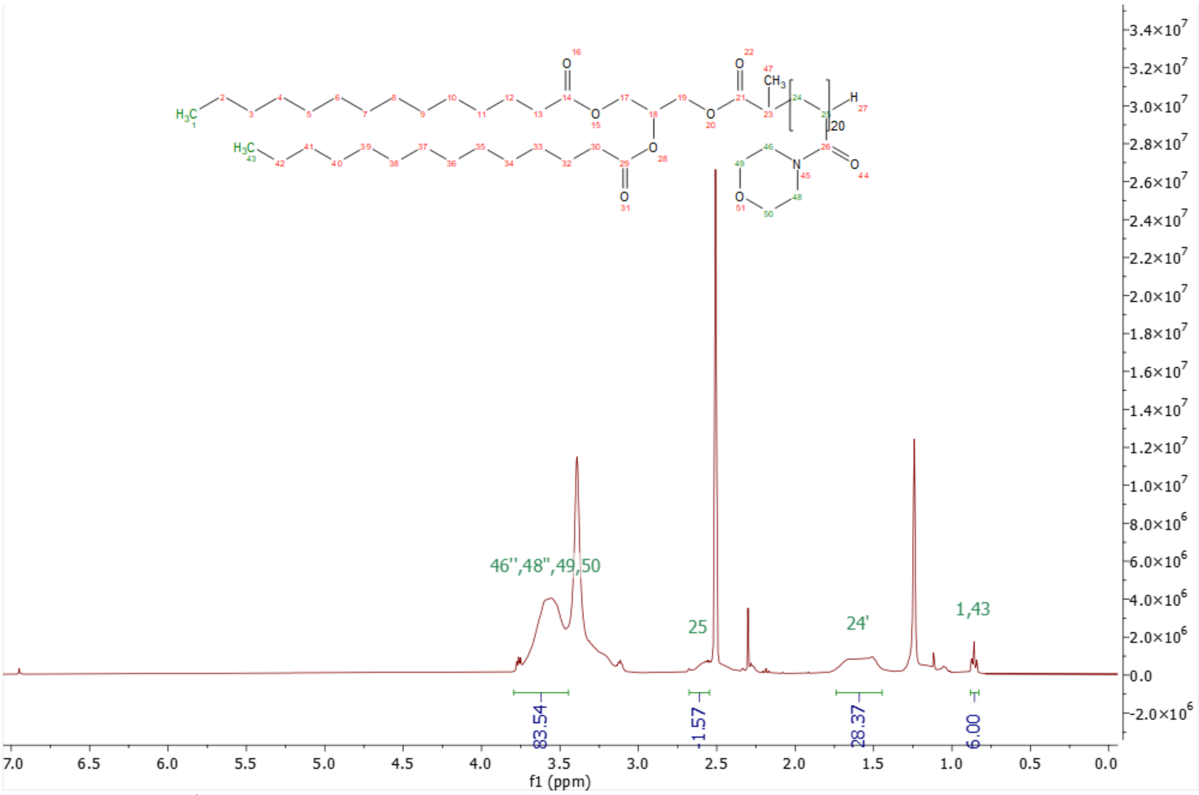

Figure S18 – [1]H NMR (d6-DMSO, 400 MHz) of $NAM_{20}$-H

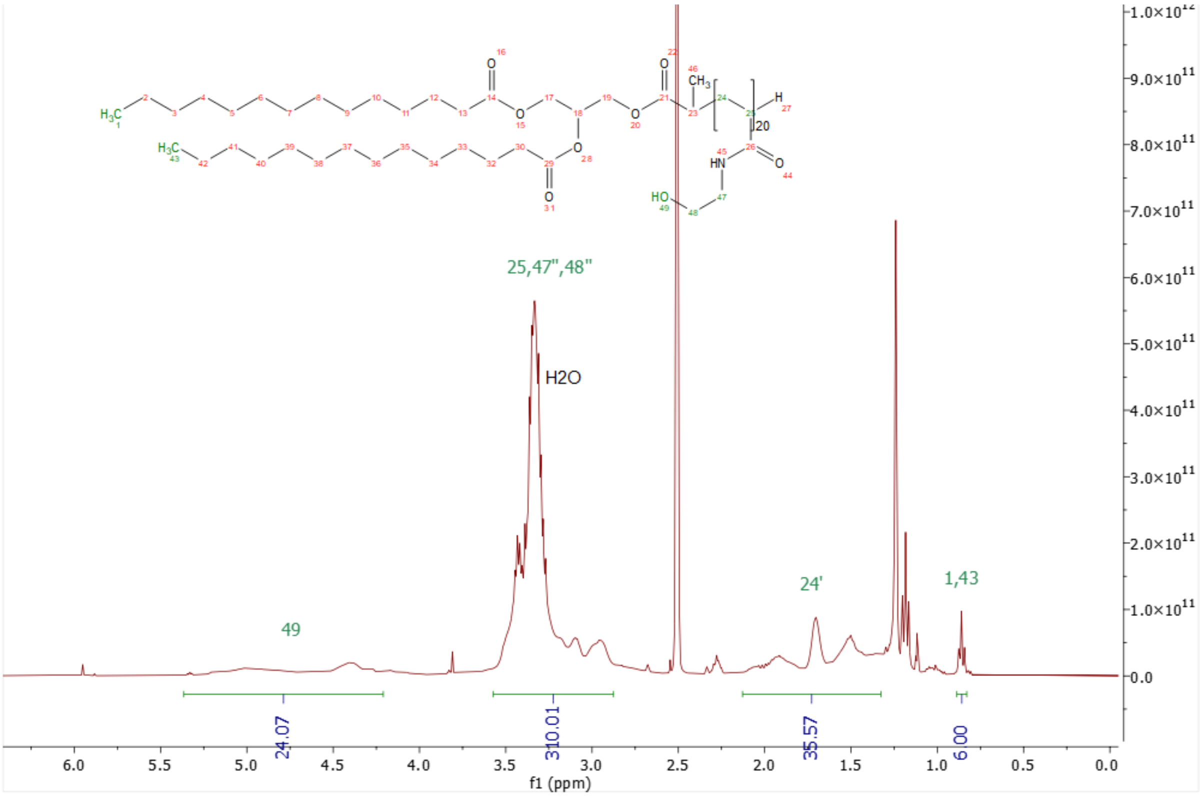

Figure S19 – [1]H NMR (d6-DMSO, 400 MHz) of $HEA_{20}$-H

##### *1.4.2.2 Remainder of 30 PAM-lipid library*

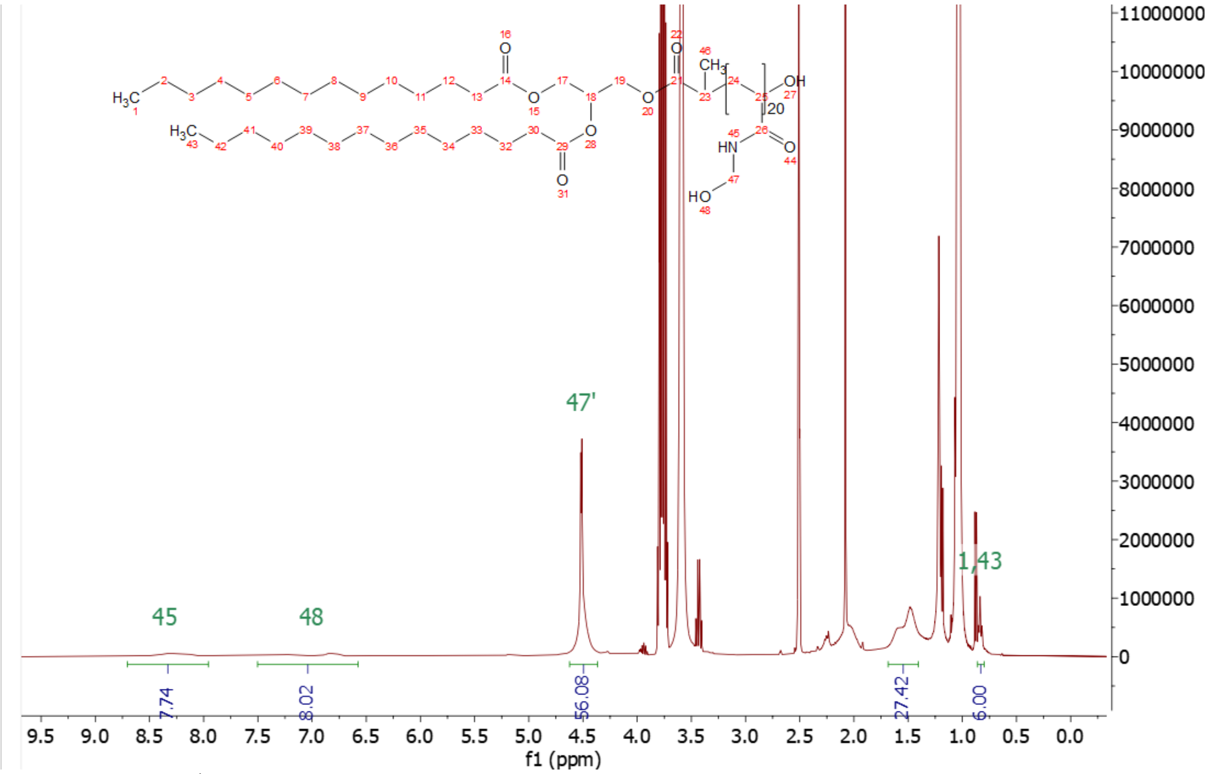

Figure S20 – [1]H NMR (d6-DMSO, 400 MHz) $HMA_{20}$-OH

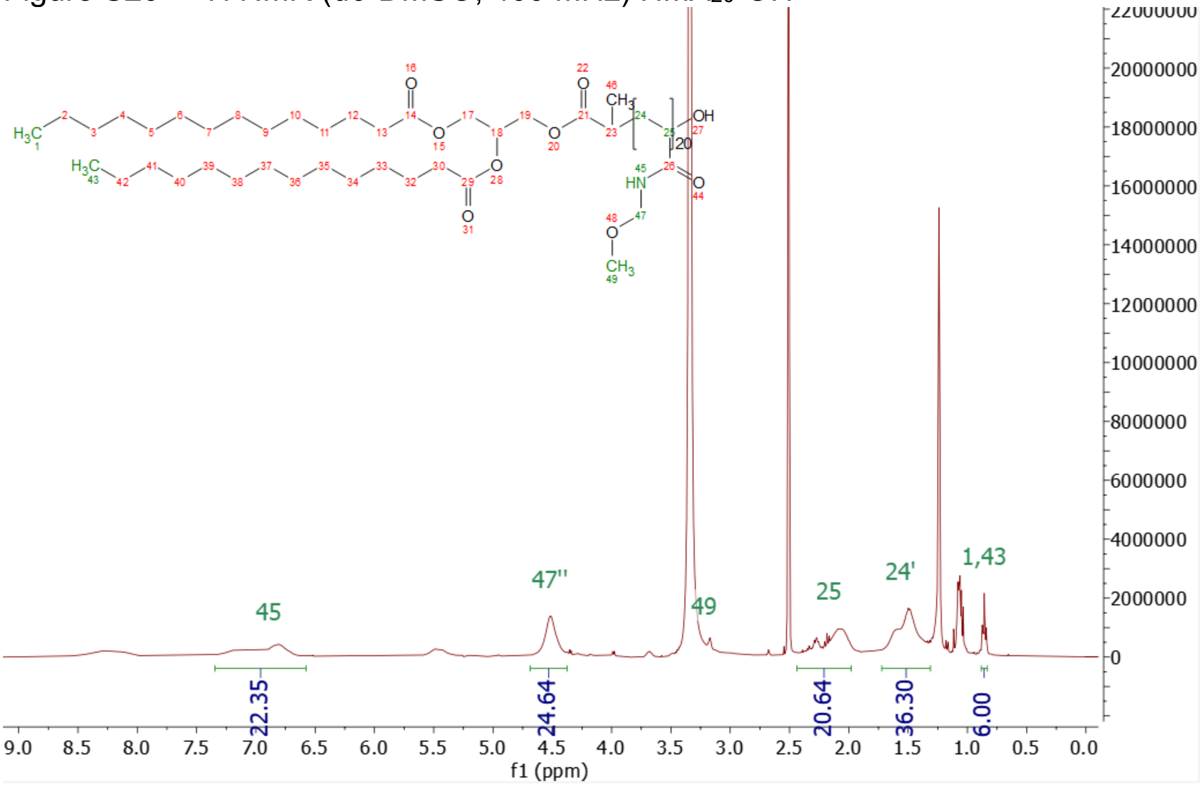

Figure S21 – [1]H NMR (d6-DMSO, 400 MHz) $MOMAM_{20}$-OH

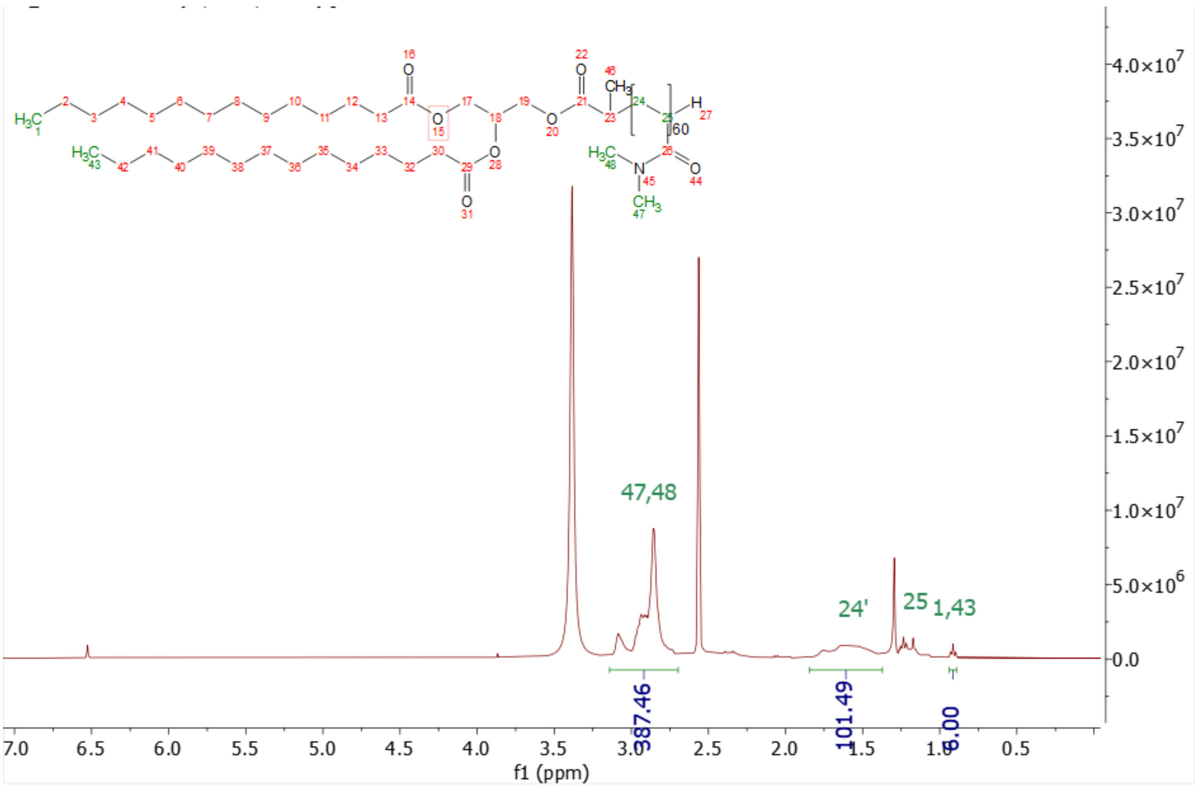

Figure S22 – [1]H NMR (d6-DMSO, 400 MHz) $DMA_{60}$-H

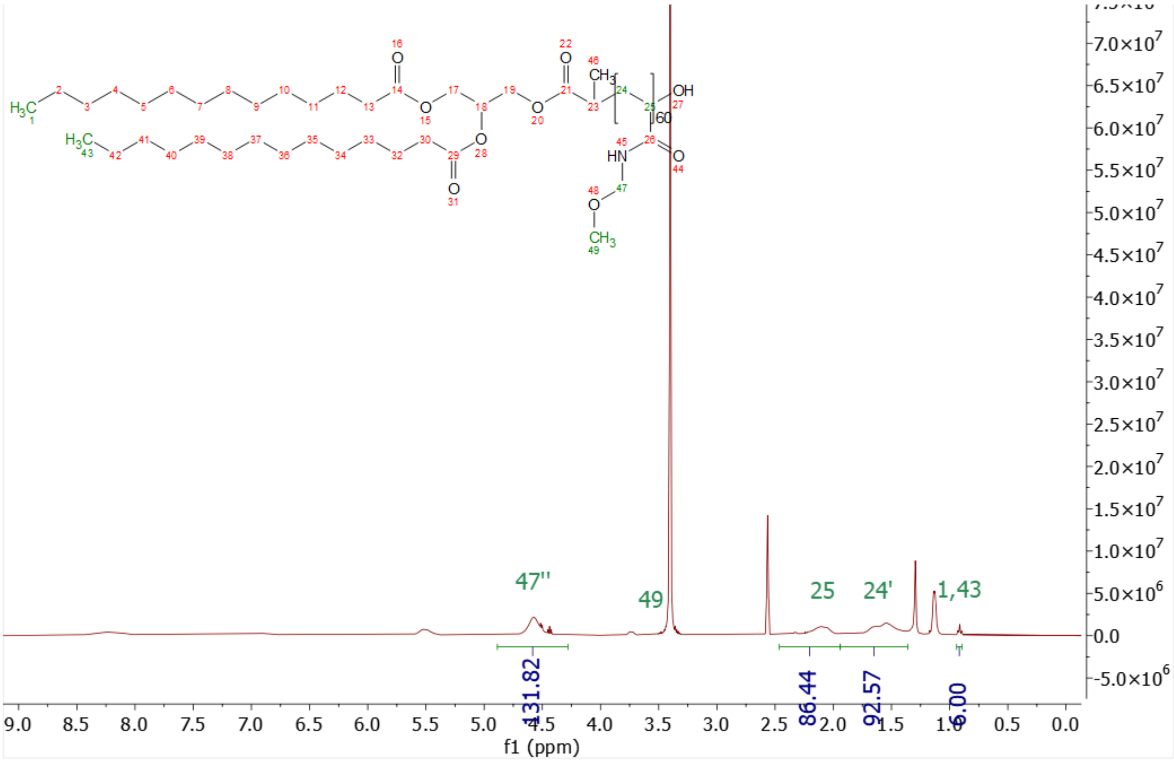

Figure S23 – [1]H NMR (d6-DMSO, 400 MHz) $MOMAM_{60}$-OH

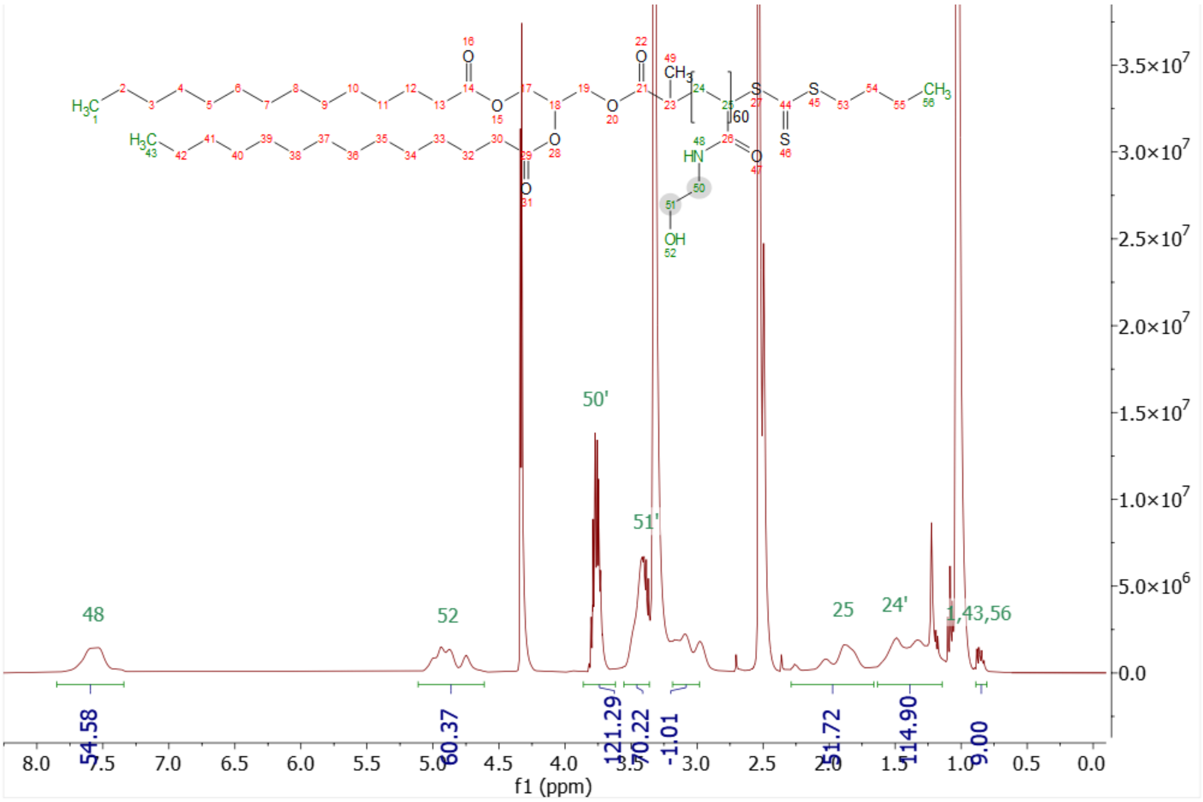

Figure S24 – [1]H NMR (d6-DMSO, 400 MHz) $HEA_{60}$-S

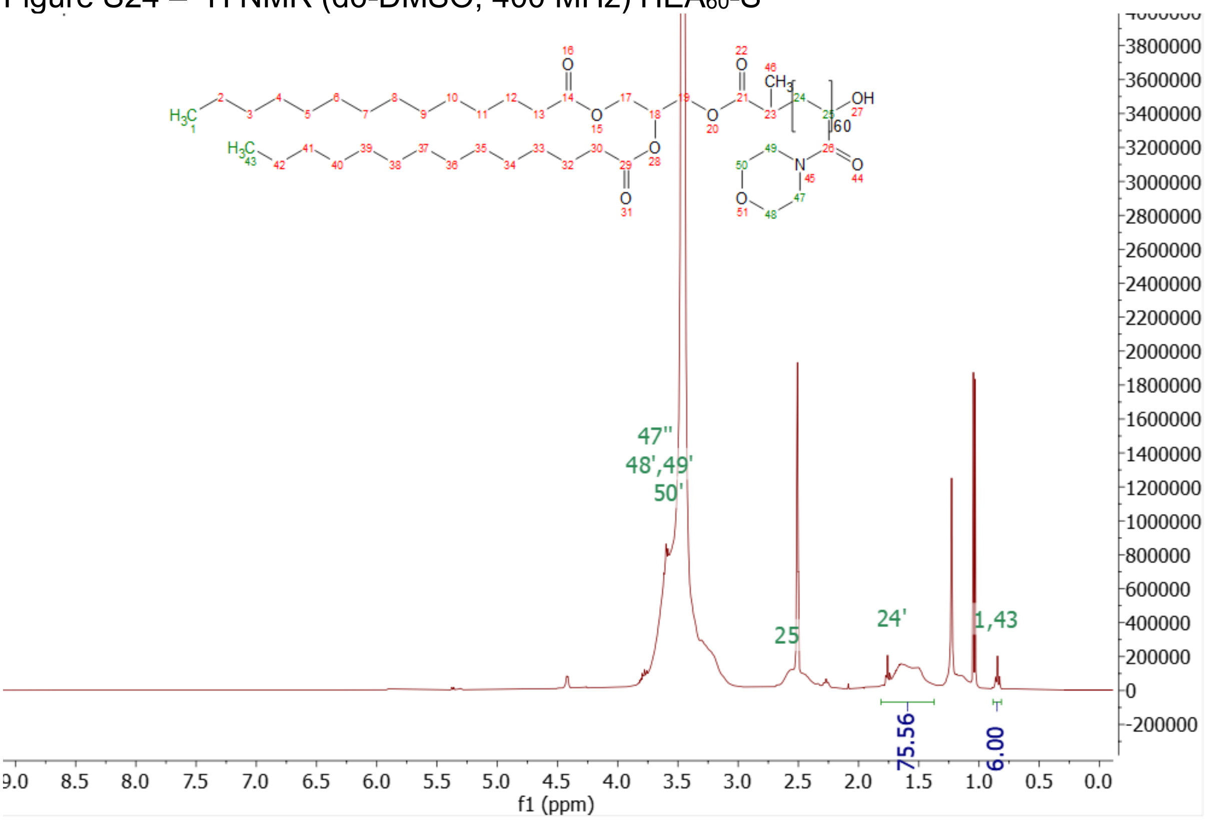

Figure S25 – [1]H NMR (d6-DMSO, 400 MHz) $NAM_{60}$-OH

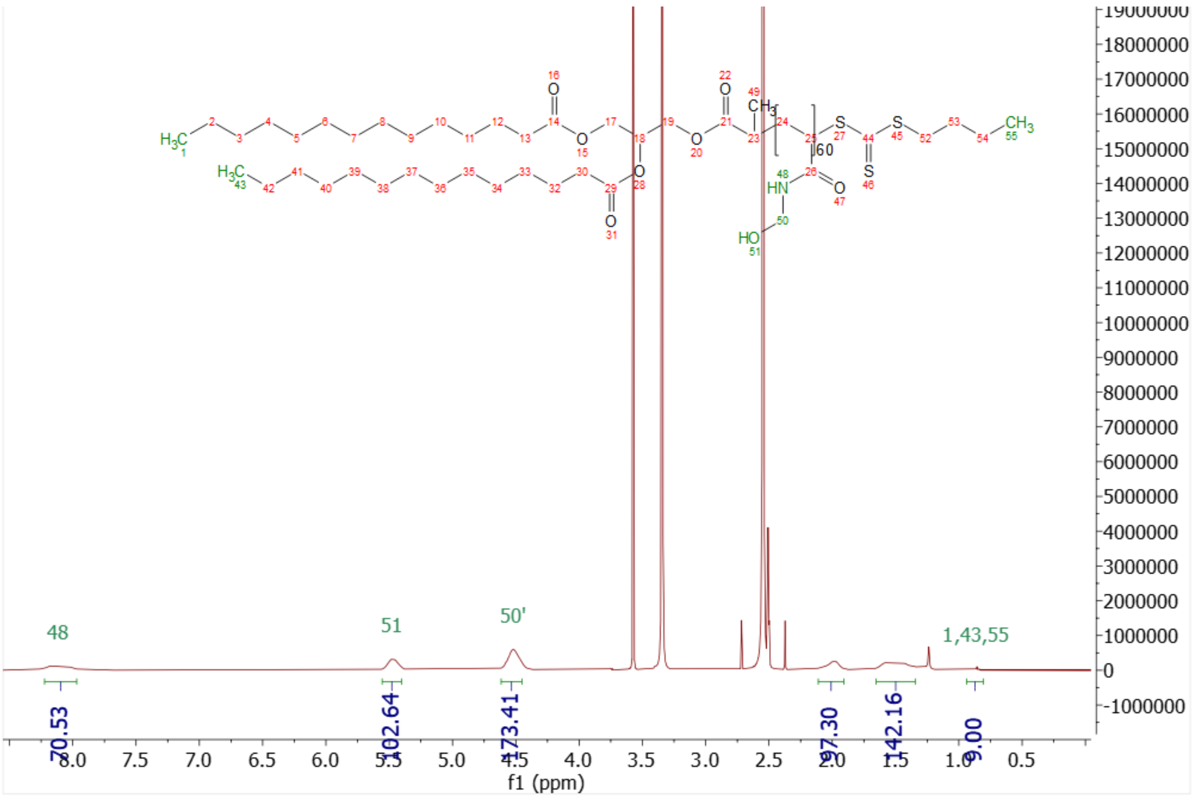

Figure S26 – $^{1}$H NMR (d6-DMSO, 400 MHz) $HMA_{60}$-S

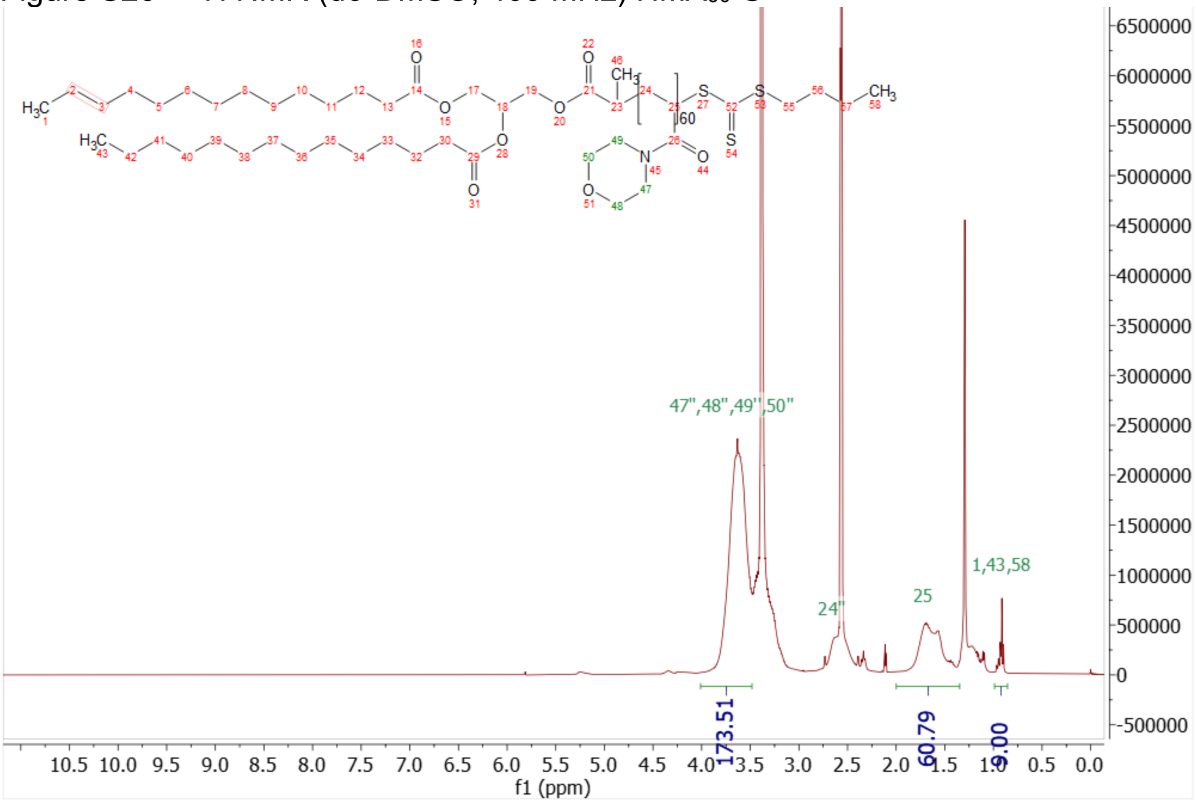

Figure S27 – $^{1}$H NMR (d6-DMSO, 400 MHz) $NAM_{60}$-S

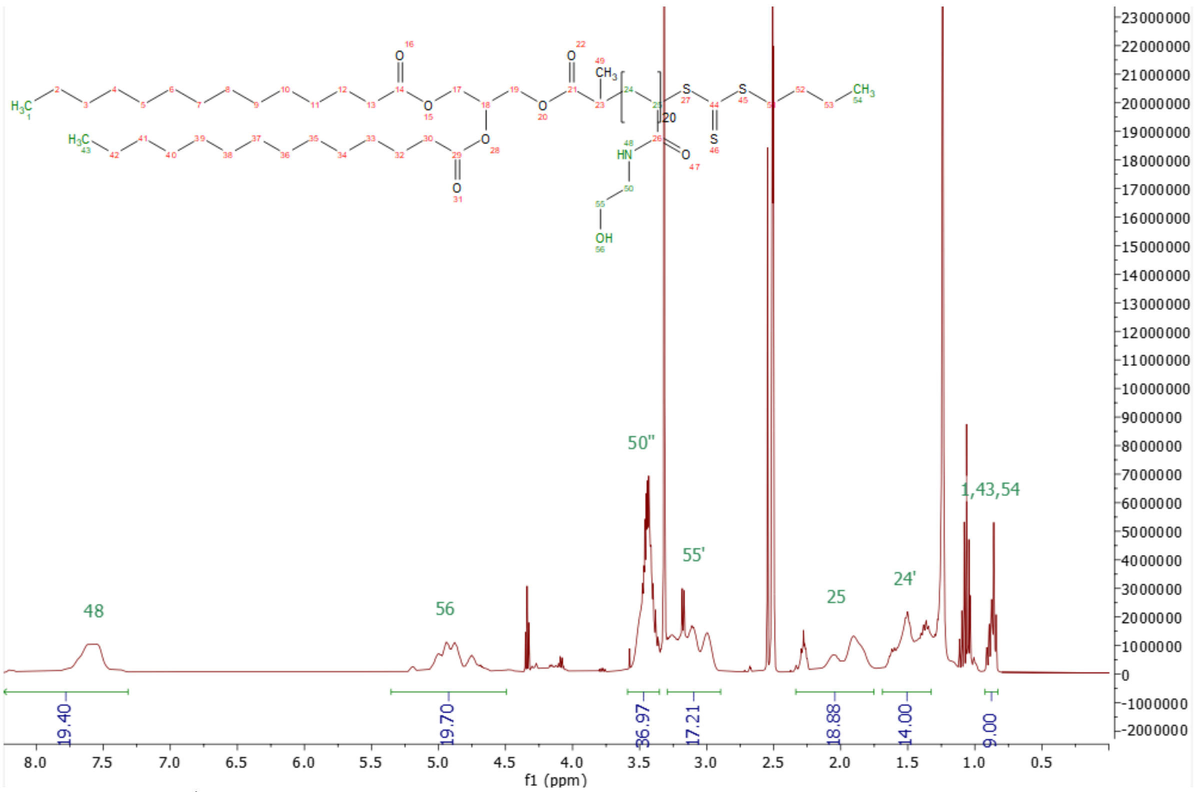

Figure S28 – $^{1}H$ NMR (d6-DMSO, 400 MHz) $HEA_{20}$-S

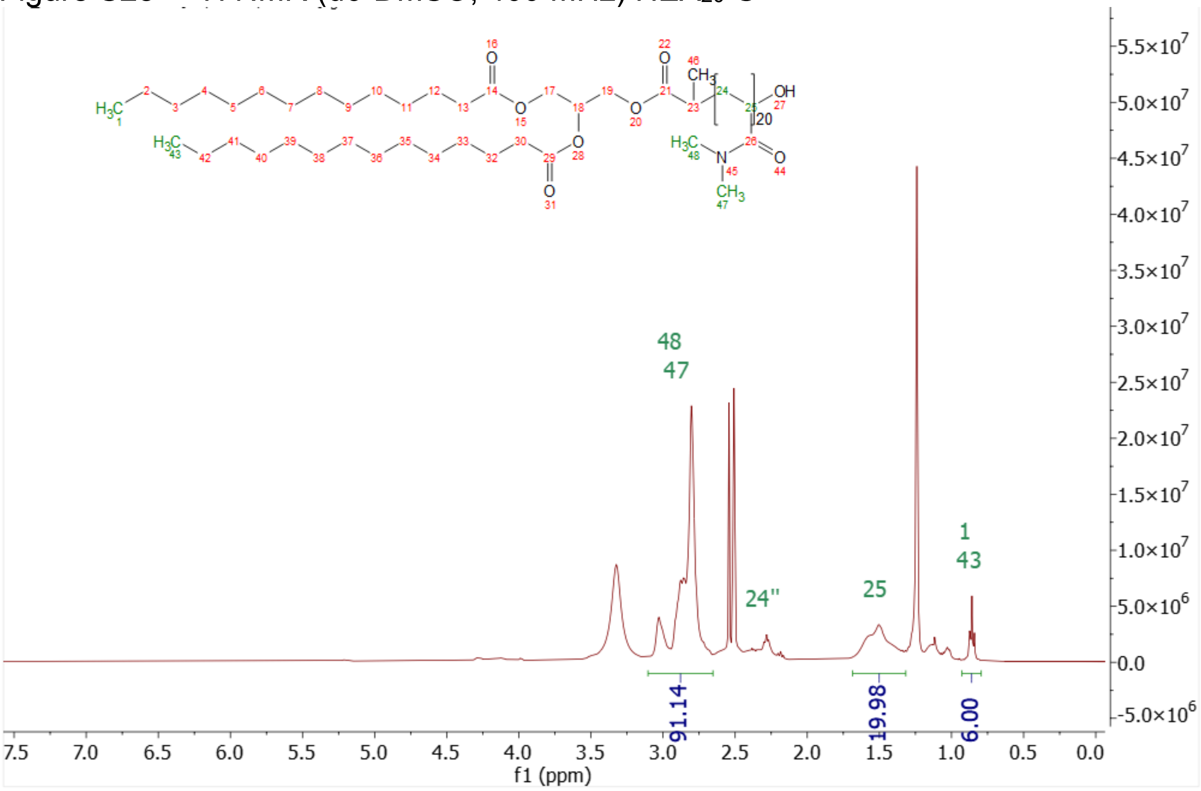

Figure S29 – $^{1}H$ NMR (d6-DMSO, 400 MHz) $DMA_{20}$-OH

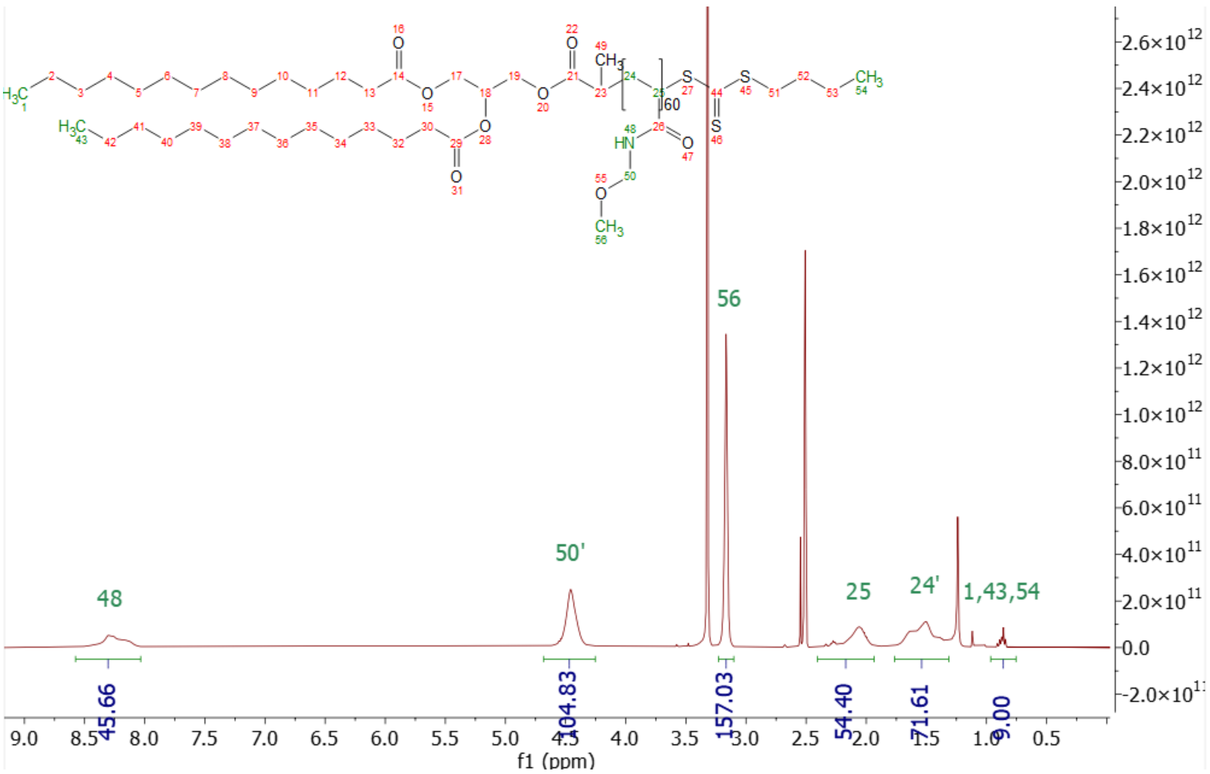

Figure S30 – [1]H NMR (d6-DMSO, 400 MHz) $MOMAM_{60}$-S

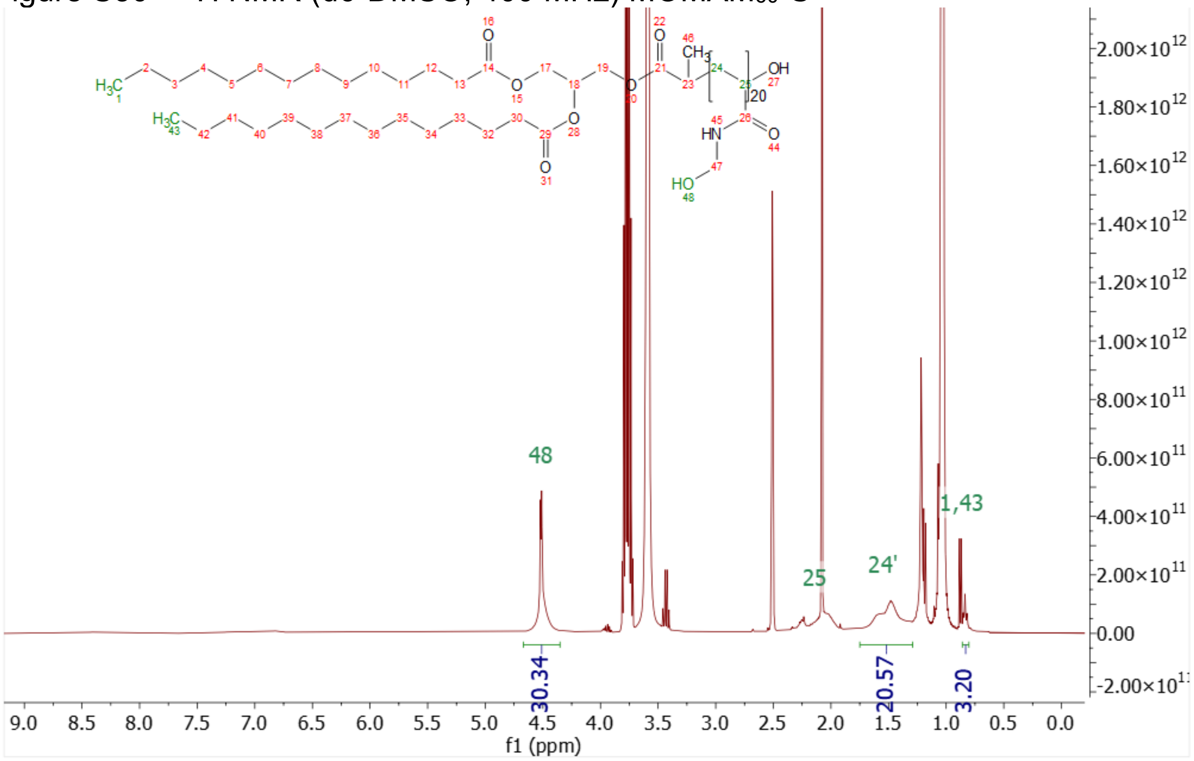

Figure S31 – [1]H NMR (d6-DMSO, 400 MHz) $HMA_{20}$-OH

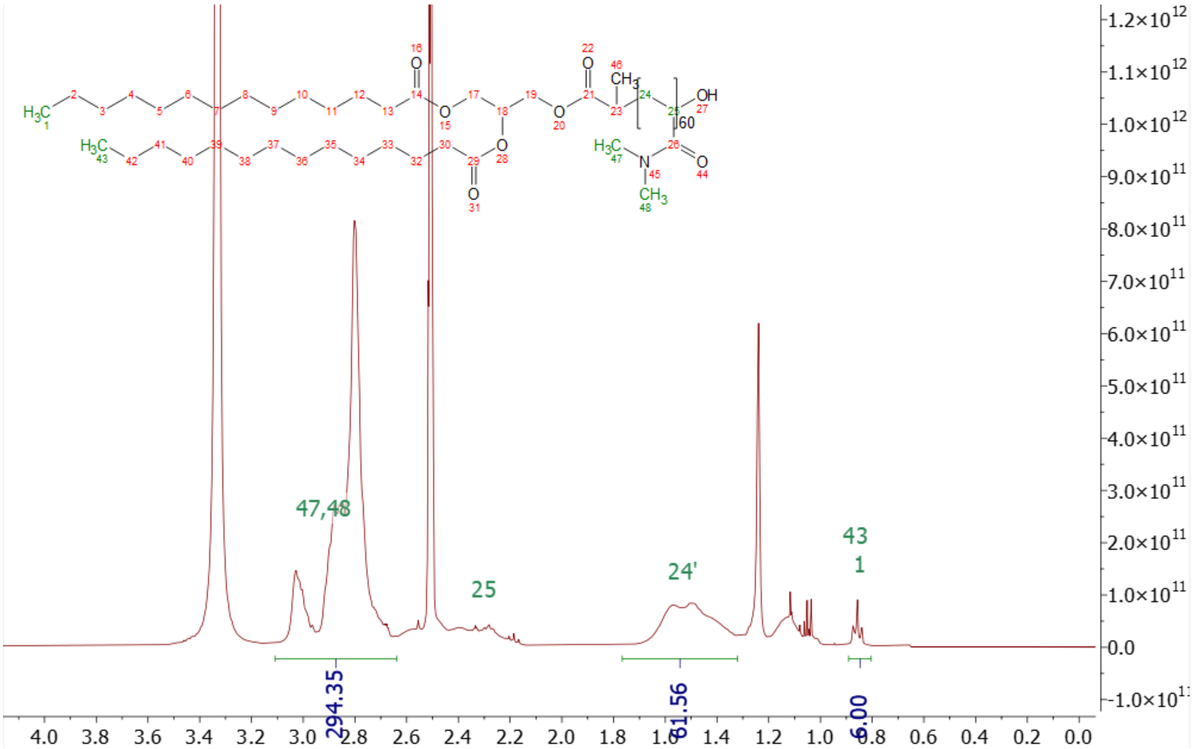

Figure S32 – $^1$H NMR (d6-DMSO, 400 MHz) $DMA_{60}$-OH

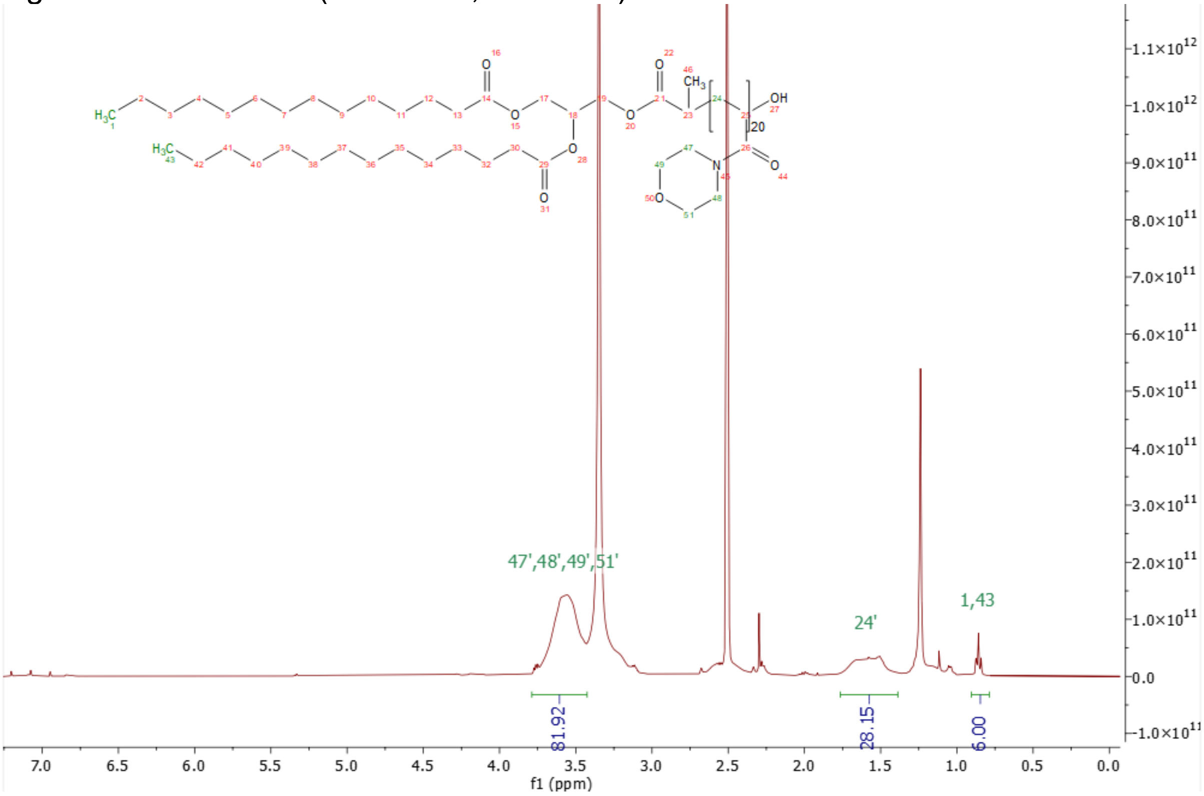

Figure S33 – $^1$H NMR (d6-DMSO, 400 MHz) $NAM_{20}$-OH

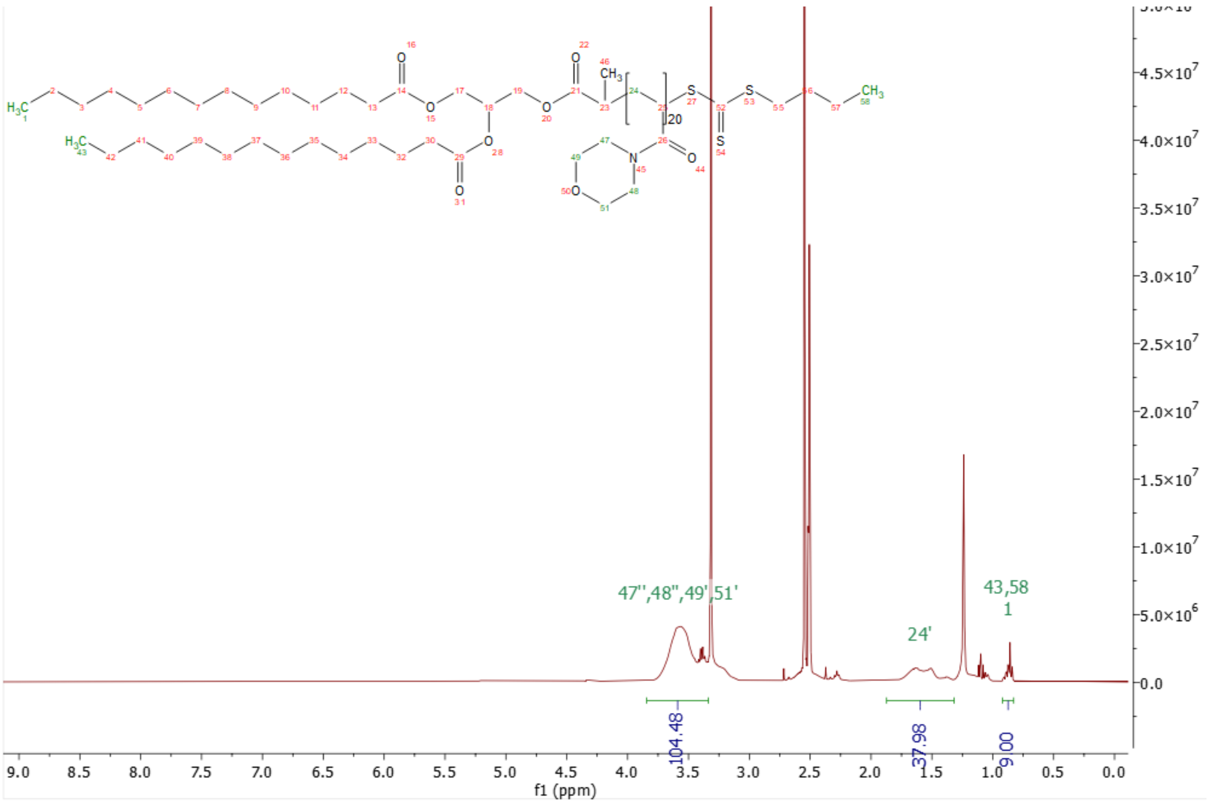

Figure S34 – [1]H NMR (d6-DMSO, 400 MHz) $NAM_{20}$-S

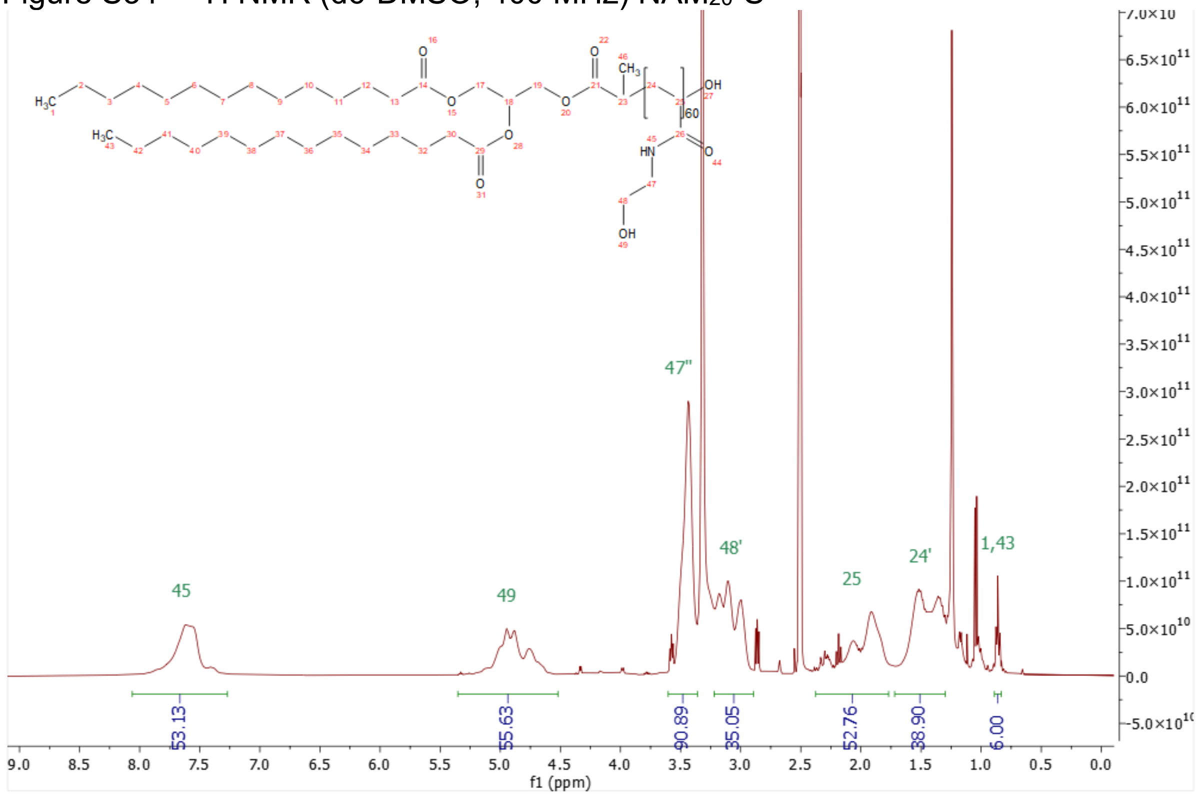

Figure S35 – [1]H NMR (d6-DMSO, 400 MHz) $HEA_{60}$-OH

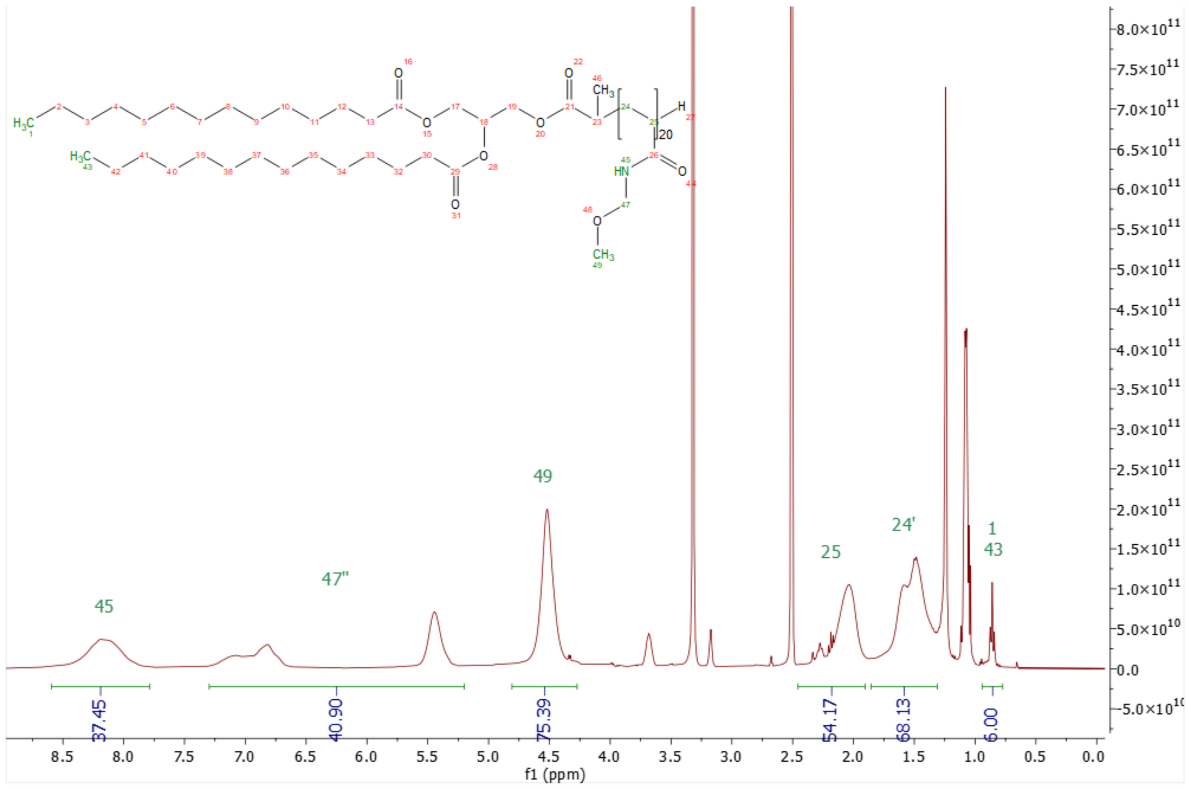

Figure S36 – $^1$H NMR (d6-DMSO, 400 MHz) MOMAM$_{20}$-H
